# Spatiotemporal organisation of residual disease in mouse and human BRCA1-deficient mammary tumours and breast cancer

**DOI:** 10.1101/2025.04.02.646768

**Authors:** Demeter Túrós, Morgane Decollogny, Astrid Chanfon, Myriam Siffert, Lou Romanens, Jean-Christophe Tille, Olivier Tredan, Intidhar Labidi-Galy, Alberto Valdeolivas, Sven Rottenberg

## Abstract

Breast cancer remains one of the prominent causes of death worldwide. Although chemotherapeutic agents often result in substantial reduction of primary or metastatic tumours, remaining drug-tolerant tumour cell populations, known as minimal residual disease (MRD), pose a significant risk of recurrence and therapy resistance. In this study, we describe the spatiotemporal organisation of therapy response and MRD in BRCA1;p53-deficient mouse mammary tumours and human clinical samples using a multimodal approach. By integrating single-cell RNA sequencing (scRNA-seq), spatial transcriptomics (ST), and imaging mass cytometry (IMC) across multiple treatment timepoints, we characterise dynamic interactions between tumour cell subpopulations and their surrounding microenvironment. Our analysis identifies a distinct, drug-tolerant epithelial-mesenchymal transition (EMT) cancer cell population, which exhibits a conserved expression program in human BRCA1-deficient tumours and significantly correlates with adverse clinical outcomes. We further reveal the spatial distribution of residual EMT-like tumour cells within specific anatomical niches, providing a framework for understanding the persistence of MRD and potential therapeutic vulnerabilities. These findings yield a comprehensive molecular roadmap of MRD, opening new avenues for therapeutic strategies targeting EMT-driven drug tolerance and tumour relapse.

## Introduction

Breast cancer accounted for 31% of cancers in females in 2023 [1]. Amongst these, triple-negative breast cancer (TNBC), the most aggressive molecular subtype with the worst prognosis, constitutes 10-15% of all breast cancer cases [2]. TNBC lacks the expression of oestrogen receptor, progesterone receptor, and human epidermal growth factor receptor-2, leading to poor responses to hormone and targeted therapies [2]. Therefore, neoadjuvant chemotherapy is frequently applied as a systemic treatment option. A common combination of chemotherapeutic drugs, called TAC regimen, consists of a taxane (*i*.*e*., docetaxel), a topoisomerase II inhibitor (*i*.*e*., doxorubicin), and an alkylating agent (*i*.*e*., cyclophosphamide). To improve treatment outcomes, other classes of drugs, such as DNA-damaging drugs and immune checkpoint inhibitors, can be added to the regimen. For instance, platinum-based drugs or poly-ADP-ribose polymerase inhibitors, both DNA-damaging agents have been proven very effective in cancer deficient in homologous recombination repair, *e*.*g*., due to BRCA1 or BRCA2 dysfunction. Given the overlap between BRCA1 mutations and the TNBC phenotype [3], the addition of such drugs has increased the pathological complete response (pCR) of non-metastatic TNBC up to 50% [4–6]. Similarly, immune checkpoint inhibitors have been shown to improve pCR [7–9] and survival as well. Despite these advances towards personalised medicine, not all patients without distant metastasis achieve pCR and there is the presence of MRD. Moreover, despite the initial chemotherapy sensitivity, patients with disseminated TNBC are usually not cured and tumours relapse from MRD. Hence, patients with MRD are at higher risk of relapse and death [10] and MRD remains a significant clinical challenge [11].

MRD plays a crucial role in cancer recurrence, and the emergence of pan-resistance from the pool of residual cells highlights the urgent need to better understand the underlying mechanisms of drug tolerance [12–14]. A simple explanation of MRD is the presence of cancer cells in the original tumour that show secondary drug resistance, *e*.*g*., due to random mutations in a pool of heterogeneous cancer cells, which are then selected for during therapy. However, there is circumstantial clinical evidence that relapsing tumours may respond again to the same therapy, and refractory disease only emerges after repeated dosing. Other studies further support the notion that additional mutations causing refractory disease can also emerge during therapy, and that there is a non-mutational biological mechanism that mediates drug tolerance, acting as a transitory phase preceding secondary resistance [15]. In this context, our previous studies using the maximum tolerated dose (MTD) of cisplatin in a genetically engineered mouse model for BRCA1-mutated breast cancer yielded valuable insights into tumour response [16]. Due to a large deletion in the BRCA1 gene, functional restoration of BRCA1 is not possible in this model. Unlike PARP inhibition, which can lead to secondary resistance, the cisplatin MTD induces extensive DNA damage that BRCA1-deficient cells are unable to repair without regaining BRCA1 function, thereby preventing the development of secondary resistance mechanisms. Despite this, tumours persist, eventually regrow, and remain responsive to subsequent cisplatin treatment [16]. This experimental setup ensures that residual tumour cells do not harbour acquired resistance mechanisms, and we can clearly focus on transient drug tolerance as a mechanism of MRD using this model. In line with other studies our previous experiments suggested that transient dormancy allows cells to evade the effects of drugs that primarily target fast-replicating cells [16–19]. Yet, cycling residual cells have also been described [20] causing debate about the relevant mechanisms involved. Moreover, various studies have shown that drug-tolerant cells exhibit distinct transcriptional profiles driven by phenotypic plasticity, notably consisting of an embryonic diapause-like signature [21, 22], epigenetic modifications [23–25], and metabolic shifts [26–29]. Furthermore, EMT phenotypes have shown significant influence on poor prognosis in pan-cancer studies [30].

To get a better understanding of the nature of transient drug tolerance as mechanism of MRD, we combined scRNA-seq, ST, and IMC in our BRCA1;p53-deficient mouse mammary tumours. Using these technologies, we uncovered the spatiotemporal dynamics of several distinct tumour cell types and cellular niches and described their interplay with the tumour microenvironment (TME). We described the functional characteristics of these molecular niches over spatial and temporal scales and directly examined their relationship with tissue-bound platinum-based chemotherapy agents by integrating these modalities. Moreover, we identified a specific tumour cell niche and its associated gene expression program consistent with quiescence and EMT phenotypes, providing a prominent survival advantage for these cells regardless of the treatment modality. Notably, while the residual tumour islands are mainly composed of the EMT niche, the composition of recurrent tumours is identical to those of the primary tumours, highlighting the dynamic and reversible aspect of MRD. Furthermore, our findings were translatable to human BRCA1-deficient breast cancers, where the expression program of the EMT niche is highly specific to the residual tumour islands. Stratifying TNBC patients based on expression levels of the EMT program further showed a significant reduction in survival outcomes, highlighting the clinical relevance of our findings. Beyond temporal profiling of MRD, we also examined the spatial distribution of the drug-tolerant cells and their possible interactions with the TME. These findings unveil key components to further understand MRD and may support the development of new therapeutic strategies to eradicate residual cancer cells.

## Results

### Spatiotemporal multi-omics profiling of MRD

To chart intratumoural heterogeneity and understand the key drivers of residual disease, we first collected primary (treatment-naïve), residual, and recurrent K14*cre*;*Brca1*^F/F^;*Trp53*^F/F^ (KB1P) mouse mammary tumours following chemotherapy. We previously demonstrated that the MTD of DNA-crosslinking therapies does not lead to acquired secondary resistance, unlike the PARP inhibitor olaparib [16, 31, 32]. Hence, this model allows us to focus on transient mechanisms of drug tolerance when treating the KB1P tumours with chemotherapeutic agents, avoiding the presence of stable, secondary drug resistance mechanisms in the residual tumours. Additionally, contrary to the traditional xenograft model, this approach enables the investigation of residual disease in an immunocompetent mouse model that highly resembles the human disease [33].

#### Single-cell sequencing reveals altered cell type composition in MRD

Using this model, we performed scRNA-seq from the primary and residual tumours (*n*=6), applied ST using the 10x Visium platform from the primary, residual, and recurrent tumour tissue sections (*n*=30), and conducted IMC on primary tumours immediately following chemotherapy treatment (4h-24h), as well as on residual tumours (*n*=7) (**Fig.1a**). To minimise the effects of individual genetic variability, all tumour samples were derived from two parental tumours that were subdivided and transplanted into the fourth mammary fat pad of syngeneic FVB/NJ mice. Tumour growth was closely monitored (**Supplementary Fig.1**) and the primary tumours (*n*_ST_=10, *n*_scRNA-seq_=3) were excised after reaching the size of 8 × 6 mm. To induce MRD, we treated animals with two chemotherapy regimens employed in clinical practice to target HR-deficient tumours by inducing DNA damage: cisplatin and TAC regimen. Subsequently, we extracted and sectioned the residual tumours after 7 (TAC: *n*_ST_=2, cisplatin: *n*_ST_=2) or 12 days (TAC: *n*_ST_=4, *n*_scRNA-seq_=3, cisplatin: *n*_ST_=3) corresponding to the minimal tumour size. Recurrent tumour samples (TAC: *n*_ST_=2, cisplatin: *n*_ST_=2) were derived after tumours again reached the criteria size following chemotherapy treatment (**Fig.1b**). Furthermore, we sampled tumours within the first 24 hours after chemotherapy to examine cisplatin drug uptake and its relationship with tumour heterogeneity (*n*_ST_=5, *n*_IMC-ROI_=53). This experimental design enabled us to acquire a detailed spatiotemporal map of residual disease progression, capturing cellular plasticity at the single-cell level across multiple pivotal time points during MRD progression, alongside spatial heterogeneity and tissue architecture. Additionally, we acquired primary and residual tumour tissue samples from human BRCA1-deficient breast cancers and performed ST (*n*_Primary_=2, *n*_Residual_=3).

**Fig. 1.**
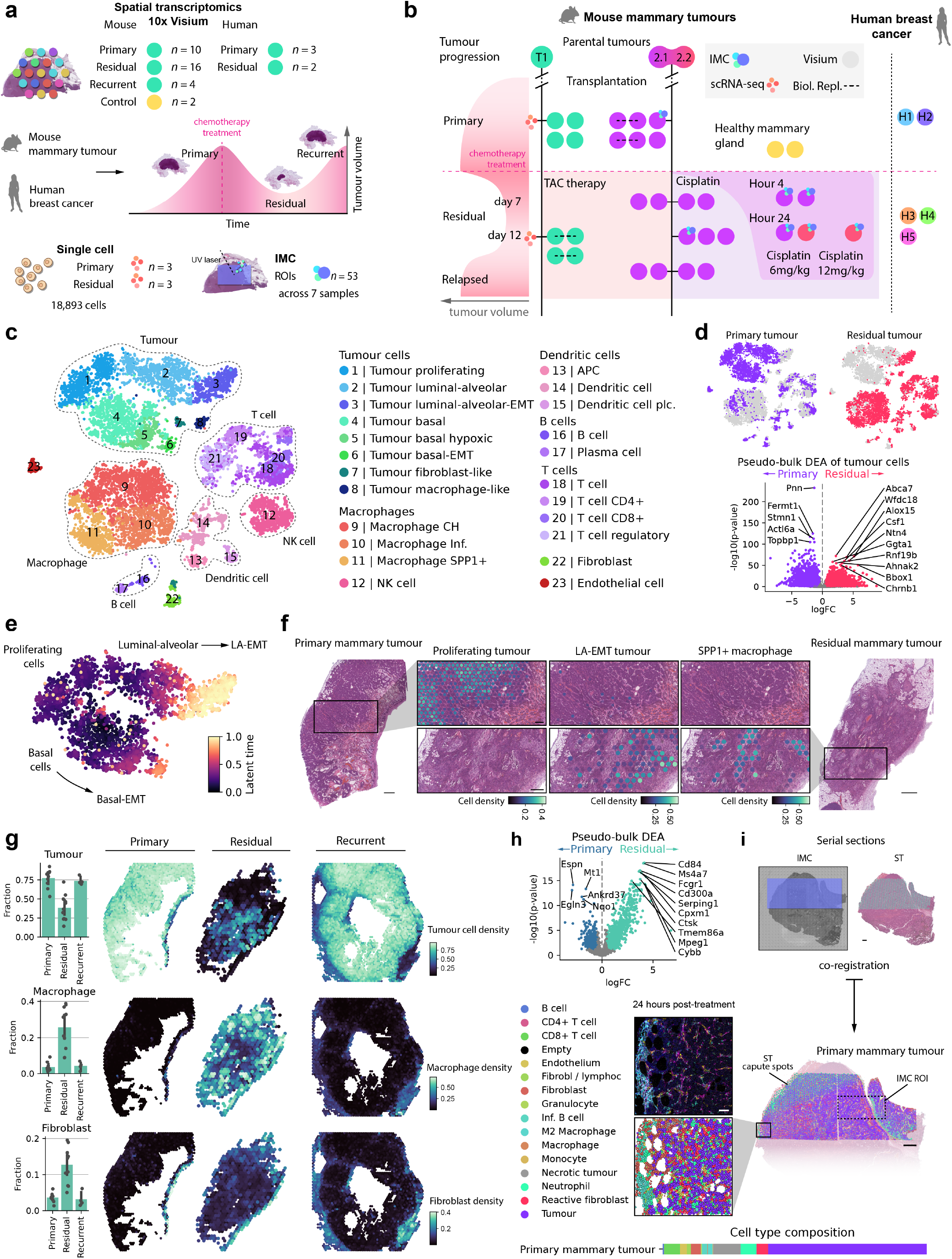
Single-cell and spatial transcriptomic profiling of MRD. **a**, Mouse mammary tumour and human breast cancer samples were collected at different stages of tumour development, remission, and recurrence before subjecting them to scRNA-seq, ST, and IMC. **b**, Mouse mammary tumours were derived and transplanted to immunocompetent animals from two parental tumours (T1, T2.1-2.2). Tumour samples and healthy mammary tissue were collected from the primary tumours, residual, and recurrent tumours. The graph indicates the sampling time points for the different modalities and treatment types. **c**, T-distributed stochastic neighbour embedding (t-SNE) representation of single cells collected from primary and residual tumours showing the major cell types and subpopulations. **d**, Cell type composition of primary and residual tumours highlighted on the t-SNE plots (upper panel) and the corresponding pseudo-bulk DGEA results of tumour cells (lower panel). **e**, RNA velocity analysis of tumour cells indicates cell transitions from proliferating to luminal-alveolar or basal cancer cell phenotypes. **f**, H&E sections of primary and residual tumour samples overlaid with cell type deconvolution results of three cell types demonstrating distinct spatial and temporal patterning during tumour progression (scale bars: 500 µm, inset scale bars: 200 µm). **g**, Cellular composition of primary, residual, and recurrent tumour samples. Barplots showing the mean cell fraction across all samples (black dots: individual ST samples, error bar: 95% CI, bar height: mean value). **h**, ST pseudo-bulk DGEA of primary and residual tumours showing top differentially expressed genes before and after chemotherapy. **i**, Construction of multimodal datasets. Serial sections of mammary tumours were first subjected to IMC and ST, followed by co-registration based on tissue morphology to align the two modalities. Using the IMC data, we annotated the tissues on the single cell level, which were coupled with transcriptional information (scale bars: 500 µm, inset scale bar: 50 µm).

We first established the cellular composition of the mammary tumours using scRNA-seq. We identified 23 distinct cell populations across 8 major cell types through unsupervised clustering of the integrated scRNA-seq data during tumour progression and MRD (**Fig.1c**). We identified a substantial tumour cell cluster (*Krt14, Krt18*) alongside several cell types from the TME, such as macrophages (*Cd68*), T cells (*Cd3*), B cells (*Cd19*), dendritic cells (*Lsp1, Flt3*), natural killer (NK) cells (*Ncr1, Klrb1c*), endothelial cells (*Pecam1*) and fibroblasts (*Fn1*).

Examination of the tumour cell cluster revealed a highly proliferative tumour cell population characterised by high expression of *Mki67, Cdk1*, and several other cell cycle and cell division genes, branching into two separate lineages; basal (*Krt17, Krt14, Krt5, Acta2*) and luminal-alveolar (*Krt19, Krt18*) tumour cells, consistent with previous findings [34, 35]. Within the main basal cell cluster, we identified a basal hypoxic cell group distinguished by the expression of hypoxia-induced genes, such as *Egln1* and *Vegfa*. In both lineages, we uncovered distinct tumour cell populations displaying EMT signature, marked by the expression of several EMT marker genes (*e*.*g*., *Clu, Crip1, Lgals3*). We further characterised two small tumour subpopulations: macrophage-like tumour cells, which express genes typically associated with hematopoietic cells (*Ctss, Cd74*), mainly macrophages, and fibroblast-like tumour cells, which share an expression signature with fibroblasts (*Zeb2, Arpc1b, Vim, Mgp*).

Within the cell types of the TME, we have further identified Cd4+, Cd8+, and regulatory T cells, naive, activated (*RelB*+), and immunosuppressive (*Siglech*+, *Ccr9*+) dendritic cells, and plasma cells. Additionally, we have identified multiple tumour-associated macrophage subtypes, such as opsonin-induced (*C1qa, C1qc, C1qb*) M2 macrophages, M1 macrophages, and *Spp1*+ macrophages which have been previously linked to ECM remodelling and immune evasion in other cancer types [36].

Examination of the differences between major cell type populations across the primary and residual states revealed a substantial enrichment of immune cells in the TME, notably a significant increase in the M2, *Spp1*+ macrophage, T cell, and NK cell populations, as well as the EMT tumour cell clusters (**Fig.1d**). Differential gene expression analysis (DGEA) (**Supplementary Table 1**) revealed, that these residual tumour cells share upregulated genes, such as *Abca7* (ABC transporter, expressed across various cancer types, involved in lipid transfers and mesenchymal transformation [37]), *Wfdc18* (associated with luminal cancer cell phenotype and stemness [38]), *Alox15* (dioxygenase, suppresses immune response in colorectal cancer [39]), *Csf1* (hemocine, usually expressed by macrophages, promotes M2 phenotype and leads to worse prognosis in breast cancer [40]), *Ntn4* (laminin-like secreted protein, secreted by breast epithelial cells and sequestered by the basement membrane [41]), *Ggta1* (alpha-1,3-galactosyltransferase), *Rnf19b* (E3 ubiquitin-protein ligase), *Ahnak2* (key regulatory role in tumour progression by activating signalling pathways such as ERK, MAPK, WNT, MEK, and promoting EMT [42]), *Bbox1* (essential gene for TNBC tumourigenesis, depletion inhibits TNBC cell growth [43]), *Chrnb1* (acetylcholine receptor subunit beta).

Top downregulated genes include *Pnn* (increased proliferation in cancer [44]), *Fermt1* (knockdown of Fermt1 significantly decreased cell proliferation by mediating EMT and cell cycle arrest in nasopharyngeal carcinoma [45]), *Stmn1* (promotes microtubule polymerisation and invasion, anti-apoptosis, and cell proliferation [46]), *Actl6a* (related to the tumourigenesis of several cancers [47]), and *Topbp1* (*BRCA1/2* and HR-deficient tumours have been reported to adapt to proliferate through CIP2A-TOPBP1 complex [48]). RNA velocity analysis of the tumour cell clusters further revealed cellular plasticity and differentiation trajectories, highlighting the transition from the proliferating tumour cell state to the luminal-alveolar or basal EMT clusters (**Fig.1e**). These state transitions imply dynamic reorganisation across tumour cell populations during disease progression. To examine this *in situ*, we next focused on mapping these cells within the tissue, allowing us to explore their spatial organisation and how these transitions unfold in the TME.

#### Cell type deconvolution identifies spatial heterogeneity in MRD

To gain a deeper understanding of the spatiotemporal organisation of these cell types in MRD, we assessed the ST data alongside the corresponding histology. Cell type deconvolution with cell2location [49] using our single-cell reference and histopathological analysis of the primary tumours revealed that cancerous cells grow in solid nests embedded in a dense fibrous stroma (**Fig.1f**).

In these primary tumours, we observed an even and extensive distribution of proliferating, basal, and luminal-alveolar tumour cells, while the EMT phenotypes were confined to smaller islands within the tumour mass. Proliferating tumour cells were densely packed, polygonal, and pleomorphic. Mitotic figures were common, some of which were atypical [50], indicative of aggressive proliferation. A mixed immune cell infiltration was mildly present, particularly at the tumour periphery (**Supplementary Fig.2a**).

In the residual tumours, the architecture and growth pattern changed drastically. The tumour cells became elongated and difficult to delineate within the highly cellular and reactive stroma (**Supplementary Fig.2b-c**). Mitotic figures were rare or remained elusive corroborated by the absenceof proliferating tumour cells and the enrichment of EMT cells. Immune cell infiltration was predominantly mononuclear, and nests of foamy macrophages were scattered within the tumour, forming specific anatomical niches occupied by *Spp1+* macrophages (**Supplementary Fig.2d**). In the recurrent stage, however, tumour architecture and cell morphology resembled those of the primary tumours once again.

The tumours also displayed large necrotic areas consistent with the morphological characteristics of BRCA1-mutated TNBC [33] (**Supplementary Fig.3**). Hypoxic tumour cell populations were predominantly located in close proximity to these necrotic cores.

By examining the overall changes in cell type composition in the tumour containing capture spots (**Supplementary Fig.4**), we observed an up to 85% reduction in tumour cell fractions (**Fig.1g**). This decrease was predominantly attributed to the reduction of proliferating, basal, and luminal-alveolar tumour cells, while the fraction of basal and luminal-alveolar EMT phenotypes, along with the macrophage-like and fibroblast-like tumour cell types showed enrichment after treatment (**Supplementary Fig.4b**). Simultaneously, the TME underwent significant remodelling with a considerable increase in immune cells, especially macrophages, and fibroblasts (**Supplementary Fig.4c**). Consistent with histopathology, the cellular composition of the recurrent tumours mirrors a state similar to that of the primary tumours.

Pseudo-bulk DGEA of tumour-containing capture spots between primary and residual tumours (**Supplementary Table 2**) showed genes driving tumour immune response related to innate immunity, mast cells, monocytes, and the complement system, such as *Cd68, Ms4a7, Fcgr1, Cd300a, Serping1, Cpxm1, Mpeg1, Cybb* and cancer proliferation-inducing genes and ECM remodelling genes, such as *Ctsk [51]* in the residual state (**Fig.1h**). Amongst the upregulated genes in primary tumours, we found genes such as *Espn* (F-actin cytoskeleton remodelling, growth and metastatic regulator in melanoma [52]), *Egln3* (downregulation increases gastric cancer progression and EMT [53]), *Mt1* (playing key roles in tumour growth, angiogenesis, immunomodulation and differentiation [54]), *Nqo1* (regulates oxidative stress, overexpressed in many cancer types [55]).

To better understand the spatial organisation of mammary tumours after chemotherapy treatment at the single cell level, we processed the IMC data by segmenting and phenotyping individual cells (**Fig.1i and Supplementary Fig.5**). Leveraging serial tissue sections, we co-registered IMC-Visium pairs based on the H&E and IMC panorama images, enabling the mapping of transcriptomic features to individual cell groups. Using this approach, we assessed the cell type composition of each tumour and examined the tissue at single-cell resolution. We identified solid tumour islands, infiltrated by various stromal and immune cells, such as fibroblasts and M2 macrophages.

In summary, by combining scRNA-seq, ST, and IMC, we mapped the spatiotemporal progression of residual disease in BRCA1;p53-deficient mammary tumours. We characterised the different cell types present in the tumours and further defined tumour subpopulations with different sensitivity to chemotherapy. This allowed us to get a deeper understanding of the underlying mechanisms responsible for those different sensitivities in their spatial context.

### Chemotherapy-induced changes in the transcriptional tissue landscape of MRD

To uncover the molecular changes during MRD and recurrence in an unbiased manner, we inferred cellular niches (*i*.*e*., tissue compartments) relying on ST data. First, we integrated and projected expression signatures into a shared embedding and leveraged our recently published machine learning-based approach, Chrysalis [56], to identify spatially and functionally distinct molecular tissue compartments in ST. This allowed us to interrogate the tumour niche composition in space and time and examine the corresponding gene expression programs. Moreover, it provided insight into how surviving cells are organised into different niches that may facilitate MRD.

Characterisation of these tissue compartments revealed five tumour cell-associated (0, 2, 5, 8, and 10) and six TME-associated (1, 4, 7, 9, 11, 12) compartments (see **Methods**) with specific spatial localisation (**Fig.2a and Supplementary Fig.6**) and gene expression programs (**Supplementary Fig.7a and Supplementary Table 3**). We identified a proliferating tumour niche (5) characterised by genes associated with cell cycle regulation and proliferation, such as genes encoding for histones (**Fig.2b**) or *Ncl* (nucleolin, chromatin decondensation), *Ccne1* (Cyclin E1), *Pclaf* (binds to PCNA, which is a cofactor for DNA polymerases), *Dbf4* (cell cycle regulation), *Pprc1* (promotes mitochondrial biogenesis), *Dhfr* (key role in thymine synthesis) and correlated with large proportions of the primary tumour mass. The hypoxic tumour niche (10) was linked to necrotic cell bodies surrounding necrotic cores, and expressed genes such as metallothioneins, *Egln1* (hypoxia-inducible factor prolyl-hydroxylase), *Slc2a3* (glucose transporter), *Pgf* (angiogenesis), *Vegfa* (angiogenesis), *Gys1* (glycogen synthase). We further identified an EMT niche (2) characterised by the presence of *Anxa1* (a wellknown immunomodulator in breast cancer [57]) and members of the *Wfdc* protease inhibitor gene family like *Slpi*, which are strongly correlated with increased metastatic potential in multiple cancer types [58–61]. Furthermore, this niche was characterised by the expression of *Lypd3* (laminin-binding protein supporting cell migration), *Ppl* (desmosome component), *Kcnq1* (potassium transport channel), *Calml3* (calmodulin, key role in cell adhesion and motility in many cancer types [62]), and *Csn3* (casein kappa, transport of inorganic ions and amino acids) among others. Compartments 0 and 8 were less ubiquitous and were localised to certain samples, representing a subset of proliferating compartments and a niche eliminated by chemotherapy, respectively.

**Fig. 2.**
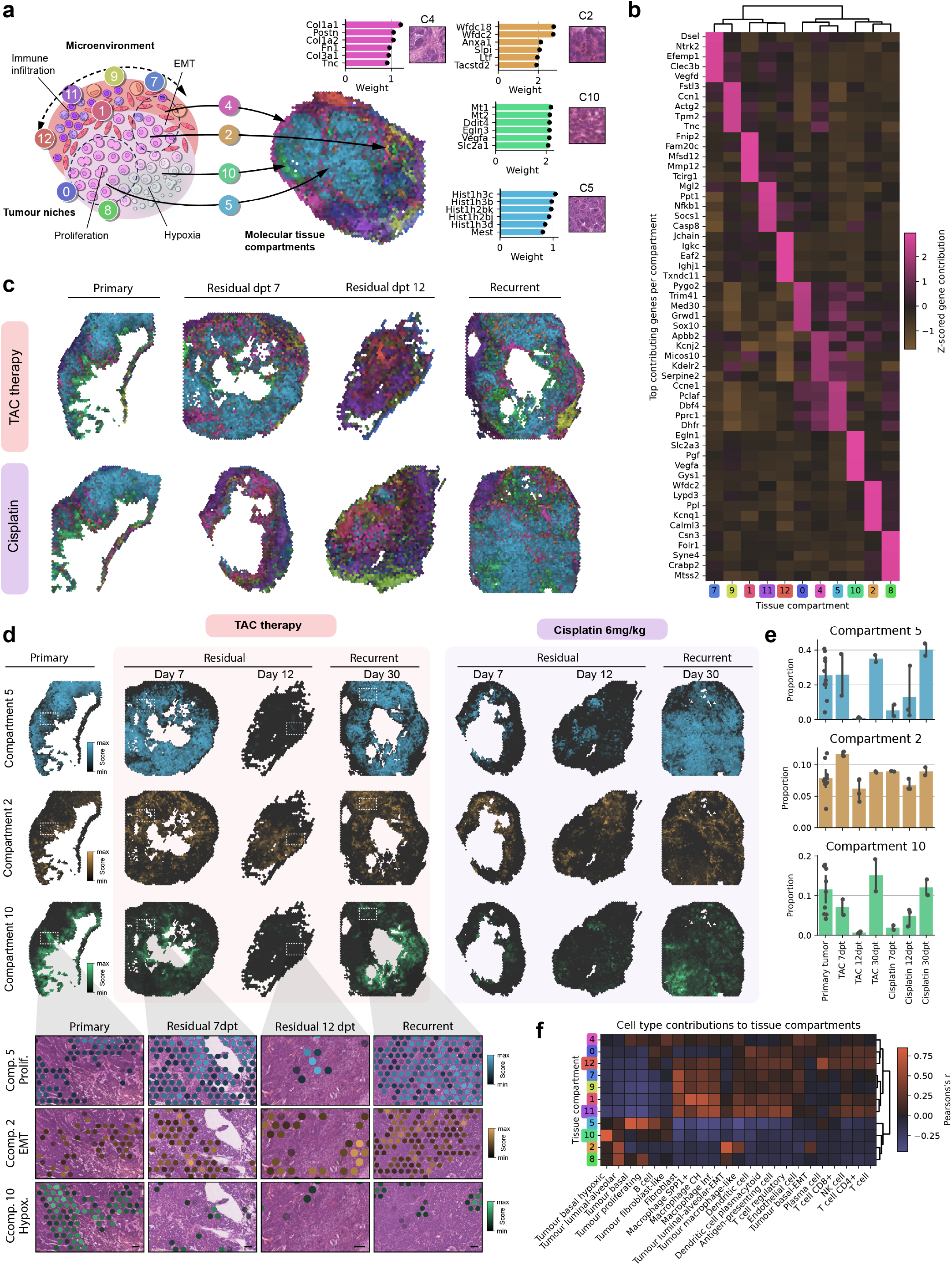
Spatiotemporal dynamics of cellular niches during tumour progression. **a**, Schematic representation (left) of the different molecular tissue compartments identified by Chrysalis and their MIP on one representative primary tumour tissue section (right). For niches 4, 2, 10, and 5, five genes with the highest weights from their corresponding gene expression program are shown alongside the histology image tile of these niches. **b**, Heatmap showing the contributions of top genes to the identified tissue compartments. **c**, MIP of the identified tissue compartments on representative tumour tissue sections of primary, residual (7 and 12 dpt), and recurrent tumours (∼30 dpt) after TAC or cisplatin treatment. Projections show substantial spatiotemporal reorganisation after chemotherapy and during tumour recurrence. **d**, Spatial plots of compartment scores of proliferating (5), EMT (2), and hypoxic (10) niches on representative tumour tissue sections of primary, residual and recurrent tumours after TAC or cisplatin treatment. Zoom-in image tiles with histology overlayed show the fine-scale spatial distribution of these tumour niches (scale bar: 100 µm). **e**, Proportions of the proliferating (5), EMT (2), and hypoxic (10) niches across all tumour samples. While proliferative and hypoxic compartments are greatly affected by treatment, the EMT compartment fraction remains constant (bar height: mean compartment fraction, black dots: individual ST samples, error bar: 95% CI, bar height: mean value). **f**, Heatmap displaying the correlation between cell type and tissue compartment fractions across all tumour samples.

We revealed TME-associated niches, ranging from fibrotic stroma to increasing degrees of immune infiltration (from left to right: 4, 9, 7, 1, 11, 12; **Fig.2a**). Compartment 4 was located in close proximity to tumour cells and showed genes expressed mainly by fibroblasts, such as collagens, and other extracellular matrix components: *Tnc* (tenascin), *Postn* (periostin), *Fn1* (fibronectin). Compartment 9 exhibited an expression program composed of collagens and smooth muscle cell-related genes, whereas Compartment 7 described a niche with light immune infiltration, as exemplified by genes related to the complement system (*e*.*g*., *Cb4*) and macrophages (*e*.*g*., *Retnla*). Compartment 1 was dominated by genes such as cathepsins and *Cd68*, indicating monocytic cell populations, which is also evident in niche-associated histology. Compartment 11 associated genes, for example, *Retnla, Cd74, C3*, and *Ccl8* indicated lymphocyte infiltration and activated immune response. The expression program of Compartment 12 involved genes, such as *Igkc, Jchain, Ighj1*, and other immunoglobulin genes indicating B cell activation and humoral immune response. This compartment coincided with the localisation of plasmacytic cells in the histology.

To gain a deeper understanding of the cellular evolution of the uncovered cellular niches during tumour growth, MRD, and relapse, we investigated the four different time points: primary tumour, MRD at 7 and 12 days post-treatment (dpt), and recurrent tumour at ∼30 dpt, in both TAC- and cisplatin-treated samples. Inspecting the maximum intensity projection (MIP)-based visualisation of the compartments revealed substantial changes in cellular niche composition during MRD (**Fig.2c and Supplementary Fig.7b**). Primary tumours were predominantly composed of the proliferating (5) tumour niche that was surrounded by fibrotic stroma (4) and other immune cell-dominated compartments. Surrounding the necrotic cores, the hypoxic tumour niche (10) was dominant, while the identified EMT niche was distributed throughout the tumour without a discernible pattern of localisation. In the residual tumours, we observed substantial remodelling: the majority of tumour tissue is taken over by compartments such as 11 and 12, indicating significant immune cell infiltration and phagocytic activity by macrophages. Examining the recurrent tumours, the tissue composition resembles the original state observed before chemotherapy.

Next, we focused on characterising the temporal dynamics of the proliferating (0), hypoxic (10) and EMT (2) niches (**Fig.2d**). Proliferating and hypoxic tumour cell niches diminished significantly after chemotherapy in both treatment groups, while the EMT (2) compartment remained largely unchanged. Quantifying tumour niche coverage, we confirmed the complete depletion of the proliferating and hypoxic compartments in MRD (**Fig.2e**). Yet, no significant change in the EMT compartment was detected, highlighting the drug-tolerant nature and stability of this tumour niche and its associated cell types. We also uncovered drug-specific changes, influencing the kinetics of tumour reorganisation. In the TAC-treated samples the maximum effect was detected at day 12 post-treatment, whereas in the cisplatin-treated tumours, this was observed earlier, on day 7. These trends were also apparent in the TME-associated niches (**Supplementary Fig.8-9**). Amongst these, the plasma cell-driven cellular niche was enriched specifically in the TAC-treated MRD samples. Strikingly, the spatial composition in the recurrent tumours closely resembled that of the primary tumours.

We further investigated the cellular composition of the identified niches (**Fig.2f**) by calculating the correlation with the cell type deconvolution results. Tumour niches correlated specifically with identified tumour cell types. Proliferating(5) niche could be assigned to proliferating and basal tumour cells, while the hypoxic tumour cells were exclusively mapped to the hypoxic compartment (10). The EMT niche was highly correlated with the luminal-alveolar-EMT cell population. Immune cell types in the TME could be assigned to the TME niches (9, 7, 1, 11, 12) with substantial enrichment in macrophages in compartments 1 and 11, and plasma cells for Compartment 12. Some compartments could not be attributed to cell types due to unmapped cell states or cells not recovered in our single-cell reference.

These results showed reversible dynamic reorganisation and interplay between distinct cellular niches in MRD. Despite this remodelling, the abundance of the EMT tumour niche (2) remained unaffected by chemotherapy.

### Functional characterisation of MRD

We then sought to characterise the spatiotemporal changes in functional treatment response and how the identified cellular niches can contribute to these functional characteristics. For this purpose, we started by calculating pathway activity scores for signalling pathways involved in cancer using PROGENy [63] (**Fig.3a, Supplementary Fig.10**). In the primary tumours, we have found strong downregulation of the Trail apoptotic pathway, that only diminished during the time points of maximum response and is further downregulated in the recurrent tumours. When inhibited, the Trail apoptotic pathway has been shown to play an important role in tumour onset and progression [64]. Similar trends were observed for p53, EGFR, TGFb, and JAK-STAT pathways. Interestingly, JAK-STAT upregulation has been associated with early-onset cancer progression, however we only measured activation during MRD [65]. On the contrary, we observed significant down-regulation in MAPK pathway activity during MRD, which might be the result of the diminishing proliferating tumour cell population. Studying the pathway correlations with the cellular niches, we found, that the TME-associated niches correlate with increased NFkB, Trail, p53, TGFb, EGFR, JAK-STAT, and TNFa pathways activities (**Fig.3b**). Proliferating tumour niche (5) specifically correlated with MAPK, PI3K, and Estrogen pathways, while the hypoxic niche (10) aligns with elevated hypoxia activity. Investigation of the spatial pathway activity maps further corroborated the TME-associated JAK-STAT pathway enrichment and the association of MAPK activation and Trail inhibition to the proliferating tumour mass (**Fig.3c**).

**Fig. 3.**
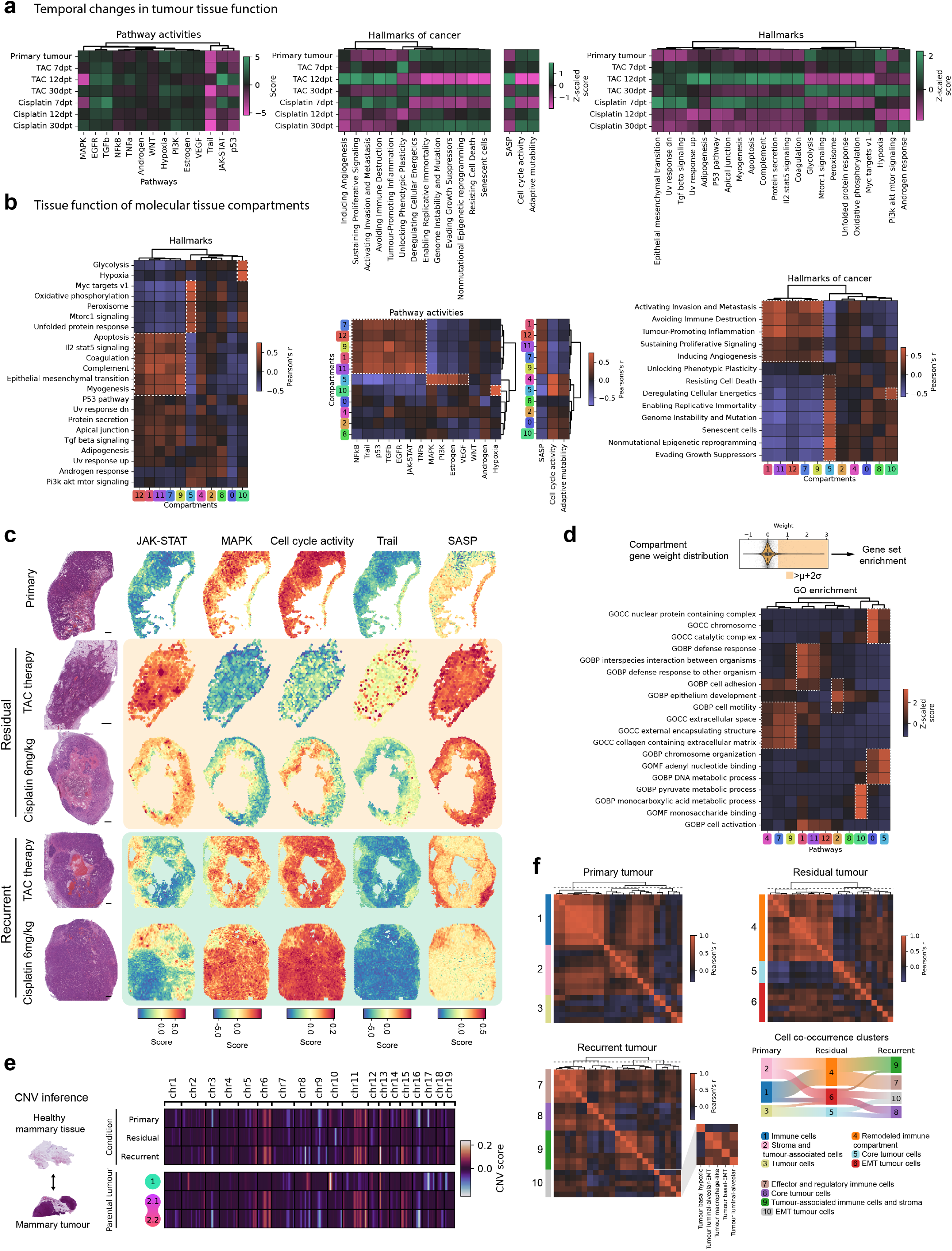
Functional dissection of tumour dynamics across spatial and temporal scales. **a**, Heatmaps illustrating changes in intracellular pathway activities, significantly enriched gene sets from the Hallmarks collection, Hallmarks of Cancer, and other gene set scores across various time points, from primary tumours to MRD and recurrence. **b**, Heatmaps showing correlations of cellular niches with intracellular pathway activities, significantly enriched Hallmarks, and Hallmarks of Cancer gene sets. Terms highlighted in the main text are marked with white dashed-line rectangles. **c**, Spatial plots of selected gene set and pathway activity scores showing distinct spatiotemporal organisation shown for representative samples, alongside the H&E staining (scale bar: 500 µm). **d**, GO enrichment heatmap of gene expression programs associated with each cellular niche. Genes for each compartment were selected if their contribution to cellular niches exceeded the mean plus two standard deviations (µ+2σ). Terms highlighted in the main text are marked with white dashed-line rectangles. **e**, CNV inference was conducted by contrasting mammary tumour samples with healthy mammary tissue samples. Heatmaps demonstrating chromosomal signatures ordered according to the different time points and different parental tumours. Inferred CNV alterations highly correlate with parental tumours. **f**, Heatmaps show cell co-localisation clusters determined based on dendrogram distance cutoff from root (3.5, dashed line). Sankey diagram demonstrates the reorganisation of clusters during tumour progression.

We followed the characterisation using the Hallmarks of Cancer gene sets from a recent study [66], and revealed cancer hallmarks activated during MRD, such as *tumour-promoting inflammation* and *avoiding immune destruction* (**Fig.3a and Supplementary Fig.11**). These cancer hallmarks weree strongly correlated with TME compartments (**Fig.3b**). In the proliferating tumour niche, we found increased expression of gene sets associated with *evading growth suppressors* and *enabling replicative immortality*, while in the hypoxic niche, a strong correlation was observed with *deregulating cellular energetics*. These expression programs were downregulated during MRD as these tumour cell populations receded in response to chemotherapy.

Since drug-tolerant residual cells have been shown to exhibit a senescence-like phenotype [67, 68], we sought to investigate the senescence-associated secretory phenotype (SASP) within MRD and the identified cellular niches. Our analysis revealed significant upregulation of the SASP signature [69] in MRD compared to the primary and recurrent tumour stages (**Fig.3a and Supplementary Fig.12**). Correlating this signature score with the tissue compartments showed a strong anti-correlation with the proliferating tumour niche and strong correlations with the TME-associated immune cell compartments. However, no significant correlation was observed with the EMT, or the hypoxic tumour niches (**Fig.3b**). At the capture spot level (**Fig.3c**), elevated SASP levels were predominantly associated with TME, particularly with the fibrotic tissue structures surrounding the tumour.

We also calculated the cell cycle activity based on the upregulation of different cell cycle phase marker genes, since quiescence and cell cycle alterations are known features of drug-tolerant tumour phenotypes [16, 70]. Our analysis revealed a significant decrease of cell cycle activity in the residual tumours 12 days after TAC treatment and 7 days after cisplatin treatment (**Fig.3a and Supplementary Fig.12**). The decrease was followed by the recovery of cell cycle activity in the recurrent tumour samples. We revealed a strong correlation with the proliferating niche, which only diminished at the time points of maximum treatment response, and moderate correlations with the hypoxic, and moderate anticorrelation with the TME-associated niche (**Fig.3b**), corroborated by the spatial distribution of activity scores (**Fig.3c**). Furthermore, we assessed the enrichment of the adaptive mutability signature [71] (**Fig.3a-b** and **Supplementary Fig.12**), which is described by the decreased expression of DNA repair factors, thereby enhancing the possibility of gaining additional mutations and secondary resistance. We observed downregulation of this signature in the residual tumours consistent with the decrease of proliferating tumour cell abundance.

Continuing the functional characterisation, we assessed the enrichment of 50 gene sets from the Hallmarks MsigDB collection [72] with overrepresentation analysis (ORA). Significantly enriched terms (adjusted *P* < 0.05) were first analysed along the temporal axis (**Fig.3a and Supplementary Fig.13-14**). Hallmarks that showed downregulation at maximum treatment responses include *Myc target genes, Mtorc1 signalling, oxidative phosphorylation, unfolded protein response, peroxisome*, and *glycolysis*. Similar to this pattern, *hypoxia, Pi3k-akt-mTOR signalling*, and *androgen response* gene sets showed depletion in the residual tumours but interestingly, the lowest values correspond to 12 dpt in the cisplatin-treated residual tumours, indicating a more delayed response in these expression programs. Amongst these, *glycolysis* and *hypoxia* terms highly correlated with the hypoxic (10) compartment and *Myc target genes, oxidative phosphorylation, unfolded protein response, peroxisome*, and *Mtorc1 signalling* was associated with the proliferating (5) tumour niche (**Fig.3b**). Conversely, signatures associated with *epithelial-mesenchymal transition, complement*, and *Il2-stat signalling* were activated during MRD amongst others (**Fig.3a**). These Hallmarks were also highly correlated with TME niches (**Fig.3b**).

Building on these analyses, we then acquired specific gene signatures for each tissue compartment (**Supplementary Table 4**) by selecting the top-weighted genes (mean + 2 SD) and performed Over Representation Analysis (ORA) with the Gene Ontology (GO) [73] gene sets (**Fig.3d**). The proliferating niche (5) expression program was associated with terms related to cellular proliferation and replication, whereas the hypoxic niche (10) was correlated with altered metabolic processes. In the EMT niche (2), *cell motility* and *cell adhesion* terms were upregulated. TME niches rich in fibroblasts and connective tissue (4, 7, 9) exhibited expression programs related to extracellular structure maintenance and immune-rich (1, 11) compartments had upregulated defence response terms.

To assess the potential genetic variability of the different tumour niches, we performed copy number variation (CNV) inference, followed by unsupervised clustering based on the CNV signatures (**Fig.3e**). Consistent with the hypothesis that transient transcriptional signatures drive drug tolerance, our analysis of cellular and molecular niche compositions revealed that CNV clusters were not specifically attributed to cell types or tissue compartments (**Supplementary Fig.15**). Instead, CNV signatures differed significantly between tumours derived from different parental lines. Samples derived from parental tumour 1 were characterised by chromosomal amplifications in regions of chromosome 2, 8, 10, 13, and deletions in chromosome 17. Conversely, samples originating from parental tumour 2 (2.1, 2.2) showed amplified chromosomal regions in chromosomes 2, 5, 6, 11, 13, and 15, and deletions in chr 16.

Given the lack of response of the EMT tumour niche to treatment and the significant changes observed in the proliferating tumour compartment during MRD, we conducted detailed molecular profiling of these specific niches by performing pseudo-bulk DGEA on the subset of capture spots with high (> 0.80) compartment scores exhibiting dominant niche signatures (**Supplementary Fig.16a**). We found increased activity of *E2F1-4* and *Myc* transcription factors in the proliferating tumour niche, compared to increased activity of *Smad3, Jun, Sp1*, and *HIf1a* in the EMT niche (**Supplementary Fig.16b**). Differentially expressed genes were further linked to MAPK, oestrogen, and PI3K pathways in the proliferating tumour niche, while in the EMT tumour niche, they were associated with p53, JAK-STAT, and hypoxia pathways (**Supplementary Fig.16c**). The EMT tumour niche showed differentially expressed genes involved in epithelial-mesenchymal transition, TNFα signalling, interferon alpha/gamma response pathways, and hypoxia. In contrast, the proliferating tumour niche exhibited upregulation of G2M checkpoint, *E2f* and *Myc* target genes (**Supplementary Fig.16d-g**) from the Hallmarks collection.

To complete our spatiotemporal characterisation, we analysed the co-localisation of identified cell types across the primary, residual, and recurrent stages of the tumours, capturing dynamic reorganisation and compartmentalisation (**Fig.3f and Supplementary Table 5**). In primary tumours, we identified three distinct clusters: immune cells of the TME (1), stromal cells (2), and tumour cells (3). During MRD, stromal cells co-localised with lymphocytes and macrophages, forming a remodelled immune cluster (4). Residual tumour cells formed an EMT niche, with tumour cells showing no notable co-localisation with other cells (6) and a small fraction of remaining core tumour cells (5). In recurrent tumours, we observed marginally more separated immune clusters (7) from infiltrated stroma (9), as well as an EMT cluster (10) distinct from the bulk tumour cluster (8). Overall, the spatial organisation in recurrent tumours resembled that of the primary tumours. Furthermore, moderate anticorrelation was observed between the TME immune cells and the tumour cells, suggesting limited tumour infiltration, especially in the residual tumours.

In summary, the functional characterisation of MRD highlighted dynamic tumour reorganisation, with downregulated apoptotic and proliferative pathways, persistent EMT and immune-associated niches, and a senescence-like phenotype in drug-tolerant cells. These findings underscore the role of tumour plasticity and microenvironment interactions in therapy resistance and recurrence.

### Multimodal characterisation of cellular cisplatin uptake in mammary tumours

By combining ST and IMC we also examined at single cell resolution whether variations in drug uptake amongst different tumour subpopulations could contribute to the shifts in tumour cell proportions and cellular niches observed in residual tumours. For this purpose, we analysed 6 mg/kg cisplatin-treated tumours collected 4 hours, 24 hours, and 12 days after treatment focusing on the platinum (Pt) content with IMC. The selected tissue samples were stained with a custom panel of metal-conjugated antibodies (**Supplementary Table 12**) to segment and classify different cell types based on their protein expression signature. Using a pipeline with deep learning-based cell segmentation and cell phenotyping relying on pixel clustering with Pixie [74], we processed the IMC acquisitions (**Fig.4a**) and identified 15 different cell types, including tumour cells and several immune and stromal cell populations (**Fig.4b**).

**Fig. 4.**
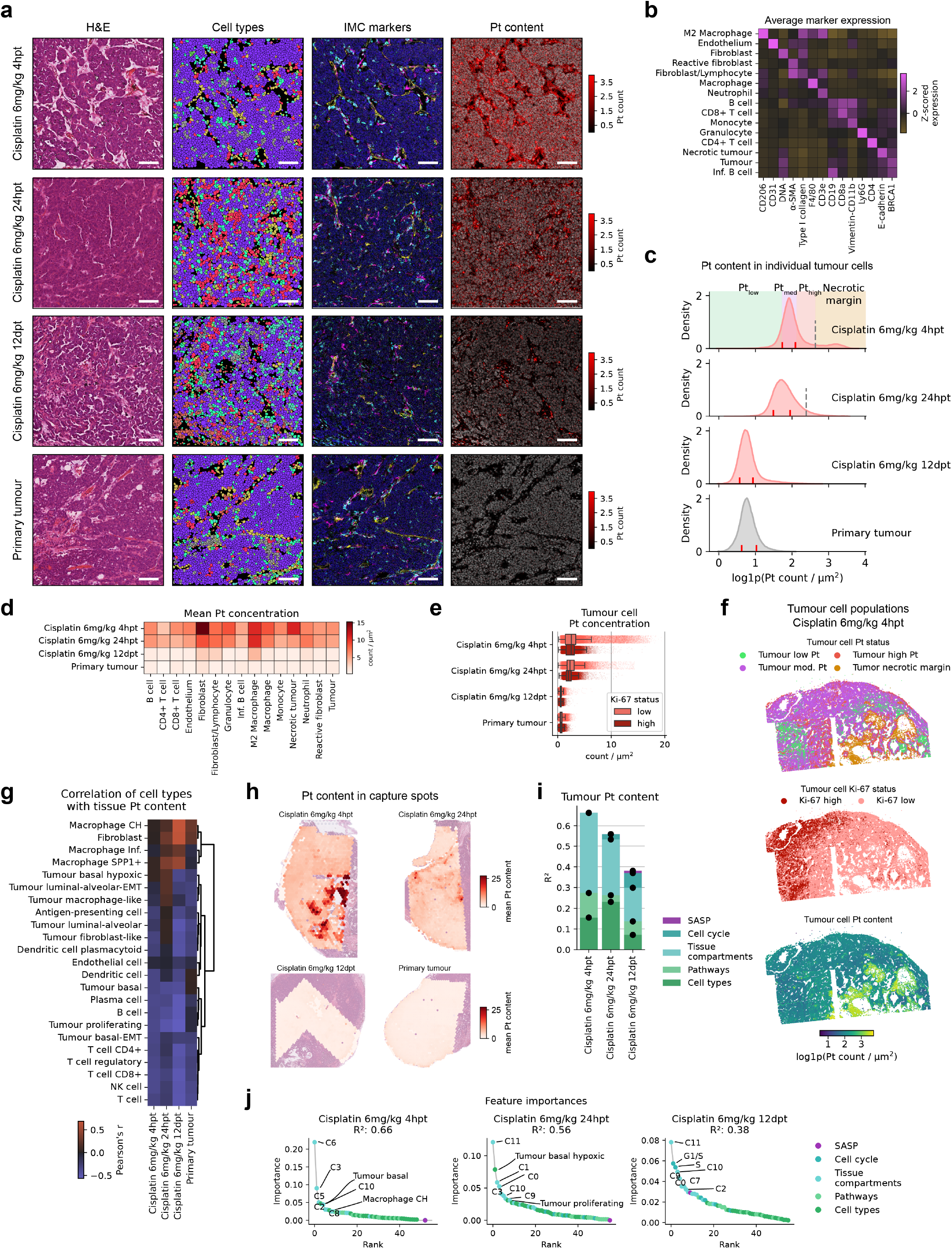
Spatial and transcriptional determinants of cisplatin uptake and distribution in mammary tumours. **a**, Representative IMC acquisitions of 6 mg/kg cisplatin-treated tumours collected at 4 hpt, 24 hpt, and 12 dpt, along with an untreated tumour sample. Corresponding histological images, segmented single cells, MIP of six IMC channels (light blue: CD206, yellow: CD3, green: CD3e, brown: F40/80, dark blue: DNA, pink: αSMA), and the total Pt signal summed across all detected isotopes are shown (scale bar: 75 µm). **b**, Heatmap of relative expression of metal-conjugated antibodies used to classify the different cell types. **c**, KDE plots showing the distribution of normalised Pt count / µm^2^ in tumour cells across different conditions. The left red line represents the 15th percentile, the right red line represents the 85th percentile, and the dotted line indicates the GMM cutoff. These thresholds were used to define four distinct classes of tumour cells: Pt_low_, Pt_mod_, Pt_high_, and necrotic margin tumour cells. **d**, Heatmap of mean Pt signal count / µm^2^ within the different cell types across different conditions. **e**, Pt count / µm^2^ within Ki-67+ proliferative and Ki-67-non-proliferative tumour cells across different conditions (centre line: median, box limits: upper and lower quartiles, whiskers: 1.5× interquartile range, salmon-pink dots: individual cells). **f**, Spatial map of tissue section with cells classified according to their Pt and Ki-67 status, as well as the measured Pt content 4 hours after 6 mg/kg cisplatin treatment. **g**, Heatmap showing the correlation of Pt content mapped to capture spots with the deconvolved cell type proportions from the ST data. **h**, Spatial maps showing Pt content measured only in tumour cells across different time points. **i**, Coefficient of determination of spatially informed random forest models trained using MISTy to infer the Pt content of tumour cells. Contribution of individual transcriptional feature sets is shown in the stacked bar plots. **j**, Rank plots showing the importance of individual transcriptional features in explaining intracellular Pt content.

First, the intracellular Pt signal intensity was assessed to infer cisplatin uptake within the tumour tissue. Since Pt content in the tissue was directly measured by IMC detectors without signal amplification, we evaluated the isotopic distribution and kinetics of Pt within the tissue. The measured abundances of ^194^Pt, ^195^Pt, and ^196^Pt closely matched the expected isotopic distribution (^190^Pt: 0.01%, ^192^Pt: 0.78%, ^194^Pt: 32.9%, ^195^Pt: 33.8%, ^196^Pt: 25.2%, ^198^Pt: 7.36%) [75], and exhibited the anticipated clearance kinetics. However, the intensity of ^198^Pt was significantly higher than expected and showed no spatial correlation with the other isotopes, leading us to exclude it from further analysis (**Supplementary Fig.17**). Similarly, the signals for ^190^Pt, and ^191^Pt were indistinguishable from those observed in untreated tumour samples, suggesting that these isotopes were below the detection limit and were therefore omitted from the analysis. Overall, the IMC acquisitions show high extratumoural concentrations but even distribution of Pt inside the cells. (**Fig.4a**). Pt concentration was the highest at 4 hours after cisplatin treatment, followed closely by 24 hours post-treatment (hpt) and decreased drastically in the residual tumour, where most cells lacked Pt signal, similarly to the untreated tumour, indicating that residual tumour cells have largely removed Pt adducts generated by cisplatin treatment [76] (**Fig.4c and Supplementary Fig.18a-b**). We further classified the tumour cells based on Pt content: Pt_low_ tumour cells below the 15th percentile, Pt_mod_ cells between the 15h and 85th percentile, and Pt_high_ cells above the 85th percentile of the Pt content distribution. Moreover, we introduced a Gaussian mixture modelbased (GMM) thresholding to examine cells with prominent Pt content visible at 4 hpt and 24 hpt on kernel density estimate (KDE) plots. We analysed the spatial distribution of these cells and observed that the tumour population with very high Pt content resides in the perinecrotic areas of the tumour, consequently, we labelled them as necrotic margin tumour cells (**Supplementary Fig.18c**).

We followed the analysis by examining the Pt content in the cell types of the TME. The highest Pt concentration was observed in the fibroblasts and M2 macrophages (**Fig.4d**). This corroborates previous findings reporting extensive collagen binding of Pt in tumours after cisplatin treatment [77]. In addition, the M2 macrophages retaining higher Pt content in the residual tumours may contribute to the downregulation of the immunosuppressive microenvironment reported after cisplatin treatment [78]. Neighbour enrichment analysis further revealed that M2 macrophages are densely co-located with fibroblasts with the exception of the residual tumour (12 dpt). This is also evident by visually inspecting the fibrotic capsule surrounding the tumour cells (**Supplementary Fig.5**). Pt_mod_ tumour cells, which constitute the majority of the tumour mass, exhibit negative enrichment scores with most other cell types (**Supplementary Fig.19**) across the four different conditions.

Next, we classified tumour cells based on their Ki-67 status to examine if active cell cycling results in alterations in cisplatin uptake (**Fig.4e**). We have found little variation in mean Pt content in cycling and non-cycling cells, suggesting uniform drug distribution across the cells at different time points. Cells observed at 4 hpt with notably higher Pt concentration in non-cycling (Ki-67 low) tumour population originate from the necrotic margin, where we also noted the absence of cycling tumour cells (**Fig.4f**). Of note, significant Pt accumulation was also observed in the necrotic tissue, even as early as 4 hpt, preceding cisplatin-induced cell death [16]. These results further suggest that the presence of fibroblasts, immune cells and necrotic tissue might influence Pt distribution within the tumour.

To explore the potential influence of transcriptional features on drug distribution, we analysed the co-registered ST data by mapping the Pt intensity from IMC measurements to the capture spots (**Fig.1i**). We first examined the correlation of intracellular Pt content with the inferred cell type abundances (**Fig.4g**). Fibroblasts, *Spp1*+ macrophages, and CH macrophages moderately correlated with Pt content, while transcriptional tumour cell states only showed weak correlation with Pt with a value of 0.25 Pearson’s r at 24 hpt exhibited by the hypoxic tumour cells followed by the luminal-alveolar-EMT cells. Weak anticorrelation was also found at 4-24 hpt exhibited by lymphocytic immune cells and proliferating tumour cells.

To further investigate transcriptional features that might influence drug distribution in the tumour cells specifically, we utilised MISTy [79], an explainable machine learning framework, to infer the Pt content measured in the tumour cells (**Fig.4h**), by leveraging spatially-informed relationships derived from a selected transcriptional feature set. In the feature set, we introduced tissue compartments predicted with Chrysalis (**Supplementary Fig.20 and Supplementary Table 6**), alongside the inferred cell type proportions, pathway activities, SASP, and cell cycle scores. The trained decision tree model explained 66% of the variance in Pt content at 4 hpt (**Fig.4i**). This decreased steadily as more time elapsed from cisplatin treatment (24hpt: 0.56 *R*^2^, 12 dpt: 0.38 *R*^2^). The majority of the explained variance is attributed to the tissue compartments followed by the inferred cell type proportions. Analysing the importance score of the fitted models revealed that the most important features are the tissue compartment and cell types related to hypoxic tumour cell neighbourhoods found next to necrotic cores (Compartment 6, 11, 3, tumour basal hypoxic), underscoring the importance of these structures capturing Pt.

Overall, these findings suggest that the drug tolerance mechanism of the residual subpopulations we identified is not directly linked to reduced drug uptake, which is consistent with previous immunohistochemical experiments using a Pt-DNA adduct-specific antibody [16].

### Tumour niches share molecular characteristics with MRD in human breast cancer

To translate our findings from the mouse model into human disease, we analysed human BRCA1-deficient breast cancer ST samples, characterising intratumoural heterogeneity and molecular features across species. The collected primary and post-neoadjuvant chemotherapy residual tumour samples were first annotated by a pathologist and capture spots containing tumour tissue were selected for further analysis (**Supplementary Fig.21a**). Pseudo-bulk DGEA between the primary and residual tumour spots revealed 2023 upregulated genes in the primary tumours, from which 203 were also found in the 799 differentially expressed genes in the mouse dataset (**Fig.5a**). While we identified 1896 upregulated genes in the human tumours and 1000 in the mouse tumours, 188 genes were shared between the two species.

**Fig. 5.**
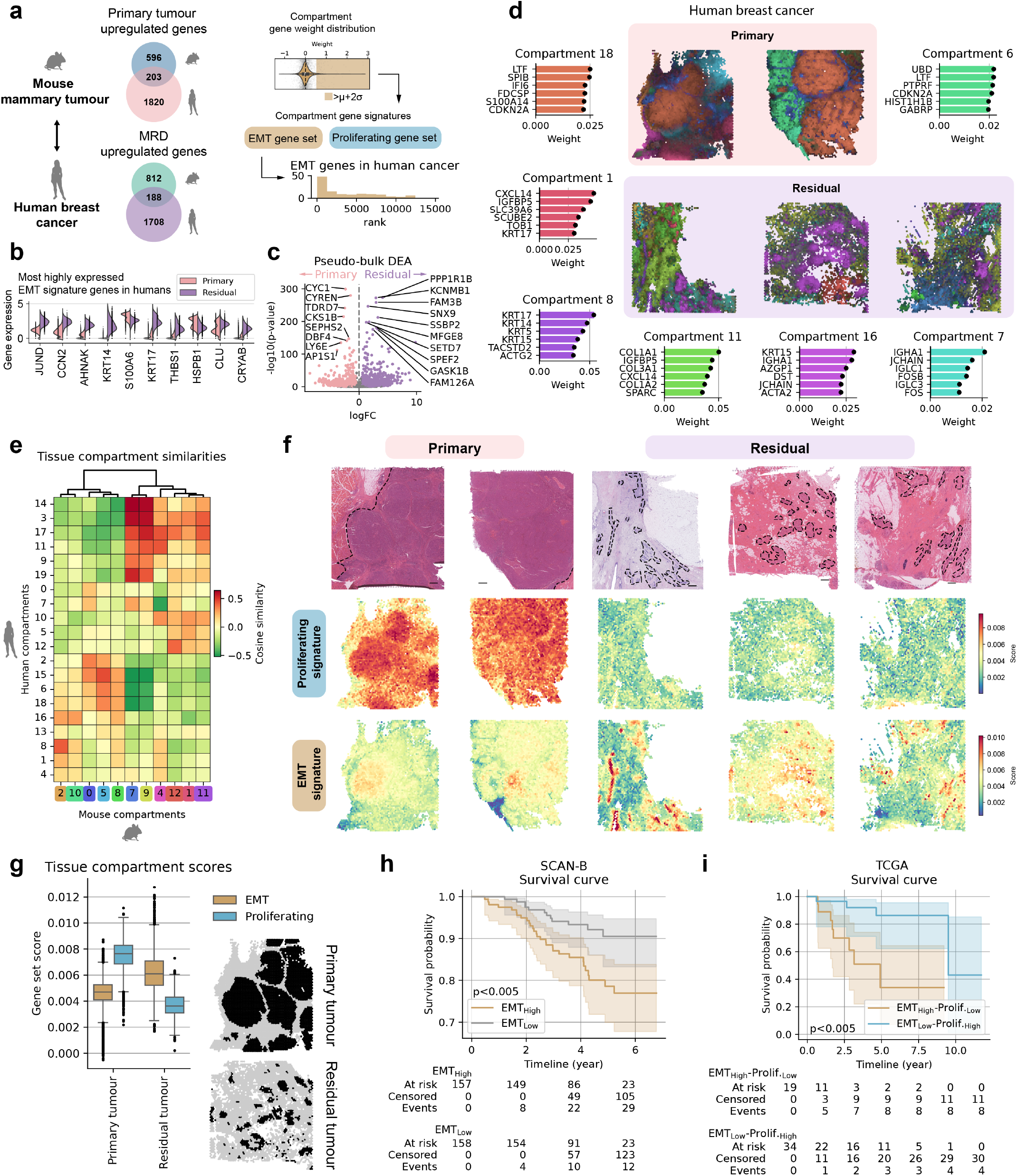
Molecular characteristics of tumour niches are shared with MRD in human breast cancer. **a**, Mouse primary tumours share 203 upregulated genes with primary human breast tumours and 188 genes in their residual state. EMT and proliferating compartment gene signatures were determined by taking top-weighted genes exceeding the mean plus two standard deviations (µ+2σ). **b**, Top ten most highly expressed EMT signature genes in the human primary and residual tumours. **c**, Pseudo-bulk DGEA of primary and residual human breast cancer ST samples. **d**, MIP of the identified tissue compartments on the primary and residual human tumour tissue sections. **e**, Heatmap showing cosine similarity between the mouse and human cellular niches identified with Chrysalis. **f**, Spatial visualisation of the established EMT and proliferating signatures on the primary and residual human tumour tissue sections. While the proliferative signatures significantly decrease after treatment, the EMT signature remains unchanged, similar to the findings in our mouse model. **g**, Distributions of EMT and proliferative signature scores across individual capture spots in primary and residual human tumour tissue (centre line: median, box limits: upper and lower quartiles, whiskers: 1.5× interquartile range, black dots: outliers). Capture spots were identified as tumour-containing based on expert annotations of the corresponding H&E-stained sections (black spots: tumour tissue, light grey spots: rest). **h**, Kaplan–Meier survival curves of TNBC patients from the SCAN-B dataset, stratified based on the expression of the EMT signature (log-rank test *P* < 0.005). **i**, Kaplan–Meier survival curves of patients belonging to the EMT_High_-Proliferating_Low_ and EMT_Low_-Proliferating_High_ classes (log-rank test *P* < 0.005).

Next, we leveraged our gene signatures for the EMT and proliferating niches. From the 120 genes identified in driving the EMT tumour niche in mouse tumours, 50 genes were amongst the top 10% of most highly expressed genes in the human tumours as well. Amongst the most highly expressed genes from the EMT signature (**Fig.5b**), several exhibited apparent differences in expression between the primary and residual states. Moreover, many of these genes play key roles in driving tumour progression in humans, such as suppressing p53-dependent senescence (*JUND*) or modulating cell differentiation, motility, and metastasis (*CCN2, AHNAK, THBS1*). Differentially expressed genes in the pseudo-bulk DGEA (**Fig.5c and Supplementary Table 7**) revealed upregulated genes, such as *PPP1R1B*, that is linked to Trastuzumab resistance in HER2-positive breast cancer [80] and *SNX9* that promote metastasis in different cancers [81]. Upregulated genes in the primary tumours, on the other hand, included many cell cycle regulators (*e*.*g*., *CYC1, CYREN, CKS1B*).

We again leveraged Chrysalis to identify molecular tissue compartments (**Fig.5d and Supplementary Table 8**). The bulk of the primary tumour tissue (Compartment 6 and 18) was characterised by genes such as *LTF, SPIB*, or *IFI6* with notable immune infiltration present in Compartment 10, represented by a variety of immunoglobulin genes, amongst others (**Supplementary Fig.21b-c**). In the residual tumour samples, we characterised three distinct tumour niches (1, 8, 16) associated with malignant epithelial cells, confirmed by histopathological examination. Additionally, an inflammatory microenvironment (Compartment 11) was observed surrounding certain residual tumour islands exemplified by *CXCL14*), and the presence of immune cell aggregates (Compartment 7).

Building on the initial characterisation, we quantified the cellular niche similarity between the mouse MRD model and residual disease in human TNBC (**Fig.5e**). Using cosine similarity, we observed moderate alignment of cellular niches across species. Notably, the EMT-associated mouse niche (2) exhibited higher similarity to the human compartment 8 and 16, while the proliferating tumour niche (5) also showed parallels with human counterparts (6, 15, 18). TME-associated compartments displayed moderate similarities, with the ECM-rich mouse niches (7, 9) showing the strongest alignment, followed by immune-rich niches (12, 1, 11). Additionally, the pseudo-bulk DGEA performed on the high scoring (> 0.80) primary (6, 18) and residual (1, 8, 16) tumour-associated tissue compartments showed further similarities with the mouse mammary tumour EMT and proliferating niches, such as downregulated *E2F1*, and *MYC* activity, and the enrichment of EMT Hallmark gene set in the residual tumour (**Supplementary Fig.22**). We noted differences in JAK-STAT, TNF signalling and interferon signalling, which is downregulated in the human residual tumour niches but upregulated in mice, consistent with the substantial immune infiltration we described.

To study the potential relevance of the expression programs of the proliferating and EMT niches in human MRD, we assessed their average expression score in the human ST samples (**Fig.5f**). We found elevated EMT signature expression score in the residual tumours, correlating with the presence of malignant epithelial cells. Whereas in the proliferating tumours, the signature’s expression was moderately elevated in a small number of capture spots. In contrast, the proliferating signature was significantly higher throughout the primary tumour tissue compared to the residual tumour samples. Using the histopathological annotations, we selected residual and primary tumour-containing capture spots, quantified their expression profiles, and identified significant changes in proliferating and EMT score distributions across the two treatment groups (**Fig.5g**).

As differences in EMT and proliferating signatures were evident in the examined spatial samples, we examined their relevance in patient survival on a larger scale. We assessed the prognostic effect of these signatures on patient survival by leveraging two extensive breast cancer genomics databases, namely The Sweden Cancerome Analysis Network - Breast (SCAN-B) [82] and The Cancer Genome Atlas Breast Invasive Carcinoma Collection (TCGA-BRCA) [83]. We specifically analysed the association of our EMT signature expression with patient survival amongst the TNBC patients (*n*=315) in the SCAN-B data source (**Fig.5h**). Stratifying patients to EMT_High_ and EMT_Low_ groups showed significant differences in survival outcomes as assessed by Kaplan-Meier analysis and log-rank test with the EMT_High_ group associated with poorer survival outcomes. For the TCGA TNBC patients (*n*=163), we introduced a K-means clustering-based approach identifying EMT_High_-Proliferating_Low_ and EMT_Low_-Proliferating_High_ patients to assess the link between proliferating and EMT signatures (see Methods). EMT_High_-Proliferating_Low_ was associated with significantly poorer survival outcomes (**Fig.5i**). Hence, the molecular characteristics of tumour niches identified in our BRCA1-deficient mouse mammary tumours closely align with those in human breast cancer, particularly in residual disease, with shared EMT and proliferative gene expression programs that correlate with poor patient survival outcomes.

## Discussion

Our study presents a comprehensive multimodal approach combining scRNA-seq, ST, and IMC to map the spatiotemporal profile of MRD using the KB1P mouse model for hereditary breast cancer. Our profiling of cellular niches, pathways, and key regulators of residual disease conserved across mice and humans, provides the opportunity to identify MRD-associated genes as potential therapeutic targets.

Through transcriptomic profiling in our mouse model, we propose the critical role of EMT niche-associated cell states and expression programs of residual tumour cells in surviving various chemotherapy treatments, involving genes such as *WFDC2, ANXA1, SLPI, LTF*, and *TACSTD2* (**Supplementary Table 4**). Additionally, we validated the presence of this expression program in human MRD and confirmed its association with poor survival outcomes. EMT has long been linked to drug resistance through mechanisms such as increased drug efflux, slowing down proliferation, avoiding apoptotic signals, or fostering immune evasion [84–88]. Specifically, in breast cancer, the EMT-related transcription factors Twist, Snail, and Foxc2 have been shown to upregulate ABC transporters [89]. Further evidence showed that the activation of Twist and Snail confers resistance to doxorubicin and docetaxel [90, 91]. In TNBC cell lines, RHOJ, which is preferentially expressed in EMT cells, enhances the DNA damage response, rendering these cells more resilient to treatment [92]. Consistent with these findings, our data emphasize the need for more effective targeted strategies against EMT cells.

Our spatiotemporal profiling uncovered a rich ecosystem composed of both tumour cells and surrounding stromal and immune cells of the TME. We described highly proliferative, basal and luminal-alveolar tumour cell lineages transitioning into dormant EMT phenotypes, which showed significant enrichment in MRD. The emergence of EMT [93] and dormancy [19, 94] have been shown to be regulated or modulated by the TME, underscoring the need to study MRD in an immunocompetent model. Additionally, we identified *Spp1*+ and complement-high macrophages, previously reported to contribute to a pro-tumour microenvironment [36, 95], that may allow surviving EMT and other tumour cell types to revert to a proliferative state. Even though our data do not directly determine how or when the dormant EMT phenotype of residual tumour cells is triggered, they suggest that both tumour-intrinsic and tumour-extrinsic factors play a role in therapy response.

Cell type deconvolution further revealed the *in situ* localisation of these cells, while cellular niche identification uncovered the organisation of these cells with distinct temporal dynamics during tumour progression, confirming that the EMT niche remains unaffected by the treatment. Our findings demonstrated the high similarity in spatial organisation between primary and recurrent tumours, highlighting the highly dynamic and reversible nature of residual disease. We showed that the majority of tumour cells are depleted at the time of maximum effect of both platinum- and combination-based chemotherapy treatments. However, the persisting residual cancer cells eventually regrow into recurrent tumours that mirror the intratumoural heterogeneity observed in the primary tumours. This process is supported by a permissive TME, which similarly to the tumour cells, undergoes extensive remodelling following treatment. While our analysis does not identify a specific tumour cell subpopulation responsible for recurrence, it provides valuable insights into the heterogeneity and plasticity underlying drug tolerance. These findings, validated across two different therapeutic strategies, suggest that the primary mechanisms of drug tolerance may be drug-independent.

Furthermore, we revealed several key pathways and processes driving tumour progression or associated with critical stages during MRD and tumour recurrence, such as the steep downregulation of MAPK pathways and an upregulation of JAK-STAT. We characterised several functional components attributed to key tumour or TME-associated tissue compartments. Notably, numerous ECM-remodelling and immuno-suppressive processes were spatially correlated with the persisting cells in the EMT niche.

We developed a multimodal assay to detect intracellular Pt content and investigate its potential effect on transcriptional changes in our cisplatin-treated samples. We confirmed that necrotic areas and fibroblasts act as major sinks for Pt within the tumour tissue. Additionally, the identified macrophages with accumulated Pt two weeks after drug treatment, which may contribute to the decreased effectiveness of the immune system in clearing tumour cells, in line with previous works showing that cisplatin-stimulated macrophages promoted EMT in ovarian cancer [96]. Consistent with previous findings [16], we demonstrated that different tumour cell populations exhibit similar drug uptake, irrespective of their transcriptional state. This observation suggests that drug tolerance may not result from uneven distribution of DNA damage across tumour cell populations.

In conclusion, we present comprehensive data capturing the unique transcriptional features and states of cancer cells, including drug-tolerant populations, as well as the dynamics of the TME, using spatiotemporal profiling with ST to better understand MRD and tumour recurrence. We laid the groundwork for deeper mechanistic understanding and validation of the identified processes, enabling the development of better diagnostic and therapeutic strategies for MRD in TNBC.

## Methods

### Experimental work

#### Collection of mouse mammary tissue

All animal experiments were approved by the Animal Ethics Committee of the canton of Bern (BE60/2023) and are in accordance with the current Swiss Acts on Animal Experimentation. Two different parental tumours (T1, T2.1-2.2) were used in this study. While they are both derived from the K14*cre*;*Brca1*^F/F^;*Trp53*^F/F^ (KB1P) mouse mammary tumour model, Tumour 1 is a GFP-labelled *Brca1*^-/-^;*p53*^-/-^ mammary tumour. Samples from Tumour 2 were transplanted and collected in two different experiments (2.1 and 2.2). The DMSO-frozen tumour pieces were thawed, washed with PBS and cut into small pieces (∼2 mm in diameter, approximately the size of a needle head). The pieces were transplanted in the fourth mammary fat pad (on the right side or both sides) of 2-4 months FVB/NJ mice. Mammary tumour size was measured by calliper measurements and tumour volume was calculated using the following formula: length × width^2^ / 2. Animals were randomly assigned to the treatment groups. When the tumours reached a size of approximately 8 × 6 mm, the tumour-bearing mice were either sacrificed to collect the primary tumours or treated with a single dose of TAC therapy as a combination of 2.5 mg/kg doxorubicin (Teva, Pharmacode 4439164) injected intravenously, 12.5 mg/kg docetaxel (Teva, Pharmacode 6984902) and 120 mg/kg cyclophosphamide (Baxter, Pharmacode 2731162), both injected intraperitoneally or with a single dose of cisplatin 6 mg/kg (Teva, Pharmacode 4333164) injected intravenously. The residual tumours were collected 7 or 12 days after treatment and the recurrent tumours were collected once they reached the size of approximately 8 × 6 mm again. Some cisplatin-treated tumours were collected 4 or 24 hours after treatment for profiling with IMC. Animals were sacrificed with CO_2_. Animal technicians were blinded to the respective treatment groups while measuring the tumours. Mammary fat pads of mice that never underwent tumour transplantation were collected as well. The mammary tumours were harvested at different experimental endpoints and processed either for tissue collection and fixation or for scRNA-seq. For ST and IMC, the tissue was fixed in 4% paraformaldehyde immediately after harvesting. The fixed samples were trimmed, embedded in paraffin and stained with H&E for a first histological assessment.

#### Mouse mammary tumour dissociation and FACS

Mice were sacrificed, tumours were collected and tissue is directly dissociated into single cells. First, the tumour was washed with PBS and the surrounding tissue (such as blood vessels, fat, and connective tissue) was removed. The tumour was then minced into very small pieces and collected together with PBS into a 15 ml tube. The sample was centrifuged at 600*g* for 5 min at 4°C. The supernatant was discarded and the tumour pieces were incubated in 8.5 ml of digestion mix (0.02 g collagenase type IV from *Clostridium histolyticum* (Sigma, C5138) in 10 ml of Advanced DMEM/F-12 (Gibco, Thermo Fisher Scientific 12634010) supplemented with a final concentration of 100 mM HEPES pH 7.5 (Sigma, H4034), 1× GlutaMAX (Gibco, 35050038), 50 units/ml penicillin-streptomycin (Gibco, 15070063) at 37°C for 15 min while shaking. The tissue was further mechanically dissociated by gently pipetting up and down. This enzymatic and mechanical dissociation step was repeated twice. Next, the samples were centrifuged at 600*g* for 5 min at 4°C, washed with the supplemented Advanced DMEM/F-12 and centrifuged again. The pellets were resuspended in 4 ml of DNase solution (45 µl of DNase I Solution 2500 units/ml (Thermo Fisher Scientific, 90083) in 5 ml of the supplemented Advanced DMEM/F-12) while gently pipetting up and down for 5 min. The cells were centrifuged again at 600*g* for 5 min at 4°C. The pellets were resuspended in 4 ml of the supplemented Advanced DMEM/F-12 and shaken by hand for 2 min. The tissue should be completely digested, if necessary, visible tissue pieces were removed from the liquid fraction with a pipette. The cells were centrifuged one last time at 600*g* for 5 min at 4°C and the pellets were resuspended in 600-800 µl freezing medium (Recovery™ Cell Culture Freezing Medium, Gibco, 12648010). Finally, the cells were filtered through 40 µm Flowmi Cell Strainers (Sigma, BAH136800040) into a cryogenic tube. 10 µl of the suspension was taken to manually count the cells and estimate the number of single cells viable before freezing. If the cells were not fully dissociated, they were centrifuged, resus-pended, and filtered again. The cells were frozen down and stored at -150°C until all the samples were collected.

Frozen vials were thawed and stained with the near-IR fluorescent reactive dye of the LIVE/DEAD™ Fixable Dead Cell Stain Kits (Invitrogen, Thermo Fisher Scientific, L34960) according to the manufacturer’s instructions. All the steps were performed protected from light to keep the fluorescence signal. The tubes and pipette tips were coated with 0.1% BSA (Bovine Serum Albumin Fraction V, Roche, 10735086001) in PBS to prevent cell loss and a wide bore pipette was used for gentler handling. The cells were resuspended in a solution of 0.4% BSA in PBS and kept on ice. The sorting of cells was done using a Beckman Coulter MoFlo ASTRIOS EQ cell sorter with a 120 µm nozzle at 10 psi system pressure in the Beckman Coulter IsoFlow sheath fluid. Sample and collection devices were cooled to 4°C. Single-stained controls and unstained cells were used to set the gates for the cell viability (ex. 640 nm, em. 795 nm) and for their GFP expression (ex. 488 nm, em. 526 nm). After gating out doublets, contaminating red blood cells and the dead cells, 3 primary tumours and 3 residual tumours were sorted according to their GFP expression. Tumour cells are enriched in the GFP-positive population, cells of the TME in the GFP-negative population. At least 25,000 cells per condition were sorted for scRNA-seq.

#### scRNA-seq data generation

The cells were processed directly after the FACS sorting. The scRNA-seq libraries were prepared at the Next Generation Sequencing Platform of the University of Bern. The 10x Genomics Chromium Single Cell 3*′* Library and Gel Bead Kit v3.1 was used according to the manufacturer’s standard protocol (cDNA amplification in 14 cycles). A targeted cell recovery of 8000 cells per sample was aimed. The sequencing of the library was performed on the NovaSeq 6000 platform (Illumina) using a S4 flow cell.

#### Collection of human tissue

All human breast tissue obtained from the University of Geneva and Centre Léon Bérard was approved by the local ethical committees in Geneva (CCER 2019-00004) and Lyon. All biopsy and tumourectomy tissue samples were obtained from women with germline BRCA1 mutations, with or without neoadjuvant chemotherapy. Participants were not specifically recruited for this study. The formalin-fixed, paraffin-embedded (FFPE) samples were profiled using 10x Visium.

#### ST data generation

FFPE mouse mammary and human breast tumour tissues were histopathologically assessed and samples with minor necrosis were selected for further ST profiling. RNA quality control of 32 mouse and 5 human samples was determined and samples were further analysed following the Visium V1 (10x Genomics) procedure; 1 human sample was processed with the Visium V2 CytAssist procedure. Briefly, 5 µm thick sections were placed on Visium slides with the ROI within the capture area and manually H&E stained and scanned with a S360 Nanozoomer (Hamamatsu) using a 40x magnification. cDNA libraries were constructed according to the Visium Gene Expression User Guide (10x Genomics) and sequenced by the Next Generation Sequencing Platform at the University of Bern on an Illumina NovaSeq 6000 System on different flow cells with 100-300 cycles.

#### Histopathological assessment

Whole-slide H&E stained images from the collected mouse mammary tumours and the human tissue samples were used. All the slides were scanned on Nanozoomer S360 (Hamamatsu) with a 40x magnification (0.23 µm/pixel resolution). The H&E images from Visium tissue sections were examined and annotated using QuPath [97] (version 0.4.3). The following regions were annotated and excluded from the downstream analysis of ST: necrosis, lymph node, and surrounding fat tissue.

#### IMC staining and measurement

FFPE cisplatin-treated tumours sections were used for both ST and IMC analyses. First, the sections of 5 µm for ST were cut, directly followed by the sections of 3.5 µm for IMC (serial sections), and finally, a last section of 3.5 µm was cut for H&E staining as a control to assess the tissue morphology of the IMC sections as well as to define the regions of interest (ROI). The slides were processed and stained based on previous work [98]. Briefly, FFPE sections on Superfrost Plus slides were dewaxed by Xylol and rehydrated stepwise by 100%, 94%, and 70% ethanol and rinsed with distilled water. Heat-induced epitope retrieval was conducted in a Decloaking Chamber™ NxGen (Biocare Medical, DC2012) at 110°C in citrate buffer pH 6 (Sigma, C9999) for 15 min. After cooling, the sections were washed with PBS and blocked with protein blocking (Tris-buffered saline 1× pH 7.5, 0.1% NaN_3_ (Merck, 106688), 3% Goat serum (Thermo Fisher Scientific, 16210-064), 0.5% Casein (Sigma, C-8654), 0.025% Tween20 (Sigma, P1379)) for 1h in a hydration chamber at room temperature. The samples were incubated with a metal-conjugated antibody cocktail (19 antibodies, **Supplementary Table 2**) overnight at 4°C. The samples were washed twice with PBS + 0.05% Tween-20 and twice with PBS only. The sections were incubated with a 1:100 dilution of 25 µmol/L Ir-Intercalator (Fluidigm, 201192A) for 30 minutes at room temperature. The samples were quickly rinsed in distilled water and air-dried before acquisition.

The stained and dried samples were then inserted into the Hyperion Tissue Imager (Standard BioTools), where a 1 µm diameter UV laser ablated the tissue at a faster rate of 200 Hz in the selected ROIs. Tissue from the ablation spot is vaporised with each laser shot, and the plume containing the heavy metals present in the tissue is transported into the Helios mass cytometer. Here the elements are atomised and ionised in the inductively coupled plasma and pulsed into the time-of-flight (TOF) chamber for detection based on their mass-to-charge ratio. The mass cytometry data files (MCD) files were acquired using the CyTOF software version 7.0.8493. In total, 44.4 mm^2^ of tumour tissue was acquired and analysed across 7 different samples.

### Computational analysis

#### scRNA-seq data processing and cell phenotyping

Read mapping and counting of 3-3 primary and residual tumour libraries (processed in a single sequencing run) was performed with Cell Ranger 6.1.2 (10x Genomics), using the mm10 reference genome for *Mus musculus*. Processing was carried out using SCANPY [99] in a Python v3.9-based environment. Quality control was performed by removing low-quality cells based on thresholds defined by gene count, and the number of expressed gene distributions (calculated with scanpy.pp.calculate_qc_metrics), as well as mitochondrial gene percentage (< 5%). This resulted in 8470 cells from the control tumours (5665 tumour cells and 2805 cells from the microenvironment) and 9502 cells from the residual tumours (1384 tumour cells and 8118 cells from the microenvironment). After integration with SCANPY, major cell types were manually annotated based on top-ranked genes calculated with the Wilcoxon test (scanpy.tl.rank_genes_groups) performed on Leiden clustering (scanpy.tl.leiden, resolution=0.2) results. Cell subpopulations were detected with either subclustering (scanpy.tl.leiden, resolution=0.6) of the major cell type clusters or for T cells, algorithmic T cell detention with TiLPRED [100]. Tumour cell populations and lineages were characterised using Wilcoxon tests in combination with signatures derived from previous works [34, 35]. To visualize the single-cell data, t-SNE plots were generated with SCANPY by processing the normalised, log1p transformed count matrix data with scanpy.tl.pca (svd_solver=‘arpack’), followed by, scanpy.pp.neighbors, and scanpy.tl.tsne with the default parameters.

#### Pseudo-bulk DGEA

Pseudo-bulk data from the tumour cells of 3-3 primary and residual scRNA-seq samples were generated by aggregating the raw counts with decoupleR [101]. Genes with low expression were filtered with decoupler.plot_filter_by_expr. For differential gene expression testing, PyDESeq2 [102] was used with the Benjamini-Hochberg procedure to control the false discovery rate. Log2 fold changes (log2FC) were corrected with the apeglm [103] shrinkage method. Differentially expressed genes were classified based on adjusted *P* values (< 0.05) and absolute log2FC (> 0.5) values.

DGEA of ST primary and residual mouse mammary tumour samples (*n*_Primary_=10, *n*_Residual_=9, samples used for the multi-modal IMC analysis were excluded) were performed by aggregating raw counts from all capture spots classified as tumour by histopathological annotation. This includes both cisplatin and TAC therapy-treated tumours, as well as both residual tumour sampling timepoints (7 dpt, 12 dpt).

For the human primary (*n*=2) and residual tumours (*n*=3), DGEA was performed similarly by aggregating raw counts from all capture spots classified as malignant epithelium by pathologists.

#### RNA velocity

Velocyto [104] (velocyto.run10x) was run on all six samples to calculate the ratio of spliced and unspliced mRNAs using the BAM files generated by Cell Ranger containing the alignment of reads to the reference genome. RNA velocity analysis was performed on the output loom files with scVelo [105] to infer latent cellular dynamics. Specifically, scvelo.pp.filter_and_normalize (min_shared_counts=20, n_top_genes=2000) was used to select highly variable genes, scvelo.pp.moments (n_pcs=30, n_neighbors=30) was used to compute moments for velocity estimation based on cell similarities. Splicing kinetics modelling to estimate RNA velocities was performed with scvelo.tl.recover_dynamics. Lastly, scvelo.tl.latent_time was used to infer an intrinsic latent time reflecting the progression of cellular states and visualised on the original t-SNE embedding of the tumour cells.

#### ST data processing

FastQ files were processed with Space Ranger (10x Genomics) *count*. Sequence reads were mapped to reference transcriptomes mm10 (mouse) and GRCh38 (human) with Mouse Transcriptome v1 and Human Transcriptome v1 probe set files. Loupe Browser from 10x Genomics was used for manual fiducial frame alignment and manual selection of tissue-covered spots for each sample. Quality control was performed with SCANPY based on our examination of the spatial and overall distribution of the total counts and number of detected genes in the capture spots. Manual thresholds were defined to remove low-quality spots (insufficient number of counts, oversaturated capture spots due to lateral diffusion) for each individual sample, and genes expressed in less than 10 capture spots were also removed (**Supplementary Table 9 and Supplementary Table 10**). Normalisation and log-transformation of the counts were performed with scanpy.pp.normalize_total (target_sum=1e4, exclude_highly_expressed=True) and scanpy.pp.log1p.

To map histopathological annotations to capture spots (**Supplementary Fig.3**), QuPath project files were queried using the Paquo (Bayer AG) Python package. Annotation polygons were then extracted and matched to capture spots using the Shapely Python package by determining whether the centroid of each capture spot falls within a polygon. To enhance the visualisation of H&E sections, the OpenCV2 Python package was used to generate segmentation masks for background removal with contour detection (cv2.Canny, cv2.dilate, cv2.erode, cv2.findContours).

#### Cell type deconvolution

To deconvolve ST data, celltype deconvolution was performed with cell2location [49]. This probabilistic model integrates scRNA-seq reference data with spatial data to map cell-type-specific expression patterns, estimating the abundance of cell types at each spatial location. We leveraged our scRNA-seq dataset to train the single-cell regression model to extract cell type-specific expression signatures. To achieve this, genes were first filtered (adata_ref, cell_count_cutoff=5, cell_percentage_cutoff2=0.03, nonz_mean_cutoff=1.12) and the regression model was trained with the maximum number of epochs set to 1000. The cell2location model was then trained (N_cells_per_location=30, detection_alpha=20) with the maximum number of epochs set to 3000. Finally, per-spot cell densities were calculated by dividing the cell type abundance values (**Supplementary Fig.23-24**) with the total number of inferred cells. Sample level cell type fractions were calculated by summing cell densities across all capture spots for each cell type and normalised by the total sum of all proportions, yielding a final set of values representing the contribution of each cell type to the overall tissue composition.

#### Cellular niche inference

Cellular niches (tissue compartments) were inferred by Chrysalis [56] on Visium ST data. Chrysalis builds on archetypal analysis to identify spatially and functionally distinct tissue compartments. It operates by fitting a simplex to the latent representation (principal components) of the gene expression matrix. The vertices of this simplex correspond to archetypes, specific combinations of PCs that represent unique biological functions, and define the tissue compartments. Chrysalis decomposes the count matrix into a basis matrix containing a set of tissue compartments with values summing up to one, and a weight matrix that represents the contribution of each gene to these compartments. For the mouse mammary tumour dataset, spatially variable genes (SVGs) were first calculated on the normalised and log-transformed samples (chrysalis.detect_svgs, parameters: min_morans=0.025, min_spots=0.05). This was followed by a sample integration with HarmonyPy (Python implementation of Harmony [106]), using sample ID as a covariate to integrate over. Principal component analysis (PCA) was then performed on the SVG matrix (chrysalis.pca), followed by archetypal analysis (chrysalis.aa), number of tissue compartments (13) was selected using the elbow method on the reconstruction error curve. Two compartments were excluded from the analysis after initial examination of the gene contributions, as these had a very low number of positively weighted genes, most likely related to technician artefacts from the data. This was further verified by inspecting the spatial localisation of these compartments.

For the human samples, integration was performed with Scanorama [107] after normalisation and log-transformation due to large interpatient heterogeneity that is better corrected with a graph-based matching algorithm by preserving local structures instead of forcing sample alignment on a global scale. We ran chrysalis.aa with n_archetypes=20 based on the reconstruction error heuristic.

Tissue compartments were annotated by examining their top contributing genes, the spatial localisation of the compartments, the corresponding H&E tiles (**Supplementary Fig.25**), and their correlation with the cell density data inferred by cell2location. The latter was calculated using pairwise Pearson correlation between inferred cell-type density and tissue compartment scores. Sample level compartment proportions were calculated by summing compartment scores across all capture spots for each tissue compartment and normalised by the total sum of all compartment scores.

Comparison between human and mouse cellular niches was conducted by first mapping the mouse genes to the human orthologues using the human-mouse mapping table from BioMart, followed by calculating the intersection of SVGs across species. Pairwise cosine similarity was calculated using the Scikit-learn [108] implementation (metrics.pairwise.cosine_similarity).

#### Functional characterisation of mammary tumours

Pathway activities were inferred using PROGENy [63, 101], a curated collection of pathways, via decoupleR. The activity of 14 pathways across all capture spots was inferred with a multivariate linear model (decoupler.run_mlm), using the top 500 genes ranked by pathway responses in mice.

Hallmarks of Cancer gene sets were acquired from the original publication [66]. Mouse gene sets were constructed using the human-mouse orthologue mapping table queried from BioMart. Gene set enrichment scores were calculated across each capture spot using SCANPY’s gene set scoring function (sc.tl.score_genes). The same procedure was followed to calculate SASP [69], adaptive mutability gene sets [71], and the cell cycle activity gene set [109]. The cell cycle activity gene set was constructed by combining common gene sets overexpressed in specific cell cycle phases, providing an approximate measure of overall cell cycle activity.

Hallmarks MsigDB [72] collection gene set scores were calculated using ORA via decoupleR. Significant gene sets were selected if the corresponding *P* values were less than 0.05 in more than 90% of the total number of capture spots.

Enrichment of GO terms [73] in the cellular niches was calculated with ORA using the gene expression programs of each niche. These gene sets were defined by including genes whose weight exceeded a threshold set at the mean plus two standard deviations (cellular niche signatures can be found in **Supplementary Table 4**). Enrichment scores were scaled, and the top three terms were subsequently selected for each cellular niche and visualised as a heatmap.

Mean pathway activities, Hallmarks of Cancer, SASP, adaptive mutability, Hallmarks, and GO gene sets were calculated by averaging the activity scores of all capture spots across each condition and visualised as heatmaps. To infer associations with cellular niches, pairwise Pearson correlation was calculated with the tissue compartment scores.

Pseudo-bulk DGEA-based functional analysis on the mouse EMT and proliferating niche dominant spots, as well as the human primary and residual tumour-containing niches, were performed by first selecting capture spots that have a compartment score of at least 0.8. DGEA was performed as described in the *Pseudo-bulk DGEA* section with filtering parameters of (mouse: min_count=10, min_total_count=200, human: min_count=300, min_total_count=200). The resulting gene-level statistics were further used to perform enrichment analysis. Transcription factor activity was calculated with decoupleR with a univariate linear model (decupler.run_ulm) using the CollecTRI [110] resource containing a curated list of transcription factors and target genes. Pathway activities were calculated using PROGENy (decoupler.run_mlm). Enrichment of Hallmarks gene sets were calculated using ORA (decoupler.get_ora_df) on the differentially expressed genes (*P*_*adjusted*_ > 0.05, log2FC < 0.5).

#### CNV inference

To predict CNV in the ST samples, inferCNVpy, a Python implementation of the inferCNV [111] package was used. The genomic positions of genes were identified using the mm10 mouse genome. CNV inference was performed on the normalised and log-transformed samples using infercnvpy.tl.infercnv with the following parameters: window_size=100, step=10, chunksize=100. Capture spots annotated as epithelial cells from the healthy control samples, immune cells, muscle, and stroma were used as a reference. CNV scores were grouped based on disease condition (primary, residual, recurrent) and parental tumour lines. To generate CNV clusters, the generic clustering workflow was performed with inferCNVpy using the default parameters (PCA, neighbourhood graph computation, Leiden clustering). CNV cluster composition violin plots (tissue compartments, inferred cell type proportions) were generated with scanpy.pl.stacked_violin.

#### Cell type co-localisation

Cell type compartments were identified in the ST samples using the cell type densities inferred with cell2location by calculating the Pearson correlation between cell types across conditions. Similarly to other heatmaps presented in this work, cell-cell colocalisation clusters were defined using hierarchical clustering with SciPy’s [112] cluster.hierarchy.linkage function with the Ward variance minimisation algorithm using a height threshold of 3.5. The temporal flow of cell-cell colocalisation clusters was visualised with a Sankey diagram.

#### IMC data co-registration and processing

IMC data acquisitions were co-registered with ST data obtained from serial tissue sections (**Supplementary Table 11**). For each sample pair (*n*=7), manual landmark points (20-40) were selected on the Visium H&E section and the optical microscopy panorama image that was captured during the IMC protocol prior to tissue ablation. These landmark point pairs were used to estimate an affine transformation matrix via the RANSAC algorithm (cv2.estimateAffine2D, parameters: cv2.RANSAC, ransacReprojThreshold=100). The resulting transformation matrix was subsequently applied to map the coordinates of Visium capture spots to the coordinate system of the IMC dataset.

To construct multimodal datasets from these two modalities, we leveraged the recently developed SpatialData [113] Python library that can effectively handle various elements (pixel images, polygon shapes, pixel masks, points), tabular data, coordinate systems, and transformations between coordinate systems. 26 channels corresponding to the detected heavy metal isotopes were extracted from the MCD files (see **Supplementary Table 12** for antibody mapping to isotope channels) and stored as SpatialData objects along with the Visium capture spots.

IMC image processing was performed by performing hot pixel removal using a median filter. A 3 × 3 sliding window was applied across the ROIs, and any pixel with a value exceeding the median of the window plus a user-defined threshold (default: 100) was replaced with the median. This method was designed to be permissive, removing only clear outlier pixels while preserving the integrity of the surrounding data. This was followed by segmenting individual cells using Mesmer [114], a deep learning-based algorithm specifically designed for multiplexed tissue imaging. Mesmer predicted segmentation masks using two input channels, one assigned to cell nuclei (constructed by aggregating the following channels: H3/^176^Yb, DNA1/^191^Ir, DNA2/^193^Ir) and one to cytoplasms (vimentin/^150^Nd, e-cadherin/^158^Gd, pan-actin/^175^Lu). Cell contour polygons were created with the marching squares algorithm implemented in the scikitimage [115] Python library (skimage.measure.find_contours, parameters: level=0.5).

Cell phenotyping was performed with Pixie [74], a pixel-clustering pipeline designed for quantitative annotation of pixel-level features through unsupervised clustering. Pixie employs Self-organizing Map (SOM), an unsupervised clustering algorithm, to identify pixel-level clusters, which are further refined through hierarchical clustering. These pixel clusters are then mapped to segmented cells, resulting in cell-level clusters that can be classified based on input channel intensities. To achieve this, the selected IMC channels containing cell type markers (**Supplementary Table 12**) were preprocessed by performing Gaussian blurring, normalisation of pixel values across channels, and empty pixel removal with create_pixel_matrix from ark.phenotyping.pixie_preprocessing. This was followed by SOM training (train_pixel_som from ark.phenotyping.pixel_som_clustering), clustering (cluster_pixels from ark.phenotyping.pixel_som_clustering), and hierarchical clustering (pixel_consensus_cluster from ark.phenotyping.pixel_meta_clustering) to generate the pixel clusters. Cell-level clustering was performed by cell SOM training (train_cell_som from ark.phenotyping.cell_som_clustering), clustering (cluster_cells from ark.phenotyping.cell_som_clustering), and hierarchical clustering (cell_consensus_cluster from ark.phenotyping.cell_meta_clustering). Cell-level clusters were manually refined to produce the final cell-type labels.

#### IMC data analysis

IMC measurements produce highly consistent raw pseudocount profiles across ROIs. Nevertheless, there is still detectable variation between the distributions of the measured heavy metal isotope intensities. Pt distributions across ROIs were equalised to produce accurate readings of intracellular Pt intensities. A cell neighbourhood graph was constructed with Squidpy’s [116] gr.spatial_neighbors (radius=(0, 40), coord_type=“generic”, delaunay=False) function based on the spatial proximity of 40 µm jointly across all ROIs for each sample. Next, a border region was defined for each ROI by shrinking its bounding box by a 25 µm margin, ensuring that the correction focused on cells located near the edges of the ROI. Cells within this margin were then used to evaluate differences in Pt intensity between neighbouring ROIs. The Pt intensity of all cells within each ROI was compared to the average Pt intensity of cells in neighbouring ROIs, with differences weighted by the number of neighbouring cells. A normalisation factor for each ROI was calculated by comparing its weighted average intensity to the global mean, and these factors were applied to scale Pt intensity values.

Intracellular Pt concentration was calculated by summing up the corrected raw counts across all Pt channels within the cell segmentation masks and dividing by the area of the cell. Log transformation was performed with scanpy.pp.log1p. tumour cells were classified based on Pt concentration as follows: Pt_low_ cells were those below the 15th percentile, Pt_mod_ cells were between the 15th and 85th percentiles, and Pt_high_ cells were above the 85th percentile of the log-transformed Pt concentration distribution. GMM-based cutoff for perinecrotic tumour cells was determined using mixture. GaussianMixture from the Scikit-learn Python package with two components.

Pt concentration was averaged per cell type across conditions and visualised as a heatmap. Ki-67 high and Ki-67 low tumour cells were classified by a cutoff determined with a GMM on the log-transformed intracellular Ki-67 concentration.

To correlate the cell types inferred by cell2location with intracellular Pt concentrations, the centroids of mapped Visium spots were expanded into discs with a diameter of 55 µm. The mean Pt concentration was then calculated across all cells within each disc. Pairwise Pearson correlation was subsequently calculated between the mean Pt concentration and the inferred cell type densities across conditions.

Contributions of molecular features in explaining intracellular Pt content were calculated using MISTy [79] via LIANA+ [117]. MISTy is an explainable machine learning framework that models the interactions between features in the spatial context. The contribution of features is analysed to highlight the relevance of potential interaction sources that originate from distinct spatial contexts, referred to as views. Custom views were constructed for the predictor variable sets (cell2location-inferred cell types, PROGENy pathway activities, Chrysalis compartments, cell cycle activities) using liana.ut.spatial_neighbors with the following parameters: bandwidth=500, cutoff=0.1. Log-transformed Pt concentration was set as a target variable and constructed as an intraview. Following that, a random forest model was trained with the bypass_intra parameter set to True. Coefficients of determination of the predictor variable sets were subsequently visualised as stacked bar plots. The importance of individual features was scaled and visualised as rank plots.

#### Niche signatures in human breast cancer

EMT and proliferating tumour niche signatures were preprocessed by mapping the mouse gene signatures to human orthologues and filtering out genes that are not expressed in the human samples. Expression scores of these gene sets were calculated with scanpy.tl.score_genes in the human ST samples. Score distributions were assessed in selected capture spots annotated as tumour by pathologists.

#### Survival analysis

SCAN-B [82] and TCGA-BRCA [83] breast cancer patient cohorts were first filtered according to their receptor expression status. We used the provided metadata in combination with GMMs to create cutoffs for the expression of ER, PR and HER genes to identify TNBC patients.

In the SCAN-B cohort, this filtering resulted in 315 TNBC patients. We stratified these patients according to their EMT expression signature calculated with scanpy.tl.score_genes to EMT_High_ (*n*=157) and EMT_Low_ (*n*=158) by a cutoff set at the median of the EMT signature distribution. Survival analysis was performed with the lifelines [118] Python package. The Kaplan-Meier estimator (lifelines.KaplanMeierFitter) was used to estimate the survival functions, followed by a log-rank test (lifelines.statistics.logrank_test) to assess statistical differences between survival distributions, resulting in a *P* value < 0.005 between the examined groups.

In the TCGA-BRCA cohort, 163 TNBC patients were identified. Patients were stratified based on a combination of EMT and proliferating niche expression signatures. For each signature, a gene expression matrix was constructed using the genes within the respective signatures. PCA (sc.pl.pca) was performed on each matrix, followed by K-means clustering (sklearn.cluster.KMeans, n_clusters=2). Patients were stratified based on the combination of the two EMT and proliferating clusters, resulting in EMT_High_-Proliferating_Low_, EMT_Low_-Proliferating_High_, EMT_Low_-Proliferating_Low_, and EMTHgh-Proliferating_High_ groups. The Kaplan-Meier estimator (lifelines.KaplanMeierFitter) was used to estimate the survival functions between EMT_High_-Proliferating_Low_ (*n*=19), and EMT_Low_-Proliferating_High_ (*n*=34) patient groups, followed by a log-rank test (lifelines.statistics.logrank_test) to assess statistical differences between survival distributions, resulting in a *P* value < 0.005 between the examined groups.

### Statistics and reproducibility

Statistical methods for each analysis are detailed in their respective sections. Pearson correlation coefficients were computed with the pearsonr function from SciPy’s stats module. Visualisations, including scatterplots, box plots, violin plots, bar plots, KDE plots, heatmaps, and line plots, were created using matplotlib [119], SCANPY, ausankey, lifelines, Chrysalis, and seaborn [120]. Investigators were blinded to treatment status during the experiments but were unblinded during the analysis phase.

## Data availability

The raw RNA sequencing data will be deposited in the Gene Expression Omnibus (GEO) database and made available upon publication. All processed ST (10x Visium), scRNA-seq (10x Chromium), and IMC (Hyperion) data, along with supplementary data required to replicate the analyses in this study, are available in the following Zenodo repositories: ST (https://doi.org/10.5281/zenodo.15102983), scRNA-seq (https://doi.org/10.5281/zenodo.15103411), and IMC (co-registered with Visium) (https://doi.org/10.5281/zenodo.15096025). The uploaded data includes count matrices (Space Ranger and Cell Ranger outputs), AnnData objects (containing cell type deconvolution, tissue compartment inference results, histopathological annotations, and gene set signatures), and raw and processed IMC acquisitions stored in SpatialData Zarr archives.

## Code availability

The code used for the data analysis presented in this article is available on GitHub (https://github.com/rottenberglab/residual-disease).

## Acknowledgements

We thank Carmen Widmer, Maurits Roorda, Diego Dibitetto and Chang He for the critical reading of the manuscript. We acknowledge the University of Geneva for providing the human samples used in this publication and sincerely thank all patient donors. We would like to thank the Stroka lab at the University of Bern for the mouse antibody panel, and the IMC Platform in Bern for the IMC acquisitions. We are grateful for the support of the Flow Cytometry and Cell Sorting Facility at the University of Bern with the acquisition of the scRNA-seq samples. We thank the NGS platform at the University of Bern for the sequencing of the scRNA-seq and ST samples. We thank Cédric Walker for the alignment of the scRNA-seq data and Michael Berger for his help with the RNA velocity analysis. We thank Chang He for his help with the histopathological examination. This work was supported by the Swiss National Science Foundation (320030M_219453 to S.R., 323530_207032 to M.D.), the Swiss Cancer League (KFS-5519-02-2022 to S.R.), the ISREC foundation, and the European Research Council (ERC-2019-AdG-883877 to S.R.).

## Author Information

### Contributions

D.T., M.D., and S.R. conceptualised and designed the study. M.D. performed mouse experiments with the technical support of M.S., G.L., and F.S.. M.D. selected tissue of interest for scRNA-seq, ST, and IMC and performed histopathological annotations with the technical support of A.C.. D.T. performed scRNA-seq, ST, and IMC data analysis. I.L. provided human breast cancer samples. A.C. performed tissue sectioning and ST experiments. D.T. and M.D. wrote the manuscript and designed the figures with input from S.R. and A.V.. S.R. and A.V. supervised the project. All authors read and approved the manuscript.

### Corresponding author

Correspondence to Sven Rotten-berg.

## Bibliography

1. Rebecca L. Siegel, Kimberly D. Miller, Nikita Sandeep Wagle, and Ahmedin Jemal. Cancer statistics, 2023. CA: A Cancer Journal for Clinicians, 73(1):17–48, 2023. ISSN 0007-9235. doi: 10.3322/caac.21763.

2. Sylvia Annabel Dass, Kim Liu Tan, Rehasri Selva Rajan, Noor Fatmawati Mokhtar, Elis Rosliza Mohd Adzmi, Wan Faiziah Wan Abdul Rahman, Tengku Ahmad Damitri Al-Astani Tengku Din, and Venugopal Balakrishnan. Triple Negative Breast Cancer: A Review of Present and Future Diagnostic Modalities. Medicina, 57(1):62, 2021. ISSN 1010-660X. doi: 10.3390/medicina57010062.

3. Fatemeh Derakhshan and Jorge S. Reis-Filho. Pathogenesis of Triple-Negative Breast Cancer. Annual Review of Pathology: Mechanisms of Disease, 17(1):181–204, 2022. ISSN 1553-4006. doi: 10.1146/annurev-pathol-042420-093238.

4. Gunter von Minckwitz, Andreas Schneeweiss, Sibylle Loibl, Christoph Salat, Carsten Denkert, Mahdi Rezai, Jens U Blohmer, Christian Jackisch, Stefan Paepke, Bernd Gerber, Dirk M Zahm, Sherko Kümmel, Holger Eidtmann, Peter Klare, Jens Huober, Serban Costa, Hans Tesch, Claus Hanusch, Jörn Hilfrich, Fariba Khandan, Peter A Fasching, Bruno V Sinn, Knut Engels, Keyur Mehta, Valentina Nekljudova, and Michael Untch. Neoadjuvant carboplatin in patients with triple-negative and HER2-positive early breast cancer (Gepar-Sixto; GBG 66): a randomised phase 2 trial. The Lancet Oncology, 15(7):747–756, 2014. ISSN 1470-2045. doi: 10.1016/s1470-2045(14)70160-3.

5. William M. Sikov, Donald A. Berry, Charles M. Perou, Baljit Singh, Constance T. Cirrincione, Sara M. Tolaney, Charles S. Kuzma, Timothy J. Pluard, George Somlo, Elisa R. Port, Mehra Golshan, Jennifer R. Bellon, Deborah Collyar, Olwen M. Hahn, Lisa A. Carey, Clifford A. Hudis, and Eric P. Winer. Impact of the Addition of Carboplatin and/or Bevacizumab to Neoadjuvant Once-per-Week Paclitaxel Followed by Dose-Dense Doxorubicin and Cyclophosphamide on Pathologic Complete Response Rates in Stage II to III Triple-Negative Breast Cancer: CALGB 40603 (Alliance). Journal of Clinical Oncology, 33(1): 13–21, 2014. ISSN 0732-183X. doi: 10.1200/jco.2014.57.0572.

6. Sibylle Loibl, Joyce O’Shaughnessy, Michael Untch, William M Sikov, Hope S Rugo, Mark D McKee, Jens Huober, Mehra Golshan, Gunter von Minckwitz, David Maag, Danielle Sullivan, Norman Wolmark, Kristi McIntyre, Jose J Ponce Lorenzo, Otto Metzger Filho, Priya Rastogi, W Fraser Symmans, Xuan Liu, and Charles E Geyer. Addition of the PARP inhibitor veliparib plus carboplatin or carboplatin alone to standard neoadjuvant chemotherapy in triple-negative breast cancer (BrighTNess): a randomised, phase 3 trial. The Lancet Oncology, 19(4):497–509, 2018. ISSN 1470-2045. doi: 10.1016/s1470-2045(18)30111-6.

7. Peter Schmid, Sylvia Adams, Hope S Rugo, Andreas Schneeweiss, Carlos H Barrios, Hiroji Iwata, Véronique Diéras, Roberto Hegg, Seock-Ah Im, Gail Shaw Wright, Volkmar Henschel, Luciana Molinero, Stephen Y Chui, Roel Funke, Amreen Husain, Eric P Winer, Sherene Loi, Leisha A Emens, and IMpassion130 Trial Investigators. Atezolizumab and Nab-Paclitaxel in Advanced Triple-Negative Breast Cancer. New England Journal of Medicine, 379(22):2108–2121, 2018. ISSN 0028-4793. doi: 10.1056/nejmoa1809615.

8. Peter Schmid, Javier Cortes, Lajos Pusztai, Heather McArthur, Sherko Kümmel, Jonas Bergh, Carsten Denkert, Yeon Hee Park, Rina Hui, Nadia Harbeck, Masato Takahashi, Theodoros Foukakis, Peter A Fasching, Fatima Cardoso, Michael Untch, Liyi Jia, Vassiliki Karantza, Jing Zhao, Gursel Aktan, Rebecca Dent, Joyce O’Shaughnessy, and KEYNOTE-522 Investigators. Pembrolizumab for Early Triple-Negative Breast Cancer. New England Journal of Medicine, 382(9):810–821, 2020. ISSN 0028-4793. doi:10.1056/nejmoa1910549.

9. Julia Foldi, Andrea Silber, Emily Reisenbichler, Kamaljeet Singh, Neal Fischbach, Justin Persico, Kerin Adelson, Anamika Katoch, Nina Horowitz, Donald Lannin, Anees Chagpar, Tristen Park, Michal Marczyk, Courtney Frederick, Trisha Burrello, Eiman Ibrahim, Tao Qing, Yalai Bai, Kim Blenman, David L. Rimm, and Lajos Pusztai. Neoadjuvant durvalumab plus weekly nab-paclitaxel and dose-dense doxorubicin/cyclophosphamide in triple-negative breast cancer. npj Breast Cancer, 7(1):9, 2021. ISSN 2374-4677. doi:10.1038/s41523-021-00219-7.

10. L. Pusztai, C. Denkert, J. O’Shaughnessy, J. Cortes, R. Dent, H. McArthur, S. Kümmel, J. Bergh, Y.H. Park, R. Hui, N. Harbeck, M. Takahashi, M. Untch, P.A. Fasching, F. Cardoso, Y. Zhu, W. Pan, K. Tryfonidis, and P. Schmid. Event-free survival by residual cancer burden with pembrolizumab in early-stage TNBC: exploratory analysis from KEYNOTE-522. Annals of Oncology, 35(5):429–436, 2024. ISSN 0923-7534. doi:10.1016/j.annonc.2024.02.002.

11. Shiyun Cui, Weici Liu, Wenxiang Wang, Keyan Miao, and Xiaoxiang Guan. Advances in the diagnosis and prognosis of minimal residual lesions of breast cancer. Pathology - Research and Practice, 245:154428, 2023. ISSN 0344-0338. doi: 10.1016/j.prp.2023.154428.

12. Piet Borst. Cancer drug pan-resistance: pumps, cancer stem cells, quiescence, epithelial to mesenchymal transition, blocked cell death pathways, persisters or what? Open Biology, 2(5):120066, 2012. doi: 10.1098/rsob.120066.

13. Michael Ramirez, Satwik Rajaram, Robert J. Steininger, Daria Osipchuk, Maike A. Roth, Leanna S. Morinishi, Louise Evans, Weiyue Ji, Chien-Hsiang Hsu, Kevin Thurley, Shuguang Wei, Anwu Zhou, Prasad R. Koduru, Bruce A. Posner, Lani F. Wu, and Steven J. Altschuler. Diverse drug-resistance mechanisms can emerge from drug-tolerant cancer persister cells. Nature Communications, 7(1):10690, 2016. doi: 10.1038/ncomms10690.

14. Giulia De Conti, Matheus Henrique Dias, and René Bernards. Fighting Drug Resistance through the Targeting of Drug-Tolerant Persister Cells. Cancers, 13(5):1118, 2021. ISSN 2072-6694. doi: 10.3390/cancers13051118.

15. Morgane Decollogny and Sven Rottenberg. Persisting cancer cells are different from bacterial persisters. Trends in Cancer, 2024. ISSN 2405-8033. doi: 10.1016/j.trecan.2024.02.002.

16. Marina Pajic, Sohvi Blatter, Charlotte Guyader, Maaike Gonggrijp, Ariena Kersbergen, Aslι Küçükosmanoėlu, Wendy Sol, Rinske Drost, Jos Jonkers, Piet Borst, and Sven Rottenberg. Selected Alkylating Agents Can Overcome Drug Tolerance of G0-like Tumor Cells and Eradicate BRCA1-Deficient Mammary Tumors in Mice. Clinical Cancer Research, 23 (22):7020–7033, 2017. ISSN 1078-0432. doi: 10.1158/1078-0432.ccr-17-1279.

17. María Soledad Sosa, Paloma Bragado, and Julio A. Aguirre-Ghiso. Mechanisms of disseminated cancer cell dormancy: an awakening field. Nature Reviews Cancer, 14(9): 611–622, 2014. ISSN 1474-175X. doi: 10.1038/nrc3793.

18. Jason R. Ruth, Dhruv K. Pant, Tien-chi Pan, Hans E. Seidel, Sanjeethan C. Baksh, Blaine A. Keister, Rita Singh, Christopher J. Sterner, Suzanne J. Bakewell, Susan E. Moody, George K. Belka, and Lewis A. Chodosh. Cellular dormancy in minimal residual disease following targeted therapy. Breast Cancer Research, 23(1):63, 2021. ISSN 1465-5411. doi: 10.1186/s13058-021-01416-9.

19. Erica Dalla, Amulya Sreekumar, Julio A. Aguirre-Ghiso, and Lewis A. Chodosh. Dormancy in Breast Cancer. Cold Spring Harbor Perspectives in Medicine, 13(11):a041331, 2023. ISSN 2157-1422. doi: 10.1101/cshperspect.a041331.

20. Yaara Oren, Michael Tsabar, Michael S. Cuoco, Liat Amir-Zilberstein, Heidie F. Cabanos, Jan-Christian Hütter, Bomiao Hu, Pratiksha I. Thakore, Marcin Tabaka, Charles P. Fulco, William Colgan, Brandon M. Cuevas, Sara A. Hurvitz, Dennis J. Slamon, Amy Deik, Kerry A. Pierce, Clary Clish, Aaron N. Hata, Elma Zaganjor, Galit Lahav, Katerina Politi, Joan S. Brugge, and Aviv Regev. Cycling cancer persister cells arise from lineages with distinct programs. Nature, 596(7873):576–582, 2021. ISSN 0028-0836. doi: 10.1038/s41586-021-03796-6.

21. Eugen Dhimolea, Ricardo de Matos Simoes, Dhvanir Kansara, Aziz Al’Khafaji, Juliette Bouyssou, Xiang Weng, Shruti Sharma, Joseline Raja, Pallavi Awate, Ryosuke Shirasaki, Huihui Tang, Brian J. Glassner, Zhiyi Liu, Dong Gao, Jordan Bryan, Samantha Bender, Jennifer Roth, Michal Scheffer, Rinath Jeselsohn, Nathanael S. Gray, Irene Georgakoudi, Francisca Vazquez, Aviad Tsherniak, Yu Chen, Alana Welm, Cihangir Duy, Ari Melnick, Boris Bartholdy, Myles Brown, Aedin C. Culhane, and Constantine S. Mitsiades. An Embryonic Diapause-like Adaptation with Suppressed Myc Activity Enables Tumor Treatment Persistence. Cancer Cell, 39(2):240–256.e11, 2021. ISSN 1535-6108. doi: 10.1016/j.ccell.2020.12.002.

22. Sumaiyah K. Rehman, Jennifer Haynes, Evelyne Collignon, Kevin R. Brown, Yadong Wang, Allison M.L. Nixon, Jeffrey P. Bruce, Jeffrey A. Wintersinger, Arvind Singh Mer, Edwyn B.L. Lo, Cherry Leung, Evelyne Lima-Fernandes, Nicholas M. Pedley, Fraser Soares, Sophie McGibbon, Housheng Hansen He, Aaron Pollet, Trevor J. Pugh, Benjamin Haibe-Kains, Quaid Morris, Miguel Ramalho-Santos, Sidhartha Goyal, Jason Moffat, and Catherine A. O’Brien. Colorectal Cancer Cells Enter a Diapause-like DTP State to Survive Chemotherapy. Cell, 184(1):226–242.e21, 2021. ISSN 0092-8674. doi: 10.1016/j.cell.2020.11.018.

23. Sreenath V. Sharma, Diana Y. Lee, Bihua Li, Margaret P. Quinlan, Fumiyuki Takahashi, Shyamala Maheswaran, Ultan McDermott, Nancy Azizian, Lee Zou, Michael A. Fischbach, Kwok-Kin Wong, Kathleyn Brandstetter, Ben Wittner, Sridhar Ramaswamy, Marie Classon, and Jeff Settleman. A Chromatin-Mediated Reversible Drug-Tolerant State in Cancer Cell Subpopulations. Cell, 141(1):69–80, 2010. ISSN 0092-8674. doi: 10.1016/j.cell.2010.02.027.

24. Gloria V. Echeverria, Zhongqi Ge, Sahil Seth, Xiaomei Zhang, Sabrina Jeter-Jones, Xinhui Zhou, Shirong Cai, Yizheng Tu, Aaron McCoy, Michael Peoples, Yuting Sun, Huan Qiu, Qing Chang, Christopher Bristow, Alessandro Carugo, Jiansu Shao, Xiaoyan Ma, Angela Harris, Prabhjot Mundi, Rosanna Lau, Vandhana Ramamoorthy, Yun Wu, Mariano J. Alvarez, Andrea Califano, Stacy L. Moulder, William F. Symmans, Joseph R. Marszalek, Timothy P. Heffernan, Jeffrey T. Chang, and Helen Piwnica-Worms. Resistance to neoadjuvant chemotherapy in triple-negative breast cancer mediated by a reversible drug-tolerant state. Science Translational Medicine, 11(488), 2019. ISSN 1946-6234. doi: 10.1126/scitranslmed.aav0936.

25. Nilanjana Mani, Ankita Daiya, Rajdeep Chowdhury, Sudeshna Mukherjee, and Shibasish Chowdhury. Epigenetic adaptations in drug-tolerant tumor cells. Advances in Cancer Research, 158:293–335, 2023. ISSN 0065-230X. doi: 10.1016/bs.acr.2022.12.006.

26. Matthew J. Hangauer, Vasanthi S. Viswanathan, Matthew J. Ryan, Dhruv Bole, John K. Eaton, Alexandre Matov, Jacqueline Galeas, Harshil D. Dhruv, Michael E. Berens, Stuart L. Schreiber, Frank McCormick, and Michael T. McManus. Drug-tolerant persister cancer cells are vulnerable to GPX4 inhibition. Nature, 551(7679):247–250, 2017. ISSN 0028-0836. doi: 10.1038/nature24297.

27. Kristina M. Havas, Vladislava Milchevskaya, Ksenija Radic, Ashna Alladin, Eleni Kafkia, Marta Garcia, Jens Stolte, Bernd Klaus, Nicole Rotmensz, Toby J. Gibson, Barbara Burwinkel, Andreas Schneeweiss, Giancarlo Pruneri, Kiran R. Patil, Rocio Sotillo, and Martin Jechlinger. Metabolic shifts in residual breast cancer drive tumor recurrence. Journal of Clinical Investigation, 127(6):2091–2105, 2017. ISSN 0021-9738. doi: 10.1172/jci89914.

28. Laura Vera-Ramirez, Suman K. Vodnala, Ryan Nini, Kent W. Hunter, and Jeffrey E. Green. Autophagy promotes the survival of dormant breast cancer cells and metastatic tumour recurrence. Nature Communications, 9(1):1944, 2018. doi: 10.1038/s41467-018-04070-6.

29. Enakshi D. Sunassee, Riley J. Deutsch, Victoria W. D’Agostino, Pol Castellano-Escuder, Elizabeth A. Siebeneck, Olga Ilkayeva, Brian T. Crouch, Megan C. Madonna, Jeffrey Everitt, James V. Alvarez, Gregory M. Palmer, Matthew D. Hirschey, and Nirmala Ramanujam. Optical imaging reveals chemotherapy-induced metabolic reprogramming of residual disease and recurrence. Science Advances, 10(14):eadj7540, 2024. doi: 10.1126/sciadv.adj7540.

30. Guangyu Wang, Dandan Xu, Zicheng Zhang, Xinhui Li, Jiaqi Shi, Jie Sun, Huan-Zhong Liu, Xiaobo Li, Meng Zhou, and Tongsen Zheng. The pan-cancer landscape of crosstalk between epithelial-mesenchymal transition and immune evasion relevant to prognosis and immunotherapy response. npj Precision Oncology, 5(1):56, 2021. ISSN 2397-768X. doi: 10.1038/s41698-021-00200-4.

31. Sven Rottenberg, Anders O. H. Nygren, Marina Pajic, Fijs W. B. van Leeuwen, Ingrid van der Heijden, Koen van de Wetering, Xiaoling Liu, Karin E. de Visser, Kenneth G. Gilhuijs, Olaf van Tellingen, Jan P. Schouten, Jos Jonkers, and Piet Borst. Selective induction of chemotherapy resistance of mammary tumors in a conditional mouse model for hereditary breast cancer. Proceedings of the National Academy of Sciences, 104(29): 12117–12122, 2007. ISSN 0027-8424. doi: 10.1073/pnas.0702955104.

32. Sven Rottenberg, Janneke E. Jaspers, Ariena Kersbergen, Eline van der Burg, Anders O. H. Nygren, Serge A. L. Zander, Patrick W. B. Derksen, Michiel de Bruin, John Zevenhoven, Alan Lau, Robert Boulter, Aaron Cranston, Mark J. O’Connor, Niall M. B. Martin, Piet Borst, and Jos Jonkers. High sensitivity of BRCA1-deficient mammary tumors to the PARP inhibitor AZD2281 alone and in combination with platinum drugs. Proceedings of the National Academy of Sciences, 105(44):17079–17084, 2008. ISSN 0027-8424. doi: 10.1073/pnas.0806092105.

33. Xiaoling Liu, Henne Holstege, Hanneke van der Gulden, Marcelle Treur-Mulder, John Zevenhoven, Arno Velds, Ron M. Kerkhoven, Martin H. van Vliet, Lodewyk F. A. Wessels, Johannes L. Peterse, Anton Berns, and Jos Jonkers. Somatic loss of BRCA1 and p53 in mice induces mammary tumors with features of human BRCA1-mutated basal-like breast cancer. Proceedings of the National Academy of Sciences, 104(29):12111–12116, 2007. ISSN 0027-8424. doi: 10.1073/pnas.0702969104.

34. Kohei Saeki, Gregory Chang, Noriko Kanaya, Xiwei Wu, Jinhui Wang, Lauren Bernal, Desiree Ha, Susan L. Neuhausen, and Shiuan Chen. Mammary cell gene expression atlas links epithelial cell remodeling events to breast carcinogenesis. Communications Biology, 4(1):660, 2021. doi: 10.1038/s42003-021-02201-2.

35. Carman Man-Chung Li, Hana Shapiro, Christina Tsiobikas, Laura M. Selfors, Huidong Chen, Jennifer Rosenbluth, Kaitlin Moore, Kushali P. Gupta, G. Kenneth Gray, Yaara Oren, Michael J. Steinbaugh, Jennifer L. Guerriero, Luca Pinello, Aviv Regev, and Joan S. Brugge. Aging-Associated Alterations in Mammary Epithelia and Stroma Revealed by Single-Cell RNA Sequencing. Cell Reports, 33(13):108566, 2020. ISSN 2211-1247. doi:10.1016/j.celrep.2020.108566.

36. Jingjing Qi, Hongxiang Sun, Yao Zhang, Zhengting Wang, Zhenzhen Xun, Ziyi Li, Xinyu Ding, Rujuan Bao, Liwen Hong, Wenqing Jia, Fei Fang, Hongzhi Liu, Lei Chen, Jie Zhong, Duowu Zou, Lianxin Liu, Leng Han, Florent Ginhoux, Yingbin Liu, Youqiong Ye, and Bing Su. Single-cell and spatial analysis reveal interaction of FAP+ fibroblasts and SPP1+ macrophages in colorectal cancer. Nature Communications, 13(1):1742, 2022. doi: 10.1038/s41467-022-29366-6.

37. Katja Zappe, Antonio Kopic, Alexandra Scheichel, Ann-Katrin Schier, Lukas Emanuel Schmidt, Yasmin Borutzki, Heidi Miedl, Martin Schreiber, Theresa Mendrina, Christine Pirker, Georg Pfeiler, Stefan Hacker, Werner Haslik, Dietmar Pils, Andrea Bileck, Christopher Gerner, Samuel Meier-Menches, Petra Heffeter, and Margit Cichna-Markl. Aberrant DNA Methylation, Expression, and Occurrence of Transcript Variants of the ABC Transporter ABCA7 in Breast Cancer. Cells, 12(11):1462, 2023. doi: 10.3390/cells12111462.

38. Samantha D. Praktiknjo, Benedikt Obermayer, Qionghua Zhu, Liang Fang, Haiyue Liu, Hazel Quinn, Marlon Stoeckius, Christine Kocks, Walter Birchmeier, and Nikolaus Rajewsky. Tracing tumorigenesis in a solid tumor model at single-cell resolution. Nature Communications, 11(1):991, 2020. doi: 10.1038/s41467-020-14777-0.

39. Rui Tian, Xiangsheng Zuo, Jonathan Jaoude, Fei Mao, Jennifer Colby, and Imad Shureiqi. ALOX15 as a suppressor of inflammation and cancer: Lost in the link. Prostaglandins & Other Lipid Mediators, 132:77–83, 2017. ISSN 1098-8823. doi: 10.1016/j.prostaglandins.2017.01.002.

40. Maria del Mar Maldonado, Jeffrey Schlom, and Duane H. Hamilton. Blockade of tumor-derived colony-stimulating factor 1 (CSF1) promotes an immune-permissive tumor microenvironment. Cancer Immunology, Immunotherapy, 72(10):3349–3362, 2023. ISSN 0340-7004. doi: 10.1007/s00262-023-03496-2.

41. Lili Yi, Yongqiang Lei, Fengjiao Yuan, Conghui Tian, Jian Chai, and Mingliang Gu. NTN4 as a prognostic marker and a hallmark for immune infiltration in breast cancer. Scientific Reports, 12(1):10567, 2022. doi: 10.1038/s41598-022-14575-2.

42. Shusen Zhang, Zhigang Cai, and Hui Li. AHNAKs roles in physiology and malignant tumors. Frontiers in Oncology, 13:1258951, 2023. ISSN 2234-943X. doi: 10.3389/fonc.2023.1258951.

43. Chengheng Liao, Yang Zhang, Cheng Fan, Laura E. Herring, Juan Liu, Jason W. Locasale, Mamoru Takada, Jin Zhou, Giada Zurlo, Lianxin Hu, Jeremy M. Simon, Travis S. Ptacek, Victor G. Andrianov, Einars Loza, Yan Peng, Huanghe Yang, Charles M. Perou, and Qing Zhang. Identification of BBOX1 as a Therapeutic Target in Triple-Negative Breast Cancer. Cancer Discovery, 10(11):1706–1721, 2020. ISSN 2159-8274. doi: 10.1158/2159-8290.cd-20-0288.

44. Steve Leu. The role and regulation of Pnn in proliferative and non-dividing cells: Form embryogenesis to pathogenesis. Biochemical Pharmacology, 192:114672, 2021. ISSN 0006-2952. doi: 10.1016/j.bcp.2021.114672.

45. Lingling Li, Piao Li, Wei Zhang, Haiting Zhou, Ergang Guo, Guoqing Hu, and Linli Zhang. FERMT1 contributes to the migration and invasion of nasopharyngeal carcinoma through epithelial–mesenchymal transition and cell cycle arrest. Cancer Cell International, 22(1): 70, 2022. ISSN 1475-2867. doi: 10.1186/s12935-022-02494-1.

46. Ruiqi Liu, Xiaodong Liang, Haiwei Guo, Shuang Li, Weiping Yao, Chenfang Dong, Jiajun Wu, Yanwei Lu, Jianming Tang, and Haibo Zhang. STNM1 in human cancers: role, function and potential therapy sensitizer. Cellular Signalling, 109:110775, 2023. ISSN 0898-6568. doi: 10.1016/j.cellsig.2023.110775.

47. Ziqing Yang, Shaomin Zou, Yijing Zhang, Jieping Zhang, Peng Zhang, Lishi Xiao, Yunling Xie, Manqi Meng, Junyan Feng, Liang Kang, Mong-Hong Lee, and Lekun Fang. ACTL6A protects gastric cancer cells against ferroptosis through induction of glutathione synthesis. Nature Communications, 14(1):4193, 2023. doi: 10.1038/s41467-023-39901-8.

48. Salomé Adam Silvia Emma Rossi, Nathalie Moatti, Mara De Marco Zompit, Yibo Xue, Timothy F. Ng, Alejandro Álvarez Quilón, Jessica Desjardins, Vivek Bhaskaran, Giovanni Martino, Dheva Setiaputra, Sylvie M. Noordermeer, Toshiro K. Ohsumi, Nicole Hustedt, Rachel K. Szilard, Natasha Chaudhary, Meagan Munro, Artur Veloso, Henrique Melo, Shou Yun Yin, Robert Papp, Jordan T. F. Young, Michael Zinda, Manuel Stucki, and Daniel Durocher. The CIP2A–TOPBP1 axis safeguards chromosome stability and is a synthetic lethal target for BRCA-mutated cancer. Nature Cancer, 2(12):1357–1371, 2021. doi: 10.1038/s43018-021-00266-w.

49. Vitalii Kleshchevnikov, Artem Shmatko, Emma Dann, Alexander Aivazidis, Hamish W. King, Tong Li, Rasa Elmentaite, Artem Lomakin, Veronika Kedlian, Adam Gayoso, Mika Sarkin Jain, Jun Sung Park, Lauma Ramona, Elizabeth Tuck, Anna Arutyunyan, Roser Vento-Tormo, Moritz Gerstung, Louisa James, Oliver Stegle, and Omer Ali Bayraktar. Cell2location maps fine-grained cell types in spatial transcriptomics. Nature Biotechnology, pages 1–11, 2022. ISSN 1087-0156. doi: 10.1038/s41587-021-01139-4.

50. Taryn A. Donovan, Frances M. Moore, Christof A. Bertram, Richard Luong, Pompei Bolfa, Robert Klopfleisch, Harold Tvedten, Elisa N. Salas, Derick B. Whitley, Marc Aubreville, and Donald J. Meuten. Mitotic Figures—Normal, Atypical, and Imposters: A Guide to Identification. Veterinary Pathology, 58(2):243–257, 2021. ISSN 0300-9858. doi: 10.1177/0300985820980049.

51. Ning Wu, YouZhi Wang, KeKe Wang, BoQiang Zhong, YiHao Liao, JiaMing Liang, and Ning Jiang. Cathepsin K regulates the tumor growth and metastasis by IL-17/CTSK/EMT axis and mediates M2 macrophage polarization in castration-resistant prostate cancer. Cell Death & Disease, 13(9):813, 2022. doi: 10.1038/s41419-022-05215-8.

52. Shau-Hsuan Li, Hung-I Lu, Wan-Ting Huang, Yen-Hao Chen, Chien-Ming Lo, Ya-Chun Lan, Wei-Che Lin, Hsin-Ting Tsai, and Chang-Han Chen. An actin-binding protein ESPN is an independent prognosticator and regulates cell growth for esophageal squamous cell carcinoma. Cancer Cell International, 18(1):219, 2018. ISSN 1475-2867. doi: 10.1186/s12935-018-0713-x.

53. Fenglin Cai, Xiuding Yang, Gang Ma, Pengliang Wang, Mengmeng Zhang, Nannan Zhang, Rupeng Zhang, Han Liang, Yongzhan Nie, Cheng Dong, and Jingyu Deng. EGLN3 attenuates gastric cancer cell malignant characteristics by inhibiting JMJD8/NF-κB signalling activation independent of hydroxylase activity. British Journal of Cancer, 130(4):597–612, 2024. ISSN 0007-0920. doi: 10.1038/s41416-023-02546-x.

54. Manfei Si and Jinghe Lang. The roles of metallothioneins in carcinogenesis. Journal of Hematology & Oncology, 11(1):107, 2018. doi: 10.1186/s13045-018-0645-x.

55. Liping Shen, Shan Jiang, Yu Yang, Hongli Yang, Yanchun Fang, Meng Tang, Rangteng Zhu, Jiaqin Xu, and Hantao Jiang. Pan-cancer and single-cell analysis reveal the prognostic value and immune response of NQO1. Frontiers in Cell and Developmental Biology, 11:1174535, 2023. ISSN 2296-634X. doi: 10.3389/fcell.2023.1174535.

56. Demeter Túrós, Jelica Vasiljevic, Kerstin Hahn, Sven Rottenberg, and Alberto Valdeolivas. Chrysalis: decoding tissue compartments in spatial transcriptomics with archetypal analysis. Communications Biology, 7(1):1520, 2024. doi: 10.1038/s42003-024-07165-7.

57. Yan Tu, Cameron N. Johnstone, and Alastair G. Stewart. Annexin A1 influences in breast cancer: Controversies on contributions to tumour, host and immunoediting processes. Pharmacological Research, 119:278–288, 2017. ISSN 1043-6618. doi: 10.1016/j.phrs.2017.02.011.

58. Chen Zhang, Haoyue Hu, Xiaoyan Wang, Yajuan Zhu, and Ming Jiang. WFDC Protein: A Promising Diagnosis Biomarker of Ovarian Cancer. Journal of Cancer, 12(18):5404–5412, 2021. ISSN 1837-9664. doi: 10.7150/jca.57880.

59. Lance L. Munn and Igor Garkavtsev. SLPI: a new target for stopping metastasis. Aging (Albany NY), 10(1):13–14, 2018. doi: 10.18632/aging.101372.

60. Masanori Oshi, Yoshihisa Tokumaru, Swagoto Mukhopadhyay, Li Yan, Ryusei Matsuyama, Itaru Endo, and Kazuaki Takabe. Annexin A1 Expression Is Associated with Epithelial–Mesenchymal Transition (EMT), Cell Proliferation, Prognosis, and Drug Response in Pancreatic Cancer. Cells, 10(3):653, 2021. doi: 10.3390/cells10030653.

61. Masaki Shiota, Anousheh Zardan, Ario Takeuchi, Masafumi Kumano, Eliana Beraldi, Seiji Naito, Amina Zoubeidi, and Martin E. Gleave. Clusterin Mediates TGF-β–Induced Epithelial–Mesenchymal Transition and Metastasis via Twist1 in Prostate Cancer Cells. Cancer Research, 72(20):5261–5272, 2012. ISSN 0008-5472. doi: 10.1158/0008-5472.can-12-0254.

62. Richard D. Bennett, Mark R. Pittelkow, and Emanuel E. Strehler. Immunolocalization of the Tumor-Sensitive Calmodulin-Like Protein CALML3 in Normal Human Skin and Hyperproliferative Skin Disorders. PLoS ONE, 8(4):e62347, 2013. doi: 10.1371/journal.pone.0062347.

63. Michael Schubert, Bertram Klinger, Martina Klünemann, Anja Sieber, Florian Uhlitz, Sascha Sauer, Mathew J. Garnett, Nils Blüthgen, and Julio Saez-Rodriguez. Perturbation-response genes reveal signaling footprints in cancer gene expression. Nature Communications, 9(1):20, 2018. doi: 10.1038/s41467-017-02391-6.

64. Ricky W. Johnstone, Ailsa J. Frew, and Mark J. Smyth. The TRAIL apoptotic pathway in cancer onset, progression and therapy. Nature Reviews Cancer, 8(10):782–798, 2008. ISSN 1474-175X. doi: 10.1038/nrc2465.

65. SJ Thomas, JA Snowden, PP Zeidler, and SJ Danson. The role of JAK/STAT signalling in the pathogenesis, prognosis and treatment of solid tumours. British Journal of Cancer, 113(3):365–371, 2015. ISSN 0007-0920. doi: 10.1038/bjc.2015.233.

66. Mustafa Sibai, Sergi Cervilla, Daniela Grases, Eva Musulen, Rossana Lazcano, ChiaKuei Mo, Veronica Davalos, Arola Fortian, Adrià Bernat Margarita Romeo, Collin Tokheim, Enrique Grande, Francisco Real, Jordi Barretina, Alexander J Lazar, Li Ding, Manel Esteller, Matthew H Bailey, and Eduard Porta-Pardo. The spatial landscape of Cancer Hallmarks reveals patterns of tumor ecology. bioRxiv, page 2022.06.18.496114, 2023. doi: 10.1101/2022.06.18.496114.

67. Kari J. Kurppa, Yao Liu, Ciric To, Tinghu Zhang, Mengyang Fan, Amir Vajdi, Erik H. Knelson, Yingtian Xie, Klothilda Lim, Paloma Cejas, Andrew Portell, Patrick H. Lizotte, Scott B. Ficarro, Shuai Li, Ting Chen, Heidi M. Haikala, Haiyun Wang, Magda Bahcall, Yang Gao, Sophia Shalhout, Steffen Boettcher, Bo Hee Shin, Tran Thai, Margaret K. Wilkens, Michelle L. Tillgren, Mierzhati Mushajiang, Man Xu, Jihyun Choi, Arrien A. Bertram, Benjamin L. Ebert, Rameen Beroukhim, Pratiti Bandopadhayay, Mark M. Awad, Prafulla C. Gokhale, Paul T. Kirschmeier, Jarrod A. Marto, Fernando D. Camargo, Rizwan Haq, Cloud P. Paweletz, Kwok-Kin Wong, David A. Barbie, Henry W. Long, Nathanael S. Gray, and Pasi A. Jänne. Treatment-Induced Tumor Dormancy through YAP-Mediated Transcriptional Reprogramming of the Apoptotic Pathway. Cancer Cell, 37(1):104–122.e12, 2020. ISSN 1535-6108. doi: 10.1016/j.ccell.2019.12.006.

68. Cihangir Duy, Meng Li, Matt Teater, Cem Meydan, Francine E. Garrett-Bakelman, Tak C. Lee, Christopher R. Chin, Ceyda Durmaz, Kimihito C. Kawabata, Eugen Dhimolea, Constantine S. Mitsiades, Hartmut Doehner, Richard J. D’Andrea, Michael W. Becker, Elisabeth M. Paietta, Christopher E. Mason, Martin Carroll, and Ari M. Melnick. Chemotherapy Induces Senescence-Like Resilient Cells Capable of Initiating AML Recurrence. Cancer Discovery, 2021. doi: 10.1158/2159-8290.cd-20-1375.

69. Dominik Saul, Robyn Laura Kosinsky, Elizabeth J. Atkinson, Madison L. Doolittle, Xu Zhang, Nathan K. LeBrasseur, Robert J. Pignolo, Paul D. Robbins, Laura J. Niedernhofer, Yuji Ikeno, Diana Jurk, João F. Passos, LaTonya J. Hickson, Ailing Xue, David G. Monroe, Tamara Tchkonia, James L. Kirkland, Joshua N. Farr, and Sundeep Khosla. A new gene set identifies senescent cells and predicts senescence-associated pathways across tissues. Nature Communications, 13(1):4827, 2022. doi: 10.1038/s41467-022-32552-1.

70. Azamat Akhmetkaliyev, Noura Alibrahim, Darya Shafiee, and Eugene Tulchinsky. EMT/MET plasticity in cancer and Go-or-Grow decisions in quiescence: the two sides of the same coin? Molecular Cancer, 22(1):90, 2023. doi: 10.1186/s12943-023-01793-z.

71. Mariangela Russo, Giovanni Crisafulli, Alberto Sogari, Nicole M. Reilly, Sabrina Arena, Simona Lamba, Alice Bartolini, Vito Amodio, Alessandro Magrì, Luca Novara, Ivana Sarotto, Zachary D. Nagel, Cortt G. Piett, Alessio Amatu, Andrea Sartore-Bianchi, Salvatore Siena, Andrea Bertotti, Livio Trusolino, Mattia Corigliano, Marco Gherardi, Marco Cosentino Lagomarsino, Federica Di Nicolantonio, and Alberto Bardelli. Adaptive mutability of colorectal cancers in response to targeted therapies. Science, 366(6472):1473–1480, 2019. ISSN 0036-8075. doi: 10.1126/science.aav4474.

72. Arthur Liberzon, Chet Birger, Helga Thorvaldsdóttir, Mahmoud Ghandi, Jill P. Mesirov, and Pablo Tamayo. The Molecular Signatures Database Hallmark Gene Set Collection. Cell Systems, 1(6):417–425, 2015. ISSN 2405-4712. doi: 10.1016/j.cels.2015.12.004.

73. Michael Ashburner, Catherine A. Ball, Judith A. Blake, David Botstein, Heather Butler, J. Michael Cherry, Allan P. Davis, Kara Dolinski, Selina S. Dwight, Janan T. Eppig, Midori A. Harris, David P. Hill, Laurie Issel-Tarver, Andrew Kasarskis, Suzanna Lewis, John C. Matese, Joel E. Richardson, Martin Ringwald, Gerald M. Rubin, and Gavin Sherlock. Gene Ontology: tool for the unification of biology. Nature Genetics, 25(1):25–29, 2000. ISSN 1061-4036. doi: 10.1038/75556.

74. Candace C. Liu, Noah F. Greenwald, Alex Kong, Erin F. McCaffrey, Ke Xuan Leow, Dunja Mrdjen, Bryan J. Cannon, Josef Lorenz Rumberger, Sricharan Reddy Varra, and Michael Angelo. Robust phenotyping of highly multiplexed tissue imaging data using pixel-level clustering. Nature Communications, 14(1):4618, 2023. doi: 10.1038/s41467-023-40068-5.

75. F.G. Kondev, M. Wang, W.J. Huang, S. Naimi, and G. Audi. The NUBASE2020 evaluation of nuclear physics properties. Chinese Physics C, 45(3):030001, 2021. ISSN 1674-1137. doi: 10.1088/1674-1137/abddae.

76. Zahid H Siddik. Cisplatin: mode of cytotoxic action and molecular basis of resistance. Oncogene, 22(47):7265–7279, 2003. ISSN 0950-9232. doi: 10.1038/sj.onc.1206933.

77. Qing Chang, Olga I. Ornatsky, Iram Siddiqui, Rita Straus, Vladimir I. Baranov, and David W. Hedley. Biodistribution of cisplatin revealed by imaging mass cytometry identifies extensive collagen binding in tumor and normal tissues. Scientific Reports, 6(1):36641, 2016. doi:10.1038/srep36641.

78. Andreas R. de Biasi, Jonathan Villena-Vargas, and Prasad S. Adusumilli. Cisplatin-Induced Antitumor Immunomodulation: A Review of Preclinical and Clinical Evidence. Clinical Cancer Research, 20(21):5384–5391, 2014. ISSN 1078-0432. doi: 10.1158/1078-0432.ccr-14-1298.

79. Jovan Tanevski, Ricardo Omar Ramirez Flores, Attila Gabor, Denis Schapiro, and Julio Saez-Rodriguez. Explainable multiview framework for dissecting spatial relationships from highly multiplexed data. Genome Biology, 23(1):97, 2022. ISSN 1474-7596. doi: 10.1186/s13059-022-02663-5.

80. Abbes Belkhiri, Altaf A. Dar, Dun Fa Peng, Mohammad H. Razvi, Cammie Rinehart, Carlos L. Arteaga, and Wael El-Rifai. Expression of t-DARPP Mediates Trastuzumab Resistance in Breast Cancer Cells. Clinical Cancer Research, 14(14):4564–4571, 2008. ISSN 1078-0432. doi: 10.1158/1078-0432.ccr-08-0121.

81. Nawal Bendris, Karla C. Williams, Carlos R. Reis, Erik S. Welf, Ping-Hung Chen, Bénédicte Lemmers, Michael Hahne, Hon Sing Leong, and Sandra L. Schmid. SNX9 promotes metastasis by enhancing cancer cell invasion via differential regulation of RhoGTPases. Molecular Biology of the Cell, 27(9):1409–1419, 2016. ISSN 1059-1524. doi: 10.1091/mbc.e16-02-0101.

82. Lao H Saal, Johan Vallon-Christersson, Jari Häkkinen, Cecilia Hegardt, Dorthe Grabau, Christof Winter, Christian Brueffer, Man-Hung Eric Tang, Christel Reuterswärd, Ralph Schulz, Anna Karlsson, Anna Ehinger, Janne Malina, Jonas Manjer, Martin Malmberg, Christer Larsson, Lisa Rydén, Niklas Loman, and Åke Borg. The Sweden Cancerome Analysis Network - Breast (SCAN-B) Initiative: a large-scale multicenter infrastructure towards implementation of breast cancer genomic analyses in the clinical routine. Genome Medicine, 7(1):20, 2015. ISSN 1756-994X. doi: 10.1186/s13073-015-0131-9.

83. Kyle Chang, Chad J Creighton, Caleb Davis, Lawrence Donehower, Jennifer Drummond, David Wheeler, Adrian Ally, Miruna Balasundaram, Inanc Birol, Yaron S N Butterfield, Andy Chu, Eric Chuah, Hye-Jung E Chun, Noreen Dhalla, Ranabir Guin, Martin Hirst, Carrie Hirst, Robert A Holt, Steven J M Jones, Darlene Lee, Haiyan I Li, Marco A Marra, Michael Mayo, Richard A Moore, Andrew J Mungall, A Gordon Robertson, Jacqueline E Schein, Payal Sipahimalani, Angela Tam, Nina Thiessen, Richard J Varhol, Rameen Beroukhim, Ami S Bhatt, Angela N Brooks, Andrew D Cherniack, Samuel S Freeman, Stacey B Gabriel, Elena Helman, Joonil Jung, Matthew Meyerson, Akinyemi I Ojesina, Chandra Sekhar Pedamallu, Gordon Saksena, Steven E Schumacher, Barbara Tabak, Travis Zack, Eric S Lander, Christopher A Bristow, Angela Hadjipanayis, Psalm Haseley, Raju Kucherlapati, Semin Lee, Eunjung Lee, Lovelace J Luquette, Harshad S Mahadeshwar, Angeliki Pantazi, Michael Parfenov, Peter J Park, Alexei Protopopov, Xiaojia Ren, Netty Santoso, Jonathan Seidman, Sahil Seth, Xingzhi Song, Jiabin Tang, Ruibin Xi, Andrew W Xu, Lixing Yang, Dong Zeng, J Todd Auman, Saianand Balu, Elizabeth Buda, Cheng Fan, Katherine A Hoadley, Corbin D Jones, Shaowu Meng, Piotr A Mieczkowski, Joel S Parker, Charles M Perou, Jeffrey Roach, Yan Shi, Grace O Silva, Donghui Tan, Umadevi Veluvolu, Scot Waring, Matthew D Wilkerson, Junyuan Wu, Wei Zhao, Tom Bodenheimer, D Neil Hayes, Alan P Hoyle, Stuart R Jeffreys, Lisle E Mose, Janae V Simons, Mathew G Soloway, Stephen B Baylin, Benjamin P Berman, Moiz S Bootwalla, Ludmila Danilova, James G Herman, Toshinori Hinoue, Peter W Laird, Suhn K Rhie, Hui Shen, Timothy Triche, Daniel J Weisenberger, Scott L Carter, Kristian Cibulskis, Lynda Chin, Jianhua Zhang, Gad Getz, Carrie Sougnez, Min Wang, Gordon Saksena, Scott L Carter, Kristian Cibulskis, Lynda Chin, Jianhua Zhang, Gad Getz, Huyen Dinh, Harsha Vardhan Doddapaneni, Richard Gibbs, Preethi Gunaratne, Yi Han, Divya Kalra, Christie Kovar, Lora Lewis, Margaret Morgan, Donna Morton, Donna Muzny, Jeffrey Reid, Liu Xi, Juok Cho, Daniel DiCara, Scott Frazer, Nils Gehlenborg, David I Heiman, Jaegil Kim, Michael S Lawrence, Pei Lin, Yingchun Liu, Michael S Noble, Petar Stojanov, Doug Voet, Hailei Zhang, Lihua Zou, Chip Stewart, Brady Bernard, Ryan Bressler, Andrea Eakin, Lisa Iype, Theo Knijnenburg, Roger Kramer, Richard Kreisberg, Kalle Leinonen, Jake Lin, Yuexin Liu, Michael Miller, Sheila M Reynolds, Hector Rovira, Ilya Shmulevich, Vesteinn Thorsson, D. Yang, Wei Zhang, Samirkumar Amin, Chang-Jiun Wu, Chia-Chin Wu, Rehan Akbani, Kenneth Aldape, Keith A Baggerly, Bradley Broom, Tod D Casasent, James Cleland, Chad Creighton, Deepti Dodda, Mary Edgerton, Leng Han, Shelley M Herbrich, Zhenlin Ju, Hoon Kim, Seth Lerner, Jun Li, Han Liang, Wenbin Liu, Philip L Lorenzi, Yiling Lu, James Melott, Gordon B Mills, Lam Nguyen, Xiaoping Su, Roeland Verhaak, Wenyi Wang, John N Weinstein, Andrew Wong, Yang Yang, Jun Yao, Rong Yao, Kosuke Yoshihara, Yuan Yuan, Alfred K Yung, Nianxiang Zhang, Siyuan Zheng, Michael Ryan, David W Kane, B Arman Aksoy, Giovanni Ciriello, Gideon Dresdner, Jianjiong Gao, Benjamin Gross, Anders Jacobsen, Andre Kahles, Marc Ladanyi, William Lee, Kjong-Van Lehmann, Martin L Miller, Ricardo Ramirez, Gunnar Rätsch, Boris Reva, Chris Sander, Nikolaus Schultz, Yasin Senbabaoglu, Ronglai Shen, Rileen Sinha, S Onur Sumer, Yichao Sun, Barry S Taylor, Nils Weinhold, Suzanne Fei, Paul Spellman, Christopher Benz, Daniel Carlin, Melisssa Cline, Brian Craft, Kyle Ellrott, Mary Goldman, David Haussler, Singer Ma, Sam Ng, Evan Paull, Amie Radenbaugh, Sofie Salama, Artem Sokolov, Joshua M Stuart, Teresa Swatloski, Vladislav Uzunangelov, Peter Waltman, Christina Yau, Jing Zhu, Stanley R Hamilton, Gad Getz, Carrie Sougnez, Scott Abbott, Rachel Abbott, Nathan D Dees, Kim Delehaunty, Li Ding, David J Dooling, Jim M Eldred, Catrina C Fronick, Robert Fulton, Lucinda L Fulton, Joelle Kalicki-Veizer, Krishna-Latha Kanchi, Cyriac Kandoth, Daniel C Koboldt, David E Larson, Timothy J Ley, Ling Lin, Charles Lu, Vincent J Magrini, Elaine R Mardis, Michael D McLellan, Joshua F McMichael, Christopher A Miller, Michelle O’Laughlin, Craig Pohl, Heather Schmidt, Scott M Smith, Jason Walker, John W Wallis, Michael C Wendl, Richard K Wilson, Todd Wylie, Qunyuan Zhang, Robert Burton, Mark A Jensen, Ari Kahn, Todd Pihl, David Pot, Yunhu Wan, Douglas A Levine, Aaron D Black, Jay Bowen, Jessica Frick, Julie M Gastier-Foster, Hollie A Harper, Carmen Helsel, Kristen M Leraas, Tara M Lichtenberg, Cynthia McAllister, Nilsa C Ramirez, Samantha Sharpe, Lisa Wise, Erik Zmuda, Stephen J Chanock, Tanja Davidsen, John A Demchok, Greg Eley, Ina Felau, Brad A Ozenberger, Margi Sheth, Heidi Sofia, Louis Staudt, Roy Tarnuzzer, Zhining Wang, Liming Yang, Jiashan Zhang, Larsson Omberg, Adam Margolin, Benjamin J Raphael, Fabio Vandin, Hsin-Ta Wu, Mark D M Leiserson, Stephen C Benz, Charles J Vaske, Houtan Noushmehr, Theo Knijnenburg, Denise Wolf, Laura Van’t Veer, Eric A Collisson, Dimitris Anastassiou, Tai-Hsien Ou Yang, Nuria Lopez-Bigas, Abel Gonzalez-Perez, David Tamborero, Zheng Xia, Wei Li, Dong-Yeon Cho, Teresa Przytycka, Mark Hamilton, Sean McGuire, Sven Nelander, Patrik Johansson, Rebecka Jörnsten, Teresia Kling, Jose Sanchez, John N Weinstein, Eric A Collisson, Gordon B Mills, Kenna R Mills Shaw, Brad A Ozenberger, Kyle Ellrott, Ilya Shmulevich, Chris Sander, and Joshua M Stuart. The Cancer Genome Atlas Pan-Cancer analysis project. Nature Genetics, 45(10):1113–1120, 2013. ISSN 1061-4036. doi: 10.1038/ng.2764.

84. Anushka Dongre, Mohammad Rashidian, Ferenc Reinhardt, Aaron Bagnato, Zuzana Keckesova, Hidde L. Ploegh, and Robert A. Weinberg. Epithelial-to-Mesenchymal Transition Contributes to Immunosuppression in Breast Carcinomas. Cancer Research, 77(15): 3982–3989, 2017. ISSN 0008-5472. doi: 10.1158/0008-5472.can-16-3292.

85. Anushka Dongre and Robert A. Weinberg. New insights into the mechanisms of epithelial–mesenchymal transition and implications for cancer. Nature Reviews Molecular Cell Biology, 20(2):69–84, 2019. ISSN 1471-0072. doi: 10.1038/s41580-018-0080-4.

86. Javier De Las Rivas, Anamaria Brozovic, Sivan Izraely, Alba Casas-Pais, Isaac P. Witz, and Angélica Figueroa. Cancer drug resistance induced by EMT: novel therapeutic strategies. Archives of Toxicology, 95(7):2279–2297, 2021. ISSN 0340-5761. doi: 10.1007/s00204-021-03063-7.

87. Yuhe Huang, Weiqi Hong, and Xiawei Wei. The molecular mechanisms and therapeutic strategies of EMT in tumor progression and metastasis. Journal of Hematology & Oncology, 15(1):129, 2022. doi: 10.1186/s13045-022-01347-8.

88. Tsukasa Shibue and Robert A. Weinberg. EMT, CSCs, and drug resistance: the mechanistic link and clinical implications. Nature Reviews Clinical Oncology, 14(10):611–629, 2017. ISSN 1759-4774. doi: 10.1038/nrclinonc.2017.44.

89. M Saxena, MA Stephens, H Pathak, and A Rangarajan. Transcription factors that mediate epithelial–mesenchymal transition lead to multidrug resistance by upregulating ABC transporters. Cell Death & Disease, 2(7):e179–e179, 2011. doi: 10.1038/cddis.2011.61.

90. Yu-xiong Feng, Ethan S. Sokol, Catherine A. Del Vecchio, Sandhya Sanduja, Jasper H.L. Claessen, Theresa A. Proia, Dexter X. Jin, Ferenc Reinhardt, Hidde L. Ploegh, Qiu Wang, and Piyush B. Gupta. Epithelial-to-Mesenchymal Transition Activates PERK–eIF2α and Sensitizes Cells to Endoplasmic Reticulum Stress. Cancer Discovery, 4(6):702–715, 2014. ISSN 2159-8274. doi: 10.1158/2159-8290.cd-13-0945.

91. Nasim Ebrahimi, Mahdokht Sadat Manavi, Ferdos Faghihkhorasani, Siavash Seifollahy Fakhr, Fatemeh Jafari Baei, Fereshteh Faghih Khorasani, Mohammad Mehdi Zare, Nazanin Pazhouhesh Far, Fatemeh Rezaei-Tazangi, Jun Ren, Russel J. Reiter, Noushin Nabavi, Amir Reza Aref, Chu Chen, Yavuz Nuri Ertas, and Qi Lu. Harnessing function of EMT in cancer drug resistance: a metastasis regulator determines chemotherapy response. Cancer and Metastasis Reviews, 43(1):457–479, 2024. ISSN 0167-7659. doi: 10.1007/s10555-023-10162-7.

92. Maud Debaugnies, Sara Rodríguez-Acebes, Jeremy Blondeau, Marie-Astrid Parent, Manuel Zocco, Yura Song, Viviane de Maertelaer, Virginie Moers, Mathilde Latil, Christine Dubois, Katia Coulonval, Francis Impens, Delphi Van Haver, Sara Dufour, Akiyoshi Uemura, Panagiota A. Sotiropoulou, Juan Méndez, and Cédric Blanpain. RHOJ controls EMT-associated resistance to chemotherapy. Nature, 616(7955):168–175, 2023. ISSN 0028-0836. doi: 10.1038/s41586-023-05838-7.

93. Vita Fedele and Davide Melisi. Permissive State of EMT: The Role of Immune Cell Compartment. Frontiers in Oncology, 10:587, 2020. ISSN 2234-943X. doi: 10.3389/fonc.2020.00587.

94. Hye-Young Min and Ho-Young Lee. Cellular Dormancy in Cancer - Mechanisms and Potential Targeting Strategies. Cancer Research and Treatment, 55(3):720–736, 2023. ISSN 1598-2998. doi: 10.4143/crt.2023.468.

95. Suzanne S. Bohlson, Sean D. O’Conner, Holly Jo Hulsebus, Minh-Minh Ho, and Deborah A. Fraser. Complement, C1q, and C1q-Related Molecules Regulate Macrophage Polarization. Frontiers in Immunology, 5:402, 2014. ISSN 1664-3224. doi: 10.3389/fimmu.2014.00402.

96. Wan Liu, Wenjing Wang, Xinran Wang, Cong Xu, Ning Zhang, and Wen Di. Cisplatin-stimulated macrophages promote ovarian cancer migration via the CCL20-CCR6 axis. Cancer Letters, 472:59–69, 2020. ISSN 0304-3835. doi: 10.1016/j.canlet.2019.12.024.

97. Peter Bankhead, Maurice B. Loughrey, José A. Fernández, Yvonne Dombrowski, Darragh G. McArt, Philip D. Dunne, Stephen McQuaid, Ronan T. Gray, Liam J. Murray, Helen G. Coleman, Jacqueline A. James, Manuel Salto-Tellez, and Peter W. Hamilton. QuPath: Open source software for digital pathology image analysis. Scientific Reports, 7 (1):16878, 2017. doi: 10.1038/s41598-017-17204-5.

98. Fabienne Birrer, Tess Brodie, and Deborah Stroka. OMIP-088: Twenty-target imaging mass cytometry panel for major cell populations in mouse formalin fixed paraffin embedded liver. Cytometry Part A, 103(3):189–192, 2023. ISSN 1552-4922. doi: 10.1002/cyto.a.24714.

99. F. Alexander Wolf, Philipp Angerer, and Fabian J. Theis. SCANPY: large-scale single-cell gene expression data analysis. Genome Biology, 19(1):15, 2018. ISSN 1474-7596. doi:10.1186/s13059-017-1382-0.

100. Santiago J. Carmona, Imran Siddiqui, Mariia Bilous, Werner Held, and David Gfeller. Deciphering the transcriptomic landscape of tumor-infiltrating CD8 lymphocytes in B16 melanoma tumors with single-cell RNA-Seq. OncoImmunology, 9(1):1737369, 2020. ISSN 2162-4011. doi: 10.1080/2162402x.2020.1737369.

101. Pau Badia-i Mompel, Jesús Vélez Santiago, Jana Braunger, Celina Geiss, Daniel Dimitrov, Sophia Müller-Dott, Petr Taus, Aurelien Dugourd, Christian H Holland, Ricardo O Ramirez Flores, and Julio Saez-Rodriguez. decoupleR: ensemble of computational methods to infer biological activities from omics data. Bioinformatics Advances, 2(1):vbac016, 2022. doi:10.1093/bioadv/vbac016.

102. Boris Muzellec, Maria Telenčzuk, Vincent Cabeli, and Mathieu Andreux. PyDESeq2: a python package for bulk RNA-seq differential expression analysis. bioRxiv, page 2022.12.14.520412, 2022. doi: 10.1101/2022.12.14.520412.

103. Anqi Zhu, Joseph G Ibrahim, and Michael I Love. Heavy-tailed prior distributions for sequence count data: removing the noise and preserving large differences. Bioinformatics, 35(12):2084–2092, 2018. ISSN 1367-4803. doi: 10.1093/bioinformatics/bty895.

104. Gioele La Manno, Ruslan Soldatov, Amit Zeisel, Emelie Braun, Hannah Hochgerner, Viktor Petukhov, Katja Lidschreiber, Maria E. Kastriti, Peter Lönnerberg, Alessandro Furlan, Jean Fan, Lars E. Borm, Zehua Liu, David van Bruggen, Jimin Guo, Xiaoling He, Roger Barker, Erik Sundström, Gonçalo Castelo-Branco, Patrick Cramer, Igor Adameyko, Sten Linnarsson, and Peter V. Kharchenko. RNA velocity of single cells. Nature, 560(7719): 494–498, 2018. ISSN 0028-0836. doi: 10.1038/s41586-018-0414-6.

105. Volker Bergen, Marius Lange, Stefan Peidli, F. Alexander Wolf, and Fabian J. Theis. Generalizing RNA velocity to transient cell states through dynamical modeling. Nature Biotechnology, 38(12):1408–1414, 2020. ISSN 1087-0156. doi: 10.1038/s41587-020-0591-3.

106. Ilya Korsunsky, Nghia Millard, Jean Fan, Kamil Slowikowski, Fan Zhang, Kevin Wei, Yuriy Baglaenko, Michael Brenner, Po-ru Loh, and Soumya Raychaudhuri. Fast, sensitive and accurate integration of single-cell data with Harmony. Nature Methods, 16(12):1289–1296, 2019. ISSN 1548-7091. doi: 10.1038/s41592-019-0619-0.

107. Brian Hie, Bryan Bryson, and Bonnie Berger. Efficient integration of heterogeneous singlecell transcriptomes using Scanorama. Nature Biotechnology, 37(6):685–691, 2019. ISSN 1087-0156. doi: 10.1038/s41587-019-0113-3.

108. Fabian Pedregosa, Gaël Varoquaux, Alexandre Gramfort, Vincent Michel, Bertrand Thirion, Olivier Grisel, Mathieu Blondel, Andreas Müller, Joel Nothman, Gilles Louppe, Peter Prettenhofer, Ron Weiss, Vincent Dubourg, Jake Vanderplas, Alexandre Passos, David Cournapeau, Matthieu Brucher, Matthieu Perrot, and Édouard Duchesnay. Scikitlearn: Machine Learning in Python. arXiv, 45(6):2041–2049, 2012. ISSN 0031-3203. doi:10.48550/arxiv.1201.0490.

109. Rahul Satija, Jeffrey A Farrell, David Gennert, Alexander F Schier, and Aviv Regev. Spatial reconstruction of single-cell gene expression data. Nature Biotechnology, 33(5):495–502, 2015. ISSN 1087-0156. doi: 10.1038/nbt.3192.

110. Sophia Müller-Dott, Eirini Tsirvouli, Miguel Vazquez, Ricardo O Ramirez Flores, Pau Badia-i Mompel, Robin Fallegger, Dénes Türei, Astrid Lægreid, and Julio Saez-Rodriguez. Expanding the coverage of regulons from high-confidence prior knowledge for accurate estimation of transcription factor activities. Nucleic Acids Research, 51(20):10934–10949, 2023. ISSN 0305-1048. doi: 10.1093/nar/gkad841.

111. Anoop P. Patel, Itay Tirosh, John J. Trombetta, Alex K. Shalek, Shawn M. Gillespie, Hiroaki Wakimoto, Daniel P. Cahill, Brian V. Nahed, William T. Curry, Robert L. Martuza, David N. Louis, Orit Rozenblatt-Rosen, Mario L. Suvà, Aviv Regev, and Bradley E. Bernstein. Single-cell RNA-seq highlights intratumoral heterogeneity in primary glioblastoma. Science, 344(6190):1396–1401, 2014. ISSN 0036-8075. doi: 10.1126/science.1254257.

112. Pauli Virtanen, Ralf Gommers, Travis E Oliphant, Matt Haberland, Tyler Reddy, David Cournapeau, Evgeni Burovski, Pearu Peterson, Warren Weckesser, Jonathan Bright, Stéfan J van der Walt, Matthew Brett, Joshua Wilson, K Jarrod Millman, Nikolay Mayorov, Andrew R J Nelson, Eric Jones, Robert Kern, Eric Larson, CJ Carey, İlhan Polat, Yu Feng, Eric W Moore, Jake VanderPlas, Denis Laxalde, Josef Perktold, Robert Cimrman, Ian Henriksen,A A Quintero, Charles R Harris, Anne M Archibald, Antônio H Ribeiro, Fabian Pedregosa, Paul van Mulbregt, SciPy 1 0 Contributors, Aditya Vijaykumar, Alessandro Pietro Bardelli, Alex Rothberg, Andreas Hilboll, Andreas Kloeckner, Anthony Scopatz, Antony Lee, Ariel Rokem, C Nathan Woods, Chad Fulton, Charles Masson, Christian Häggström, Clark Fitzgerald, David A Nicholson, David R Hagen, Dmitrii V Pasechnik, Emanuele Olivetti, Eric Martin, Eric Wieser, Fabrice Silva, Felix Lenders, Florian Wilhelm, G Young, Gavin A Price, Gert-Ludwig Ingold, Gregory E Allen, Gregory R Lee, Hervé Audren Irvin Probst, Jörg P Dietrich, Jacob Silterra, James T Webber, Janko Slavič, Joel Nothman, Johannes Buchner, Johannes Kulick, Johannes L Schönberger, José Vinícius de Miranda Cardoso, Joscha Reimer, Joseph Harrington, Juan Luis Cano Rodríguez, Juan Nunez-Iglesias, Justin Kuczynski, Kevin Tritz, Martin Thoma, Matthew Newville, Matthias Kümmerer, Maximilian Bolingbroke, Michael Tartre, Mikhail Pak, Nathaniel J Smith, Nikolai Nowaczyk, Nikolay Shebanov, Oleksandr Pavlyk, Per A Brodtkorb, Perry Lee, Robert T McGibbon, Roman Feldbauer, Sam Lewis, Sam Tygier, Scott Sievert, Sebastiano Vigna, Stefan Peterson, Surhud More, Tadeusz Pudlik, Takuya Oshima, Thomas J Pingel, Thomas P Robitaille, Thomas Spura, Thouis R Jones, Tim Cera, Tim Leslie, Tiziano Zito, Tom Krauss, Utkarsh Upadhyay, Yaroslav O Halchenko, and Yoshiki Vázquez-Baeza. SciPy 1.0: fundamental algorithms for scientific computing in Python. Nature Methods, 17 (3):261–272, 2020. ISSN 1548-7091. doi: 10.1038/s41592-019-0686-2.

113. Luca Marconato, Giovanni Palla, Kevin A. Yamauchi, Isaac Virshup, Elyas Heidari, Tim Treis, Wouter-Michiel Vierdag, Marcella Toth, Sonja Stockhaus, Rahul B. Shrestha, Benjamin Rombaut, Lotte Pollaris, Laurens Lehner, Harald Vöhringer, Ilia Kats, Yvan Saeys, Sinem K. Saka, Wolfgang Huber, Moritz Gerstung, Josh Moore, Fabian J. Theis, and Oliver Stegle. SpatialData: an open and universal data framework for spatial omics. Nature Methods, 22(1):58–62, 2025. ISSN 1548-7091. doi: 10.1038/s41592-024-02212-x.

114. Noah F. Greenwald, Geneva Miller, Erick Moen, Alex Kong, Adam Kagel, Thomas Dougherty, Christine Camacho Fullaway, Brianna J. McIntosh, Ke Xuan Leow, Morgan Sarah Schwartz, Cole Pavelchek, Sunny Cui, Isabella Camplisson, Omer Bar-Tal, Jaiveer Singh, Mara Fong, Gautam Chaudhry, Zion Abraham, Jackson Moseley, Shiri Warshawsky, Erin Soon, Shirley Greenbaum, Tyler Risom, Travis Hollmann, Sean C. Bendall, Leeat Keren, William Graf, Michael Angelo, and David Van Valen. Whole-cell segmentation of tissue images with human-level performance using large-scale data annotation and deep learning. Nature Biotechnology, 40(4):555–565, 2022. ISSN 1087-0156. doi: 10.1038/s41587-021-01094-0.

115. Stefan van der Walt, Johannes L Schönberger, Juan Nunez-Iglesias, François Boulogne, Joshua D Warner, Neil Yager, Emmanuelle Gouillart, Tony Yu, and the scikit-image contributors. scikit-image: Image processing in Python. arXiv, 2014. doi: 10.48550/arxiv.1407.6245.

116. Giovanni Palla, Hannah Spitzer, Michal Klein, David Fischer, Anna Christina Schaar, Louis Benedikt Kuemmerle, Sergei Rybakov, Ignacio L. Ibarra, Olle Holmberg, Isaac Virshup, Mohammad Lotfollahi, Sabrina Richter, and Fabian J. Theis. Squidpy: a scalable framework for spatial omics analysis. Nature Methods, 19(2):171–178, 2022. ISSN 1548-7091. doi: 10.1038/s41592-021-01358-2.

117. Daniel Dimitrov, Philipp Sven Lars Schäfer, Elias Farr, Pablo Rodriguez-Mier, Sebastian Lobentanzer, Pau Badia-i Mompel, Aurelien Dugourd, Jovan Tanevski, Ricardo Omar Ramirez Flores, and Julio Saez-Rodriguez. LIANA+ provides an all-in-one framework for cell–cell communication inference. Nature Cell Biology, 26(9):1613–1622, 2024. ISSN 1465-7392. doi: 10.1038/s41556-024-01469-w.

118. Cameron Davidson-Pilon. lifelines: survival analysis in Python. Journal of Open Source Software, 4(40):1317, 2019. doi: 10.21105/joss.01317.

119. John D. Hunter. Matplotlib: A 2D Graphics Environment. Computing in Science & Engineering, 9(3):90–95, 2007. ISSN 1521-9615. doi: 10.1109/mcse.2007.55.

120. Michael Waskom. seaborn: statistical data visualization. Journal of Open Source Software, 6(60):3021, 2021. doi: 10.21105/joss.03021.

