## Supplementary Information for "Spatiotemporal organisation of residual disease in mouse and human BRCA1-deficient mammary tumours and breast cancer"

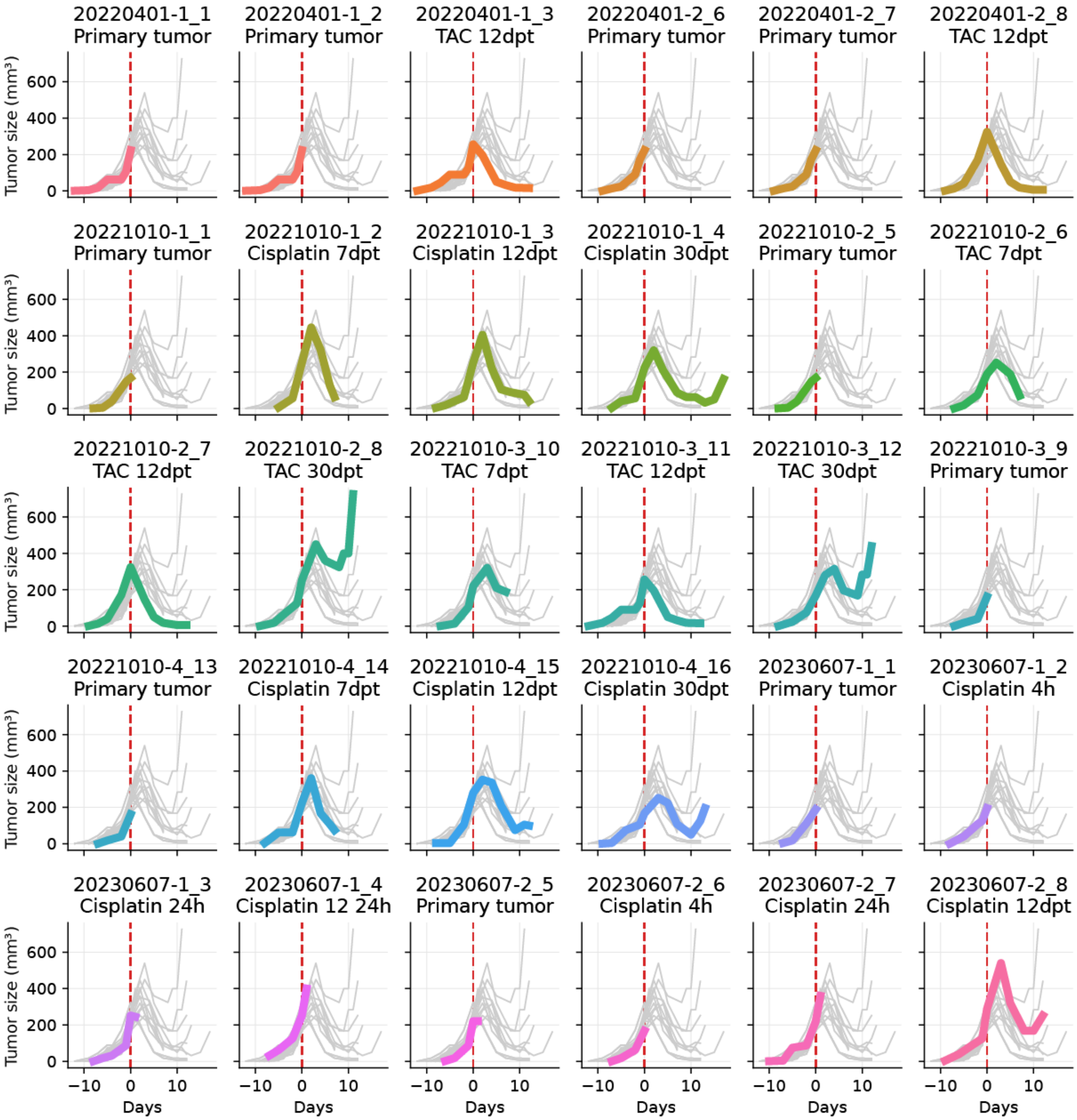


**Supplementary Fig. 1.** | **Tumour growth in the KB1P mouse model.** Tumour growth curves of KB1P mammary tumours. Red dashed lines indicate chemotherapy (cisplatin, TAC) treatments.

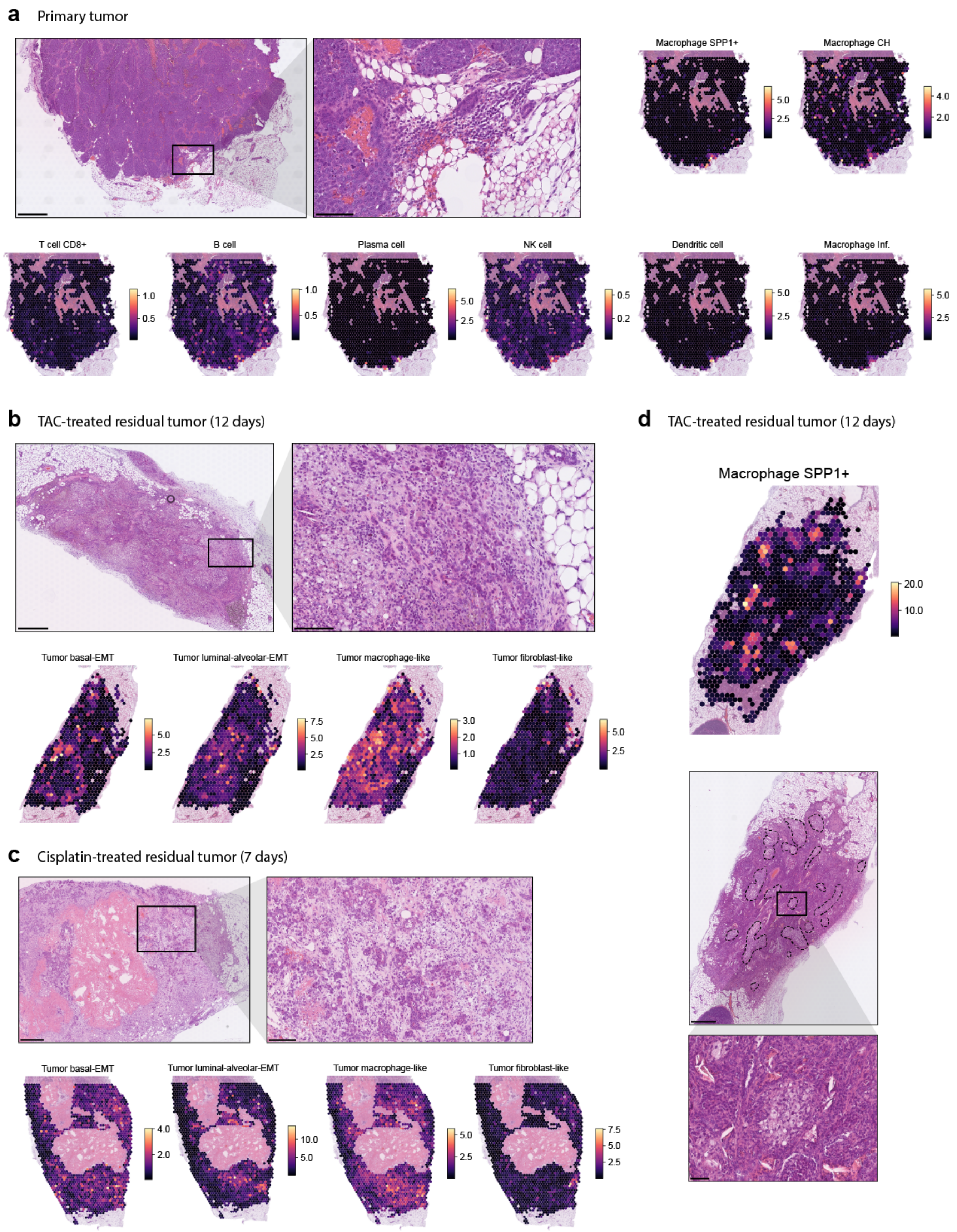


**Supplementary Fig. 2.** | **Spatiotemporal histopathological changes in MRD. a,** Representative H&E images of a primary tumour sample showing mixed immune cell infiltration along the tumour margins, with the corresponding spatial localisation of the indicated cell types within the Visium tissue section (scale bars: left-500 µm, right-100 µm). (continued on next page)

**Supplementary Fig. 2.** | (continued) **b,** Representative H&E images of a TAC-treated residual tumour with the corresponding spatial localisation of the indicated tumour cell types within the Visium tissue section. The epithelial tumour cells gradually lost cell-cell contact, became more elongated, and were increasingly difficult to identify within the reactive stroma (scale bar: left-500 µm, right-100 µm). **c,** Representative H&E images of a cisplatin-treated residual tumour with the corresponding spatial localisation of the indicated tumour cell types within the Visium tissue section. Similar morphological changes were observed in the tumour tissue, albeit to a lesser extent. Epithelial clusters remained more defined (scale bar: left-500 µm, right-100 µm). **d,** Spatial localisation of *Spp1+* macrophages within the Visium tissue section, alongside the corresponding H&E image in a TAC-treated residual tumour. Foamy *Spp1+* macrophages are delineated in blue (scale bar: 500 µm) and depicted at higher magnification (lower panel). The spatial plots in panels a, b, and d depict cell type abundance data inferred with cell2location.

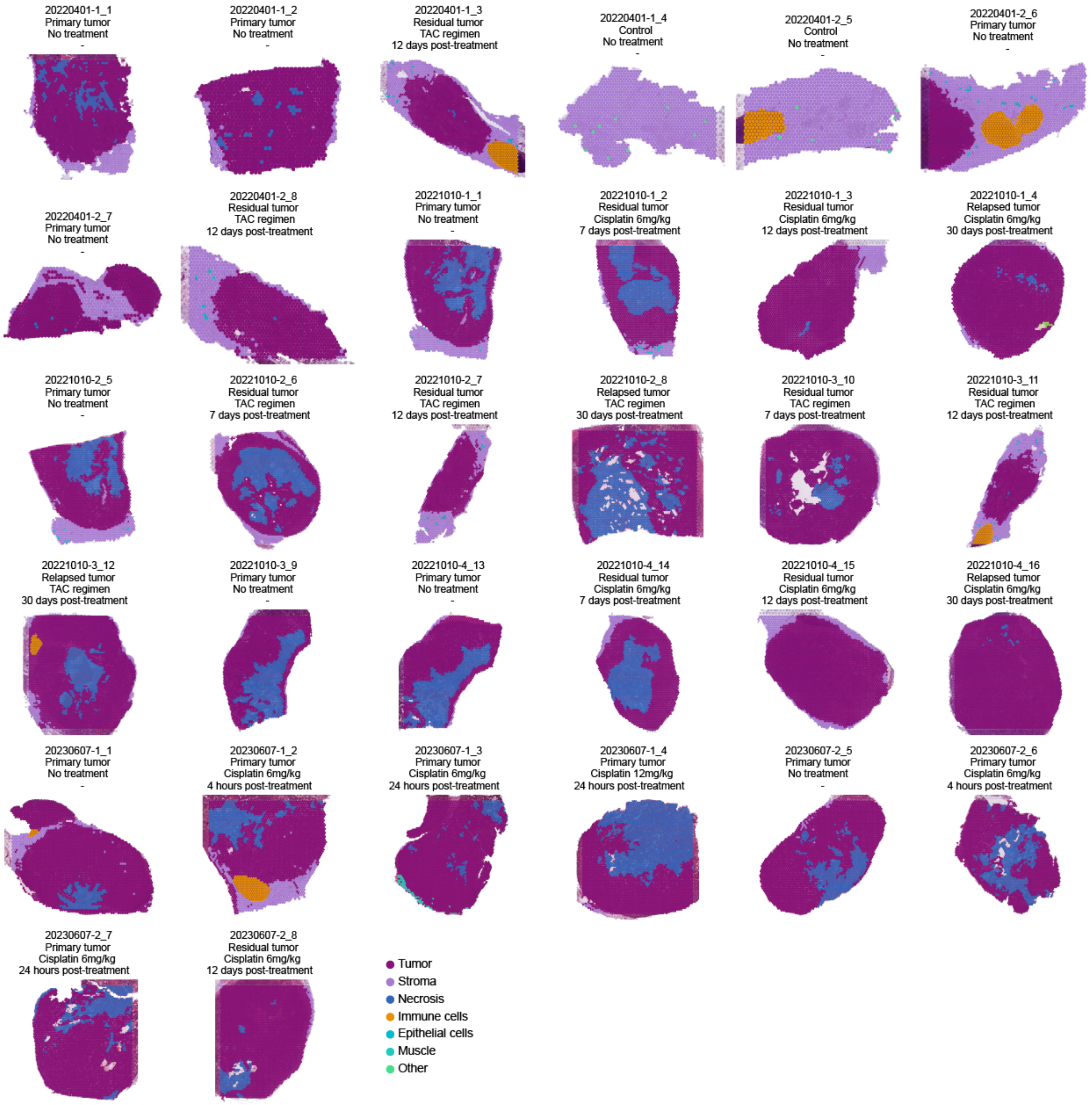


**Supplementary Fig. 3.** | **Histopathological annotations of capture spots.** Labelling of capture spots based on histopathological annotations. For downstream analysis, only capture spots containing tumour tissue were considered. Categories include: Tumour (tumour tissue), Stroma (mammary fat pad), Necrosis (dead tissue), Immune cells (lymph node), Epithelial cells (mammary gland ducts), Muscle (muscle tissue), Other/ignore (undefined tissue).

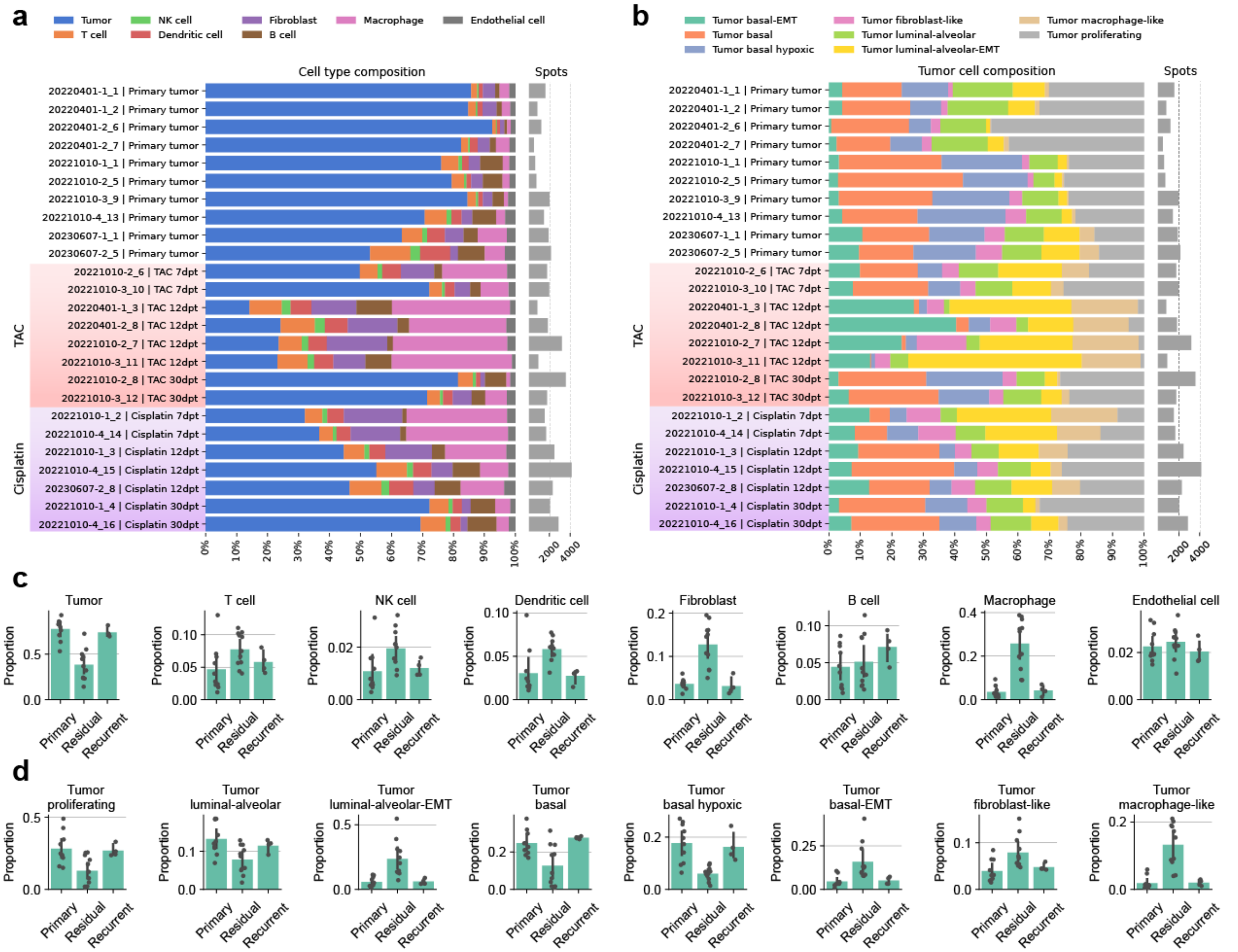


**Supplementary Fig. 4.** | **Reversible changes in tumour and immune cell composition following treatment.** **a,** Cell type composition of ST tissue samples inferred with cell type deconvolution. The proportion of tumour cells significantly decreased in residual tumours, while the proportions of immune cells and fibroblasts increased, regardless of the treatment regimen. These proportions reverted in the recurrent tumours. **b,** Tumour cell composition of ST tissue samples after cell type deconvolution. The proportions of proliferating, basal, luminal-alveolar, and basal hypoxic tumour cells significantly decreased in residual tumours, while the proportions of EMT, macrophage-like, and fibroblast-like tumour cells increased, regardless of the treatment regimen. These proportions reverted in the recurrent tumours. **c,** Barplots showing the mean cell type fraction across all samples (black dots: individual ST samples, error bar: 95% CI, bar height: mean value).

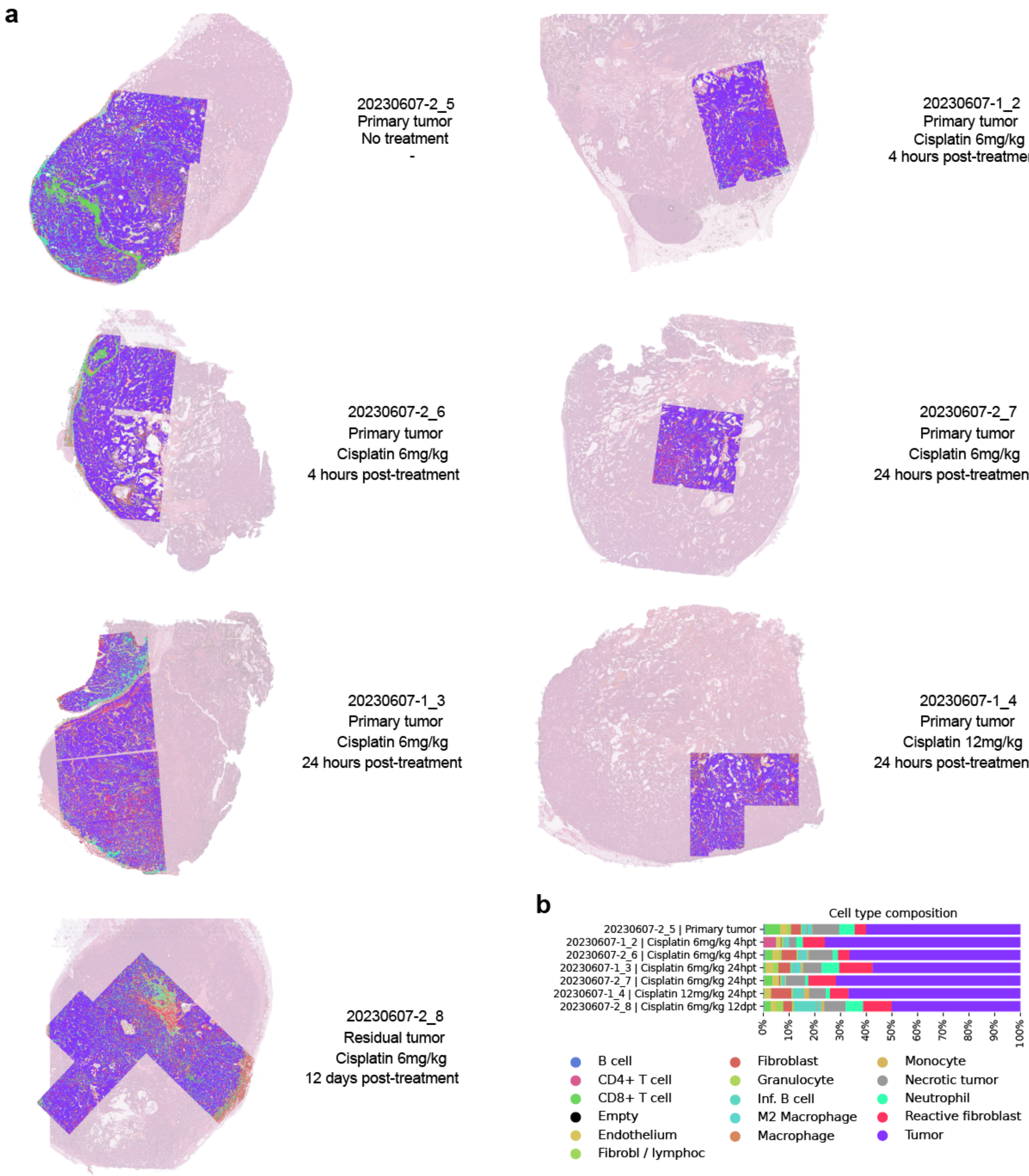


**Supplementary Fig. 5.** | ***In situ* single-cell composition of mammary tumours via IMC. a,** Reconstructed *in situ* single-cell composition of tumour sections derived from IMC measurements. **b,** Variations in cell type composition across different tumours following cisplatin treatment.

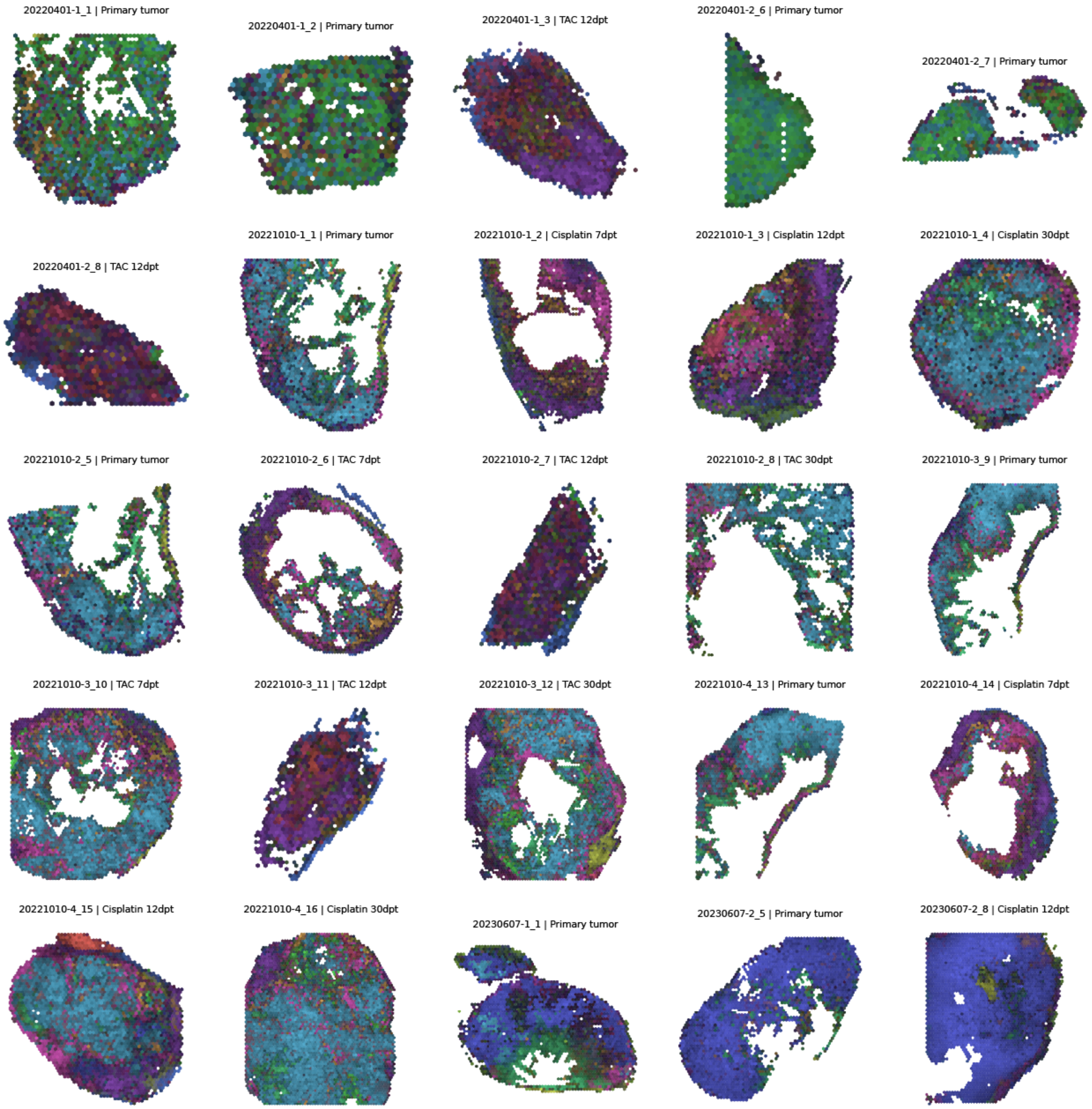


**Supplementary Fig. 6.** | **Cellular niche composition of tumours during MRD.** MIP of molecular tissue compartments (cellular niches) identified by Chrysalis across all samples in the main ST dataset.

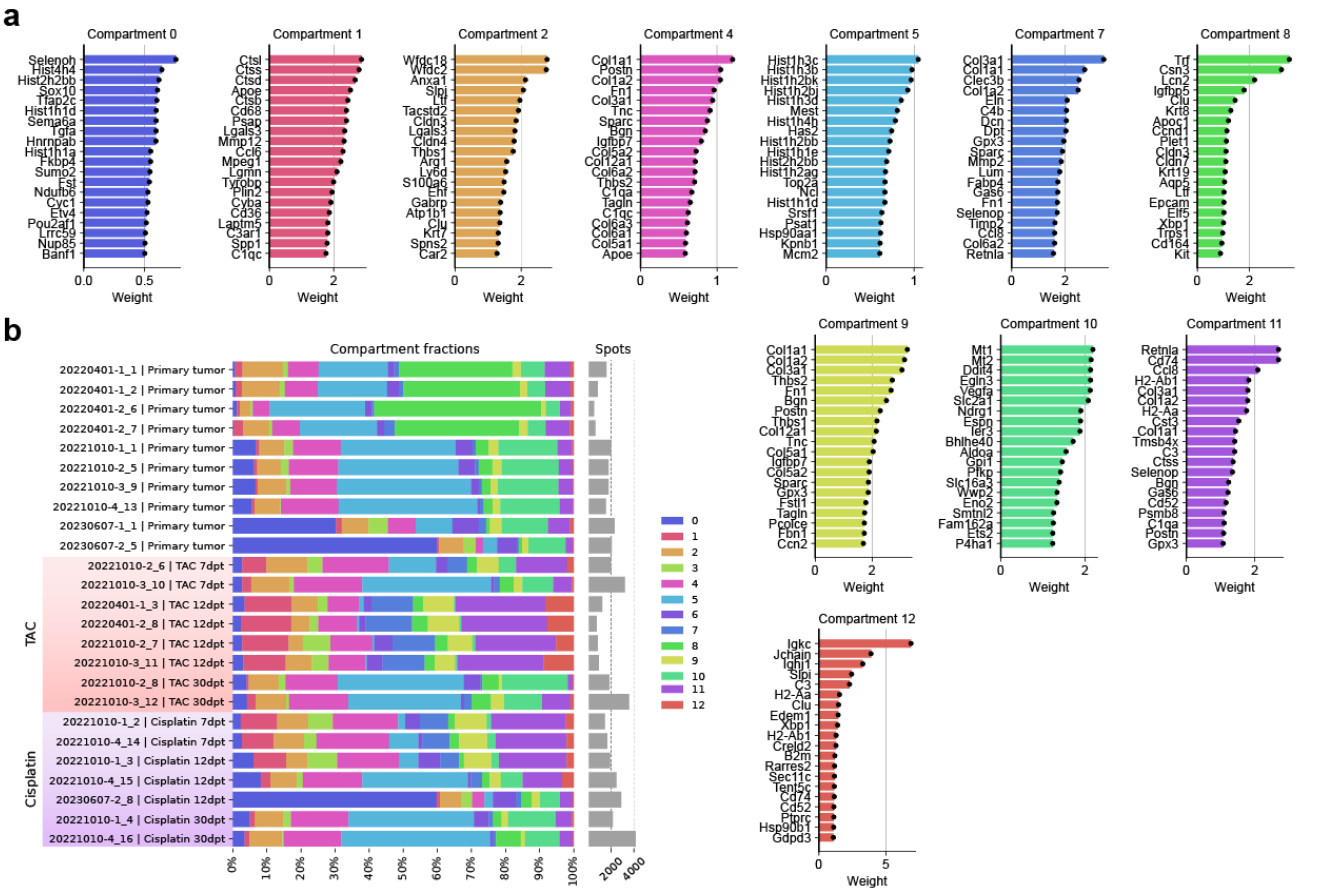


**Supplementary Fig. 7.** | **Cellular niches identified by Chrysalis. a,** Top 20 genes with the highest weights for each cellular niche identified in the main ST dataset. **b,** Cellular niche composition across all Visium tissue samples in the main ST dataset.

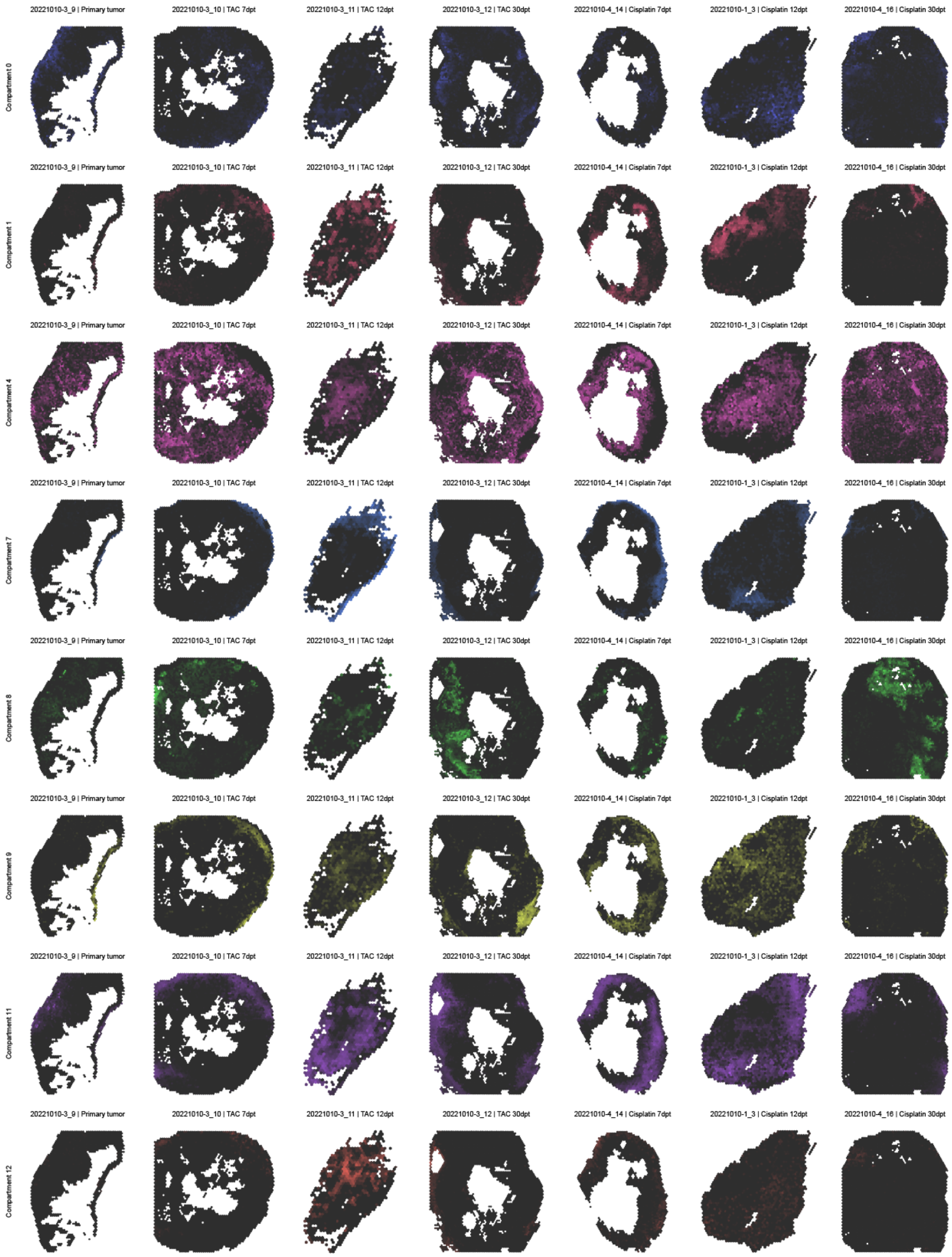


**Supplementary Fig. 8. | Spatiotemporal dynamics of cellular niches.** Spatial plots of compartment scores for cellular niches in representative tumour tissue sections from primary, residual, and recurrent tumours following TAC or cisplatin treatment (Compartments 2, 5, and 10 are shown in Fig.3d).

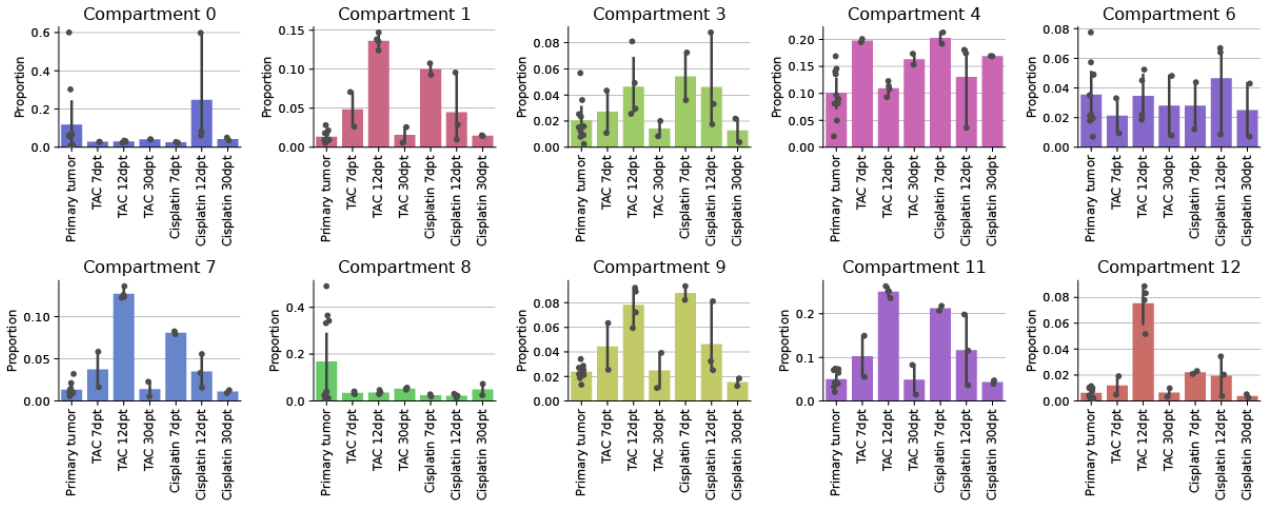


**Supplementary Fig. 9.** | **Overview of the temporal dynamics in cellular niches.** Proportions of cellular niches across all tumour samples (Compartments 2, 5, and 10 are shown in Fig.3e, bar height: mean compartment fraction, black dots: individual ST samples, error bar: 95% CI, bar height: mean value).

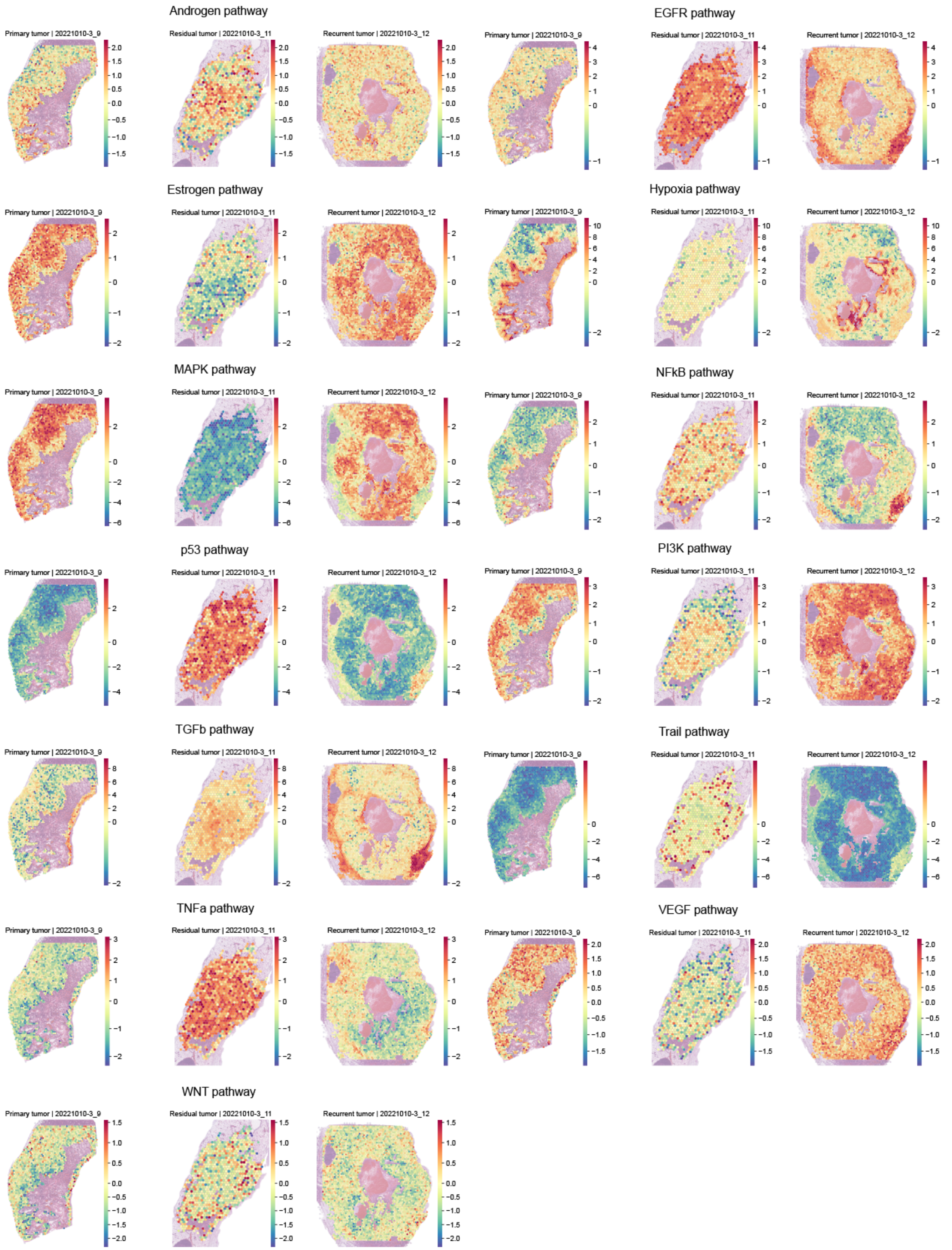


**Supplementary Fig. 10.** | **Spatial pathway activity maps inferred with PROGENy.** Spatial plots of pathway activities inferred using PROGENy, displayed for representative samples.

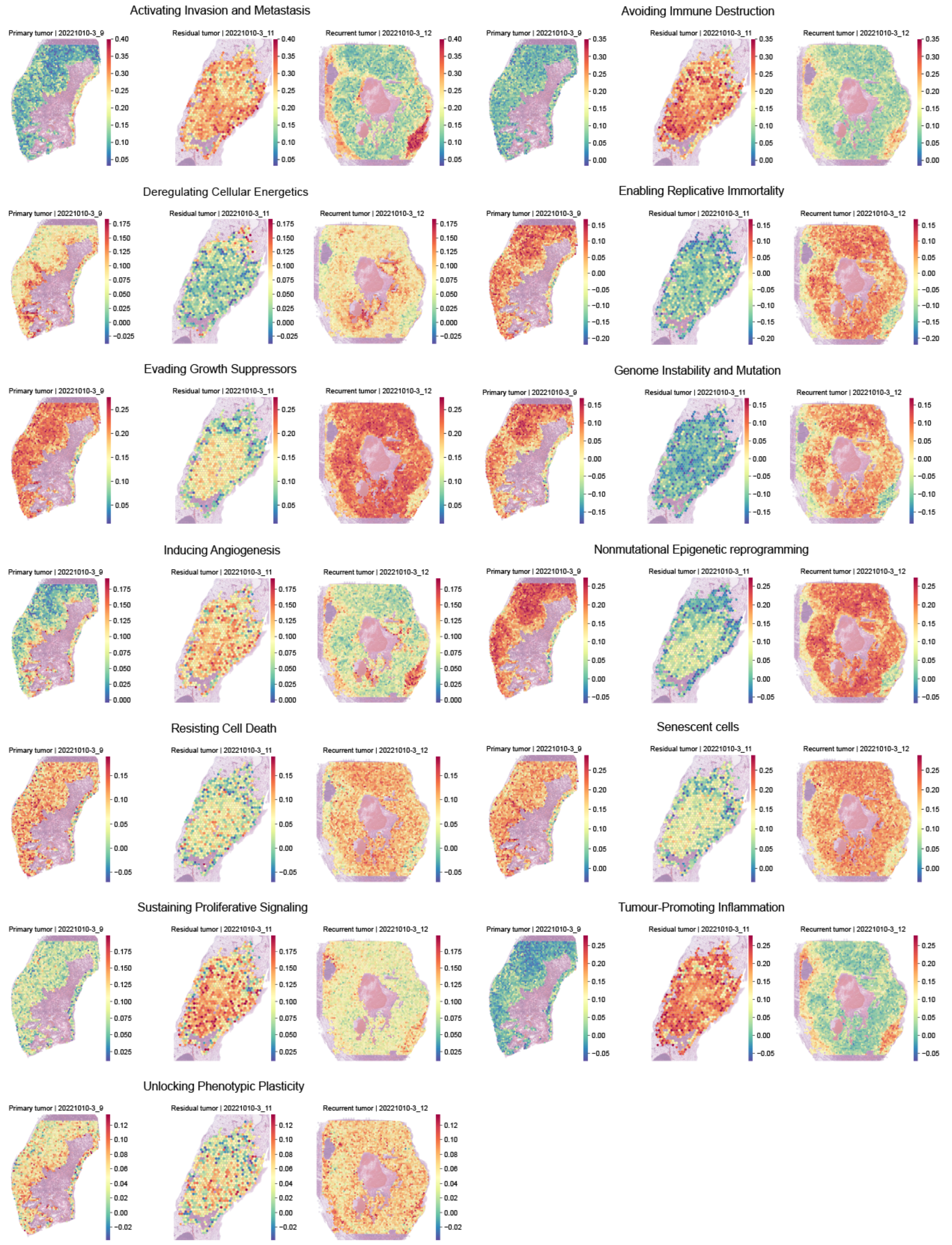


**Supplementary Fig. 11.** | **Spatial Hallmarks of Cancer enrichment scores.** Spatial plots of Hallmarks of Cancer enrichment scores shown for representative samples.

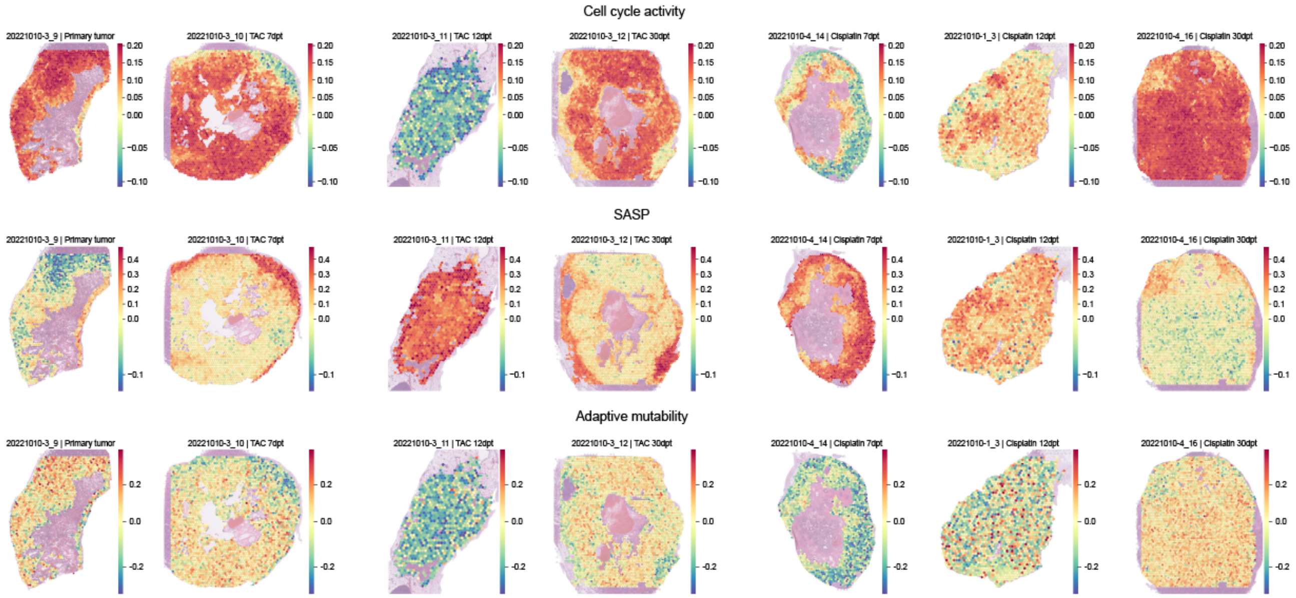


**Supplementary Fig. 12.** | **Cell cycle, SASP, and Adaptive mutability gene set activity.** Spatial plots of Cell cycle activity, SASP, and Adaptive mutability gene set scores shown for representative samples.

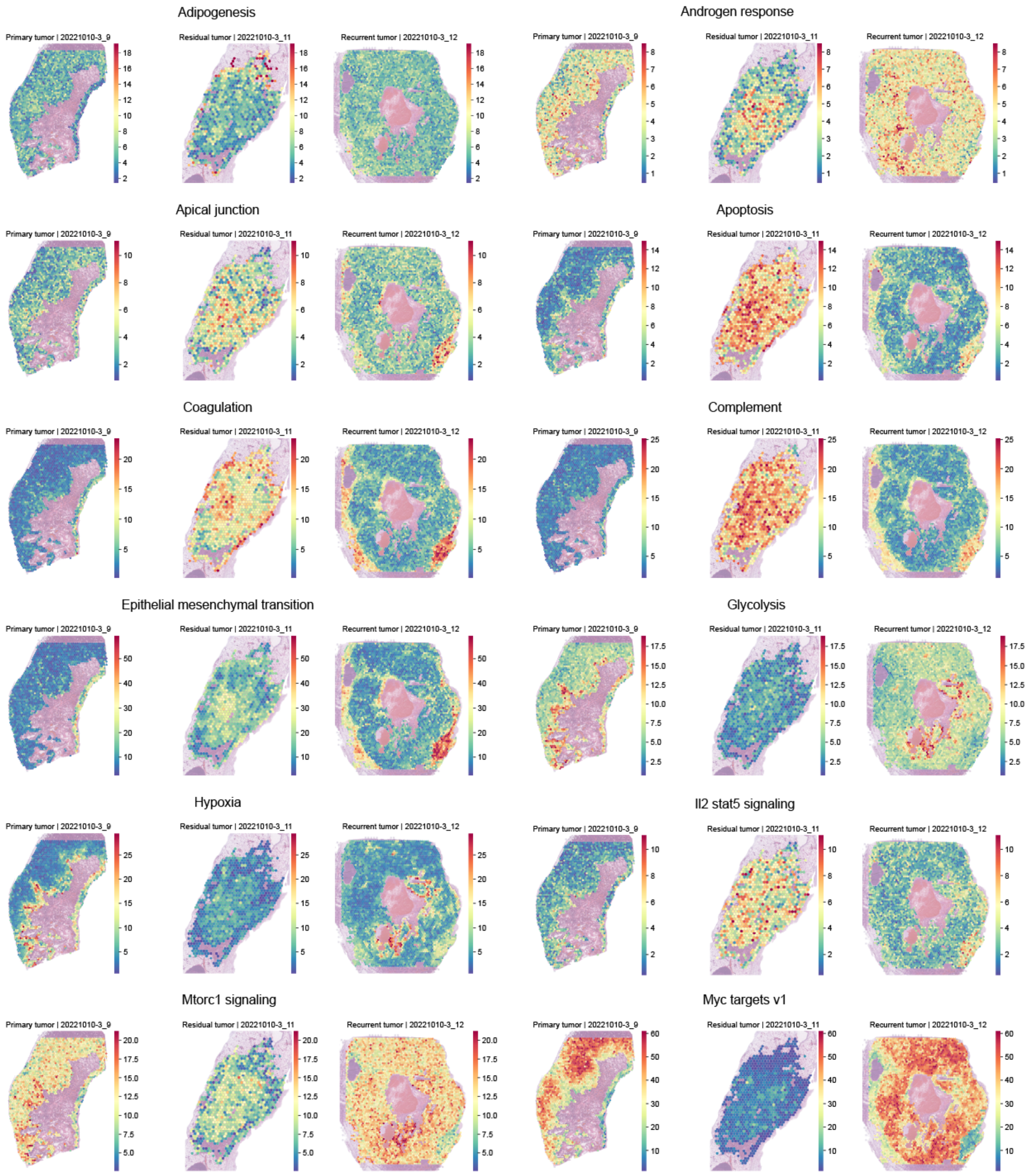


**Supplementary Fig. 13.** | **Spatial activity of Hallmarks gene sets 1/2.** Spatial plots of significantly enriched Hallmarks gene sets shown for representative samples.

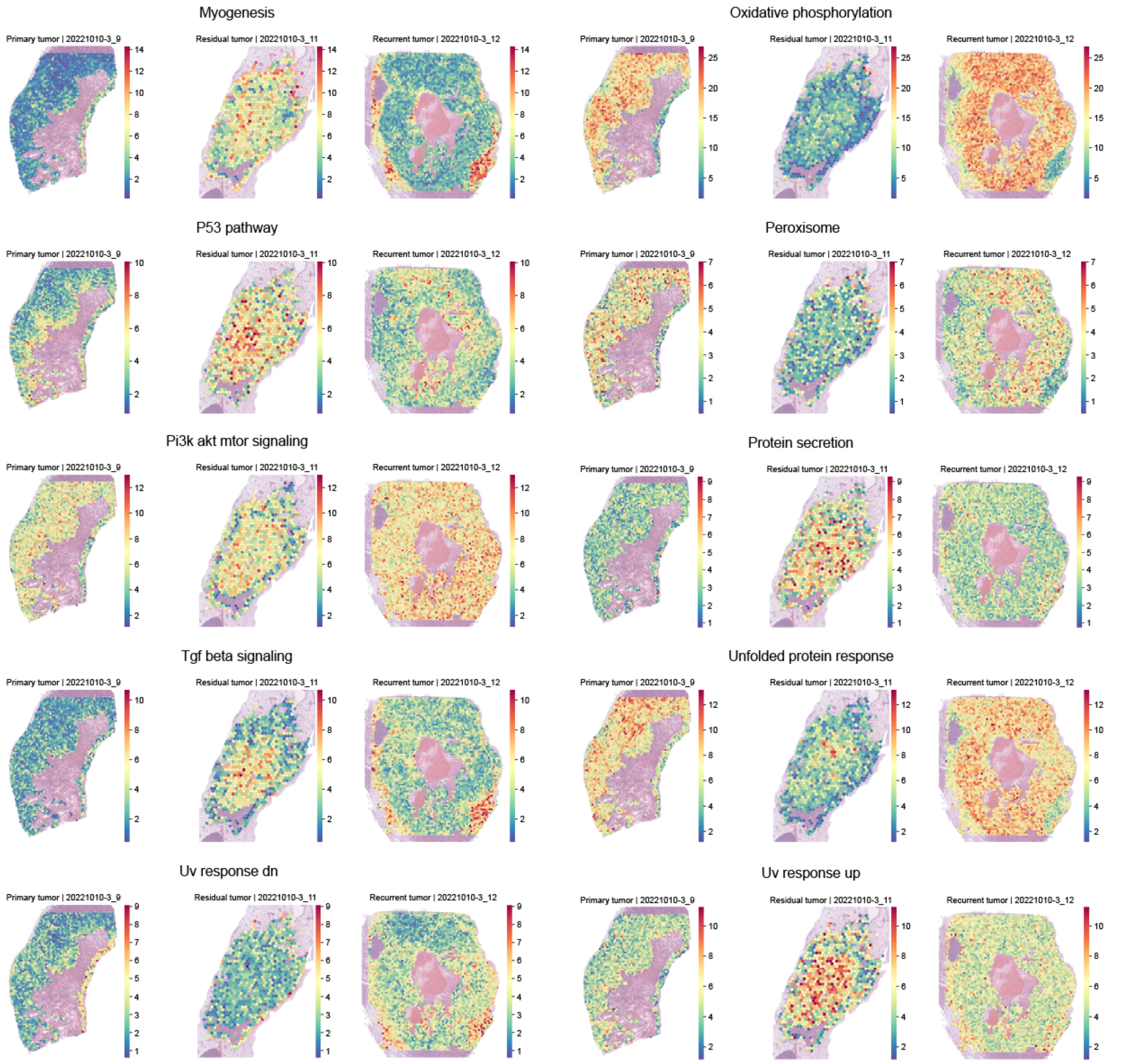


**Supplementary Fig. 14.** | **Spatial activity of Hallmarks gene sets 2/2.** Spatial plots of significantly enriched Hallmarks gene sets shown for representative samples.

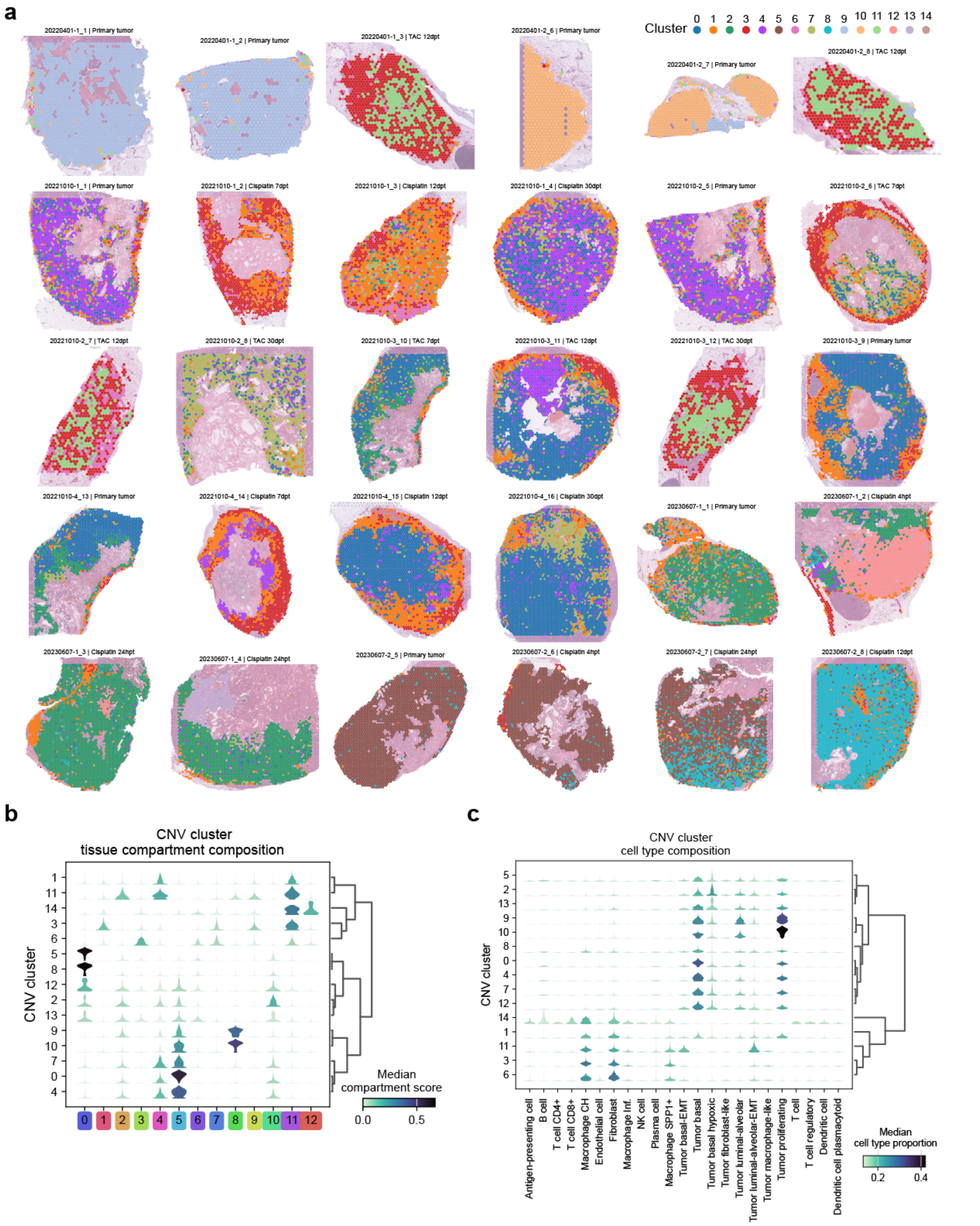

**Supplementary Fig. 15. | CNV clusters and their cellular composition. a,** Spatial distribution of CNV clusters across all Visium samples, showing that cluster distribution is more closely tied to their respective parental tumours than treatment conditions. **b,** Stacked violin plots depicting the cellular niche composition of each CNV cluster. **c,** Stacked violin plots showing the inferred cell type abundance composition of each CNV cluster.

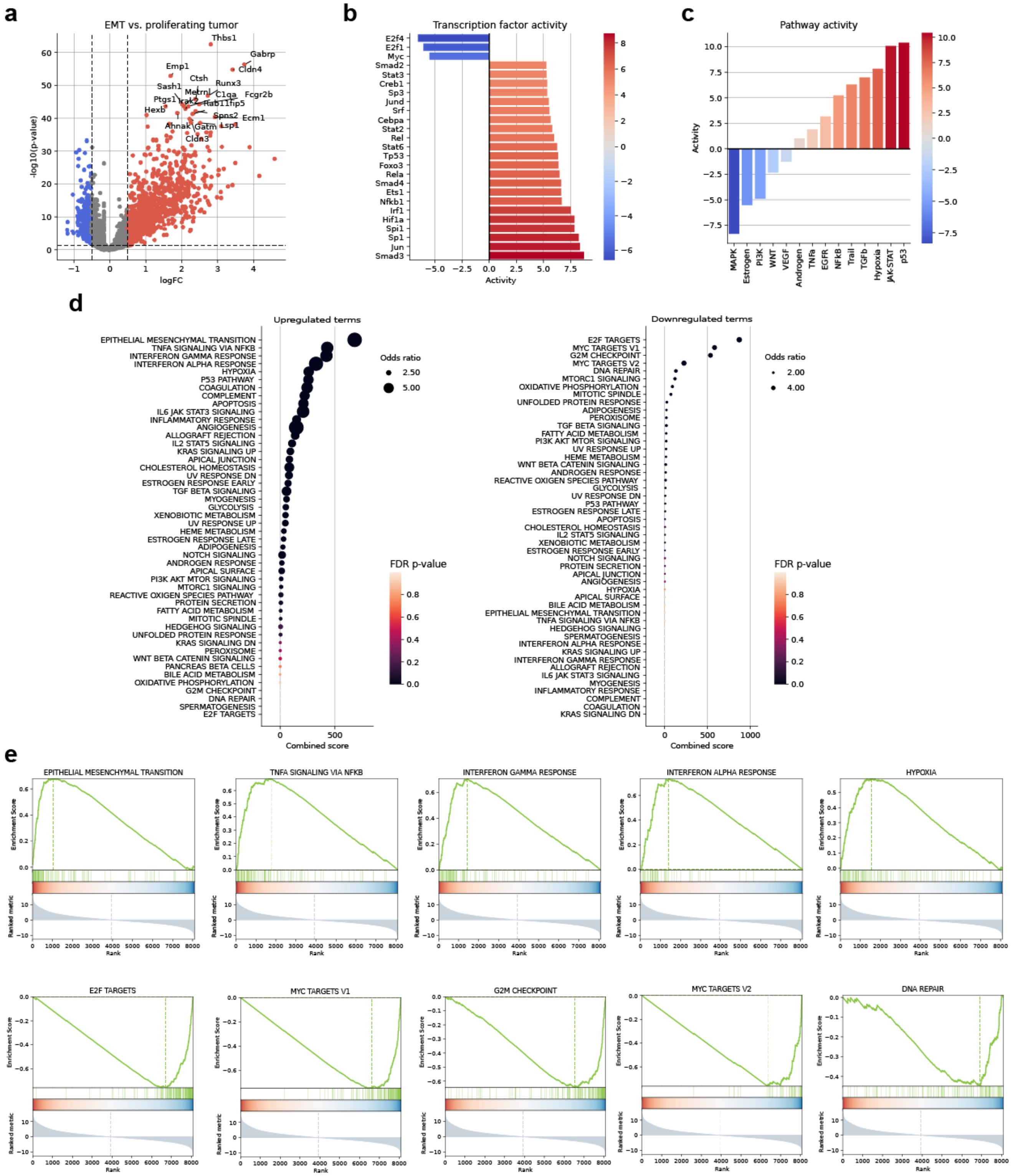

**Supplementary Fig. 16.** | **DGEA of the EMT and proliferating tumour niches. a,** Pseudo-bulk DGEA results for capture spots with dominant EMT and proliferating niche scores. **b,** Transcription factor activity inferred from the DGEA results. **c,** Pathway activity inference based on differentially expressed genes. **d,** Dot plots showing upregulated and downregulated terms from the Hallmarks gene set collection. **e,** Enrichment plots of the top five upregulated and downregulated gene sets from the Hallmarks gene set collection.

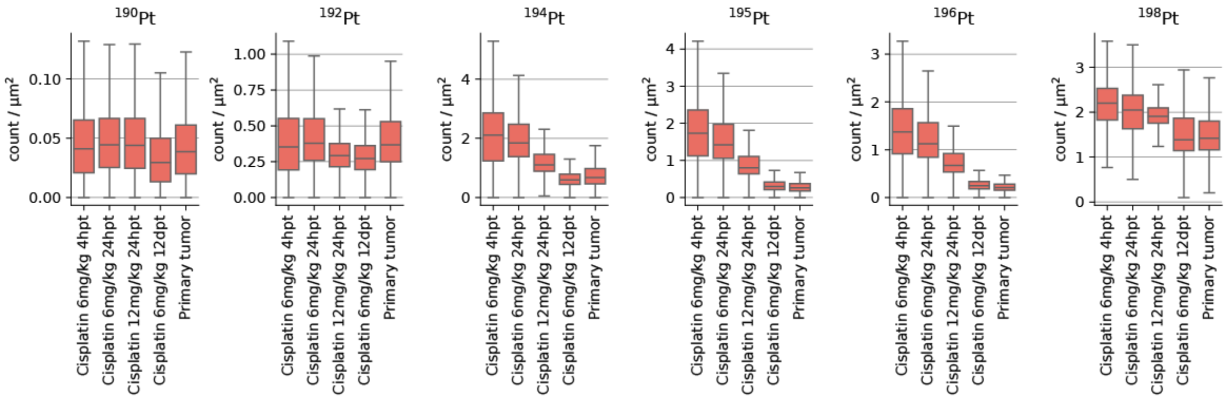


**Supplementary Fig. 17.** | **Distribution of intracellular concentration of Pt isotopes.** Bar plots showing the distribution of intracellular concentrations for the six observationally stable Pt isotopes measured with IMC across different conditions.

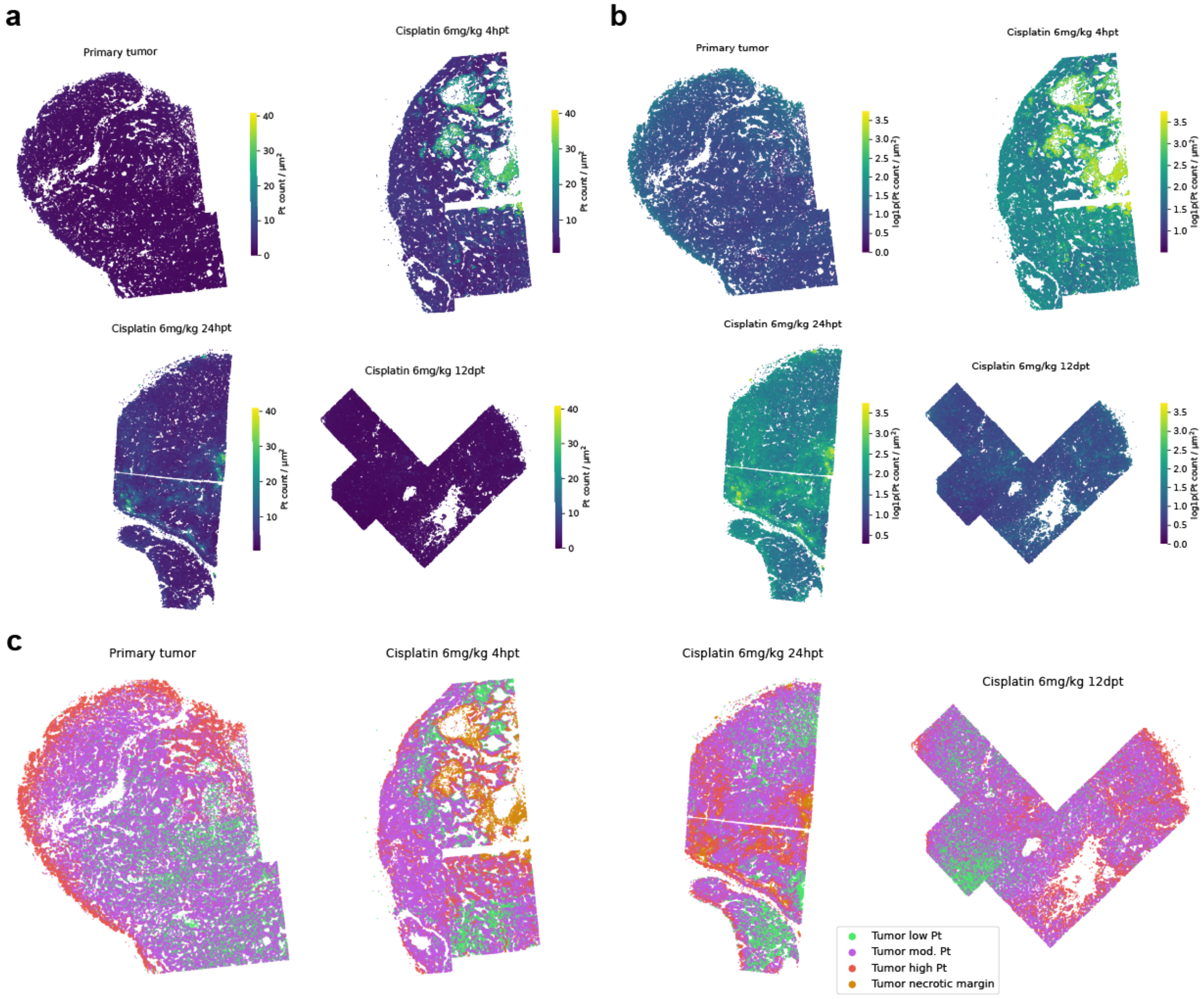


**Supplementary Fig. 18.** | **Intracellular Pt concentration in tumour cells.** **a,** Raw intracellular Pt concentration in individual tumour cells within tissue sections measured with IMC across the four conditions. **b,** Log-transformed intracellular Pt concentration in individual tumour cells within tissue sections measured with IMC across the four conditions. **c,** Classification of tumour cells into four groups based on the distribution of intracellular Pt content across the four conditions.

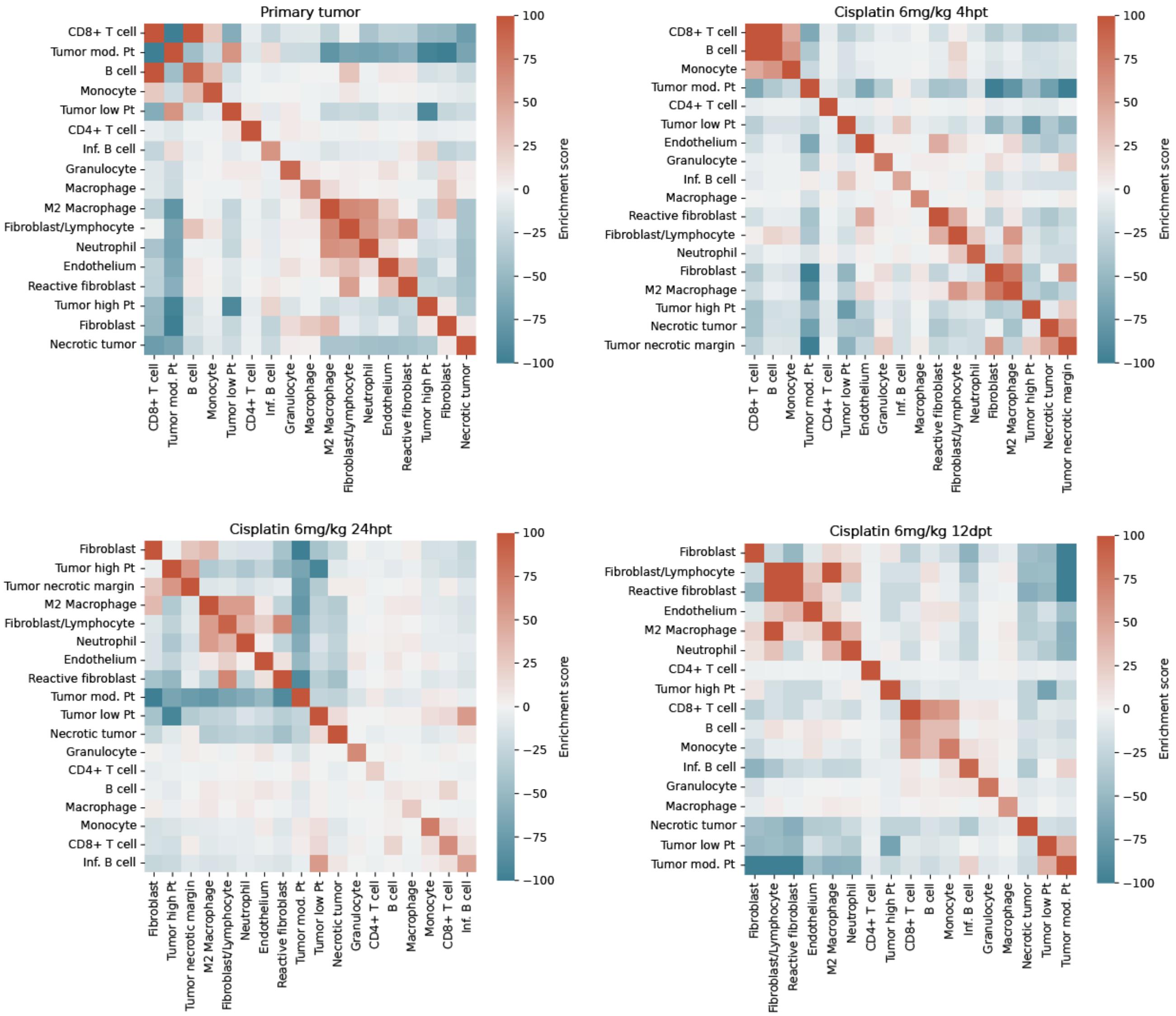


**Supplementary Fig. 19. | Neighborhood enrichment scores of cell types in IMC data.** Neighbourhood enrichment scores of cell types identified in tissue sections reconstructed with IMC across four different conditions.

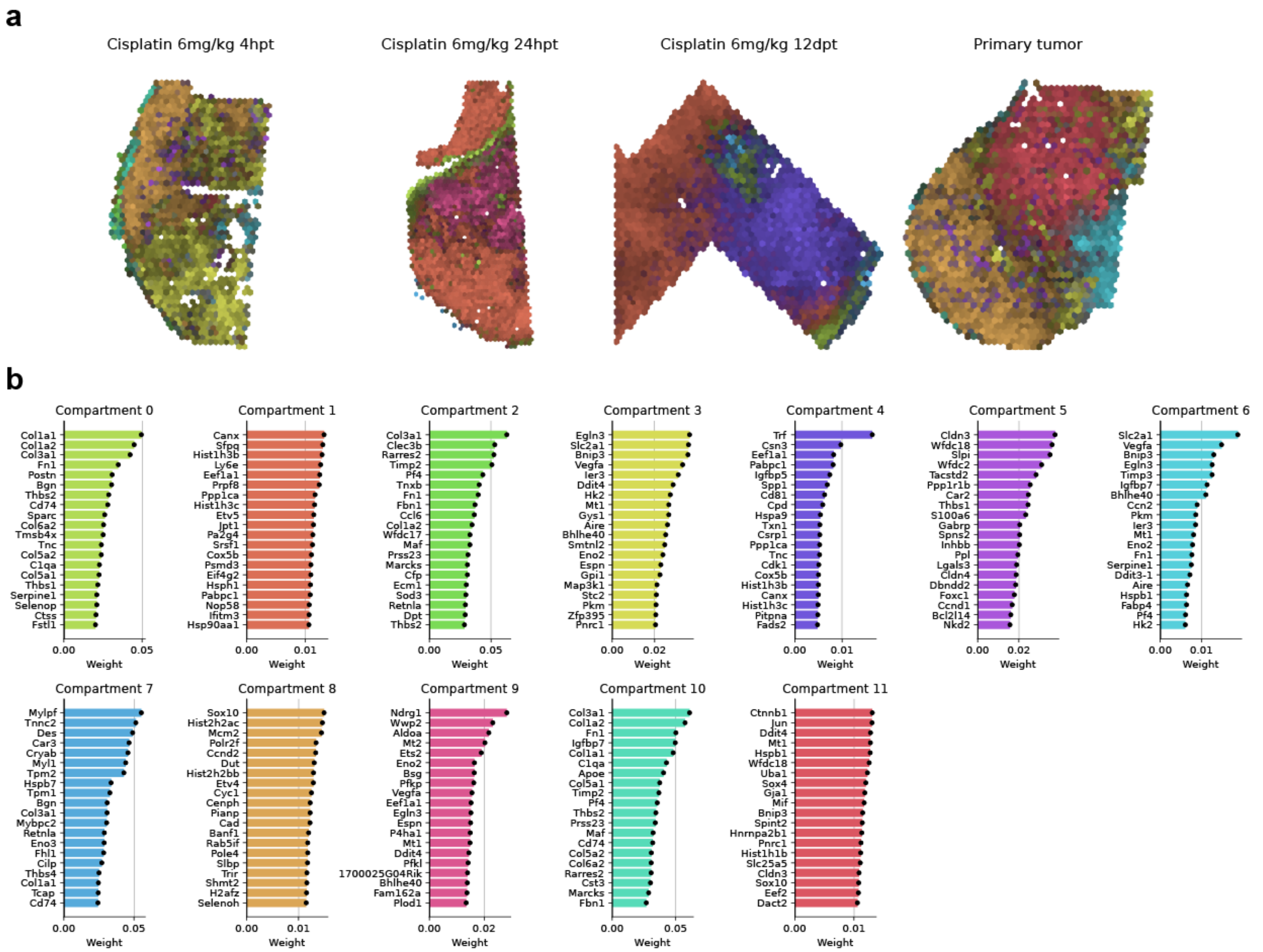


**Supplementary Fig. 20.** | **Cellular niches identified by Chrysalis in the multimodal ST-IMC dataset. a,** MIP of molecular tissue compartments (cellular niches) identified by Chrysalis across all samples in the Visium section of the multimodal ST-IMC dataset. **b,** Top 20 genes with the highest weights for each cellular niche identified in the Visium section of the multimodal ST-IMC dataset.

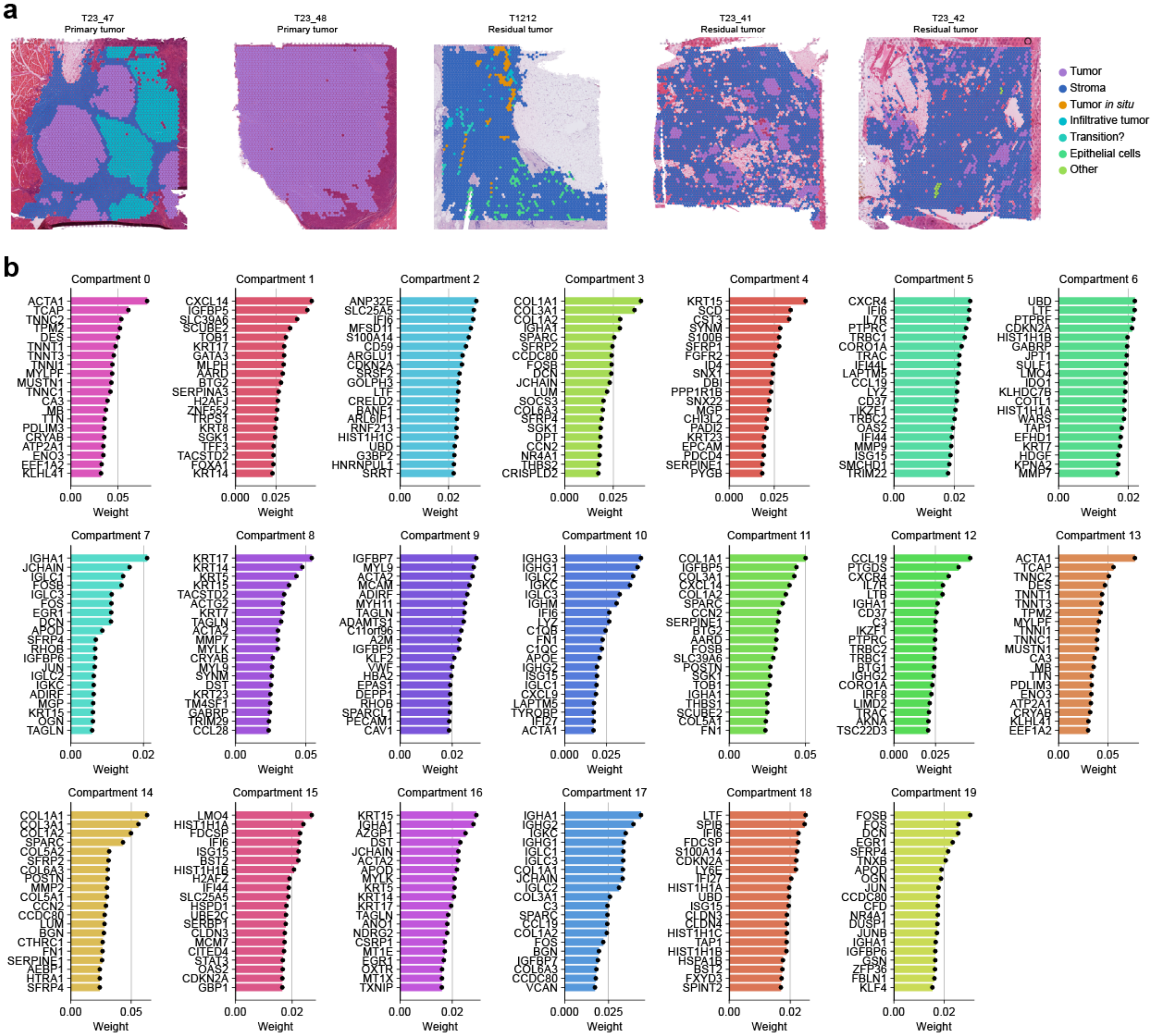


**Supplementary Fig. 21.** | **Histopathological annotations and gene weights in the human ST dataset. a,** Labeling of capture spots based on histopathological annotations. In primary tumours, tumour tissue was annotated based on invasiveness: Tumour (typical epithelial pattern), Infiltrative tumour (cells lose cell-cell contact and intermingle with immune cells and stroma). In residual tumours, epithelial tissue was annotated based on morphology: Epithelial cells (resembling healthy ducts with no signs of malignancy), Tumour *in situ* (visually malignant but well demarcated), Infiltrative tumour (visually malignant and invading surrounding tissue), Transition (cells transitioning from normal to malignant epithelial cells), Other (undefined tissue). **b,** Top 20 genes with the highest weights for each cellular niche identified in the human ST dataset.

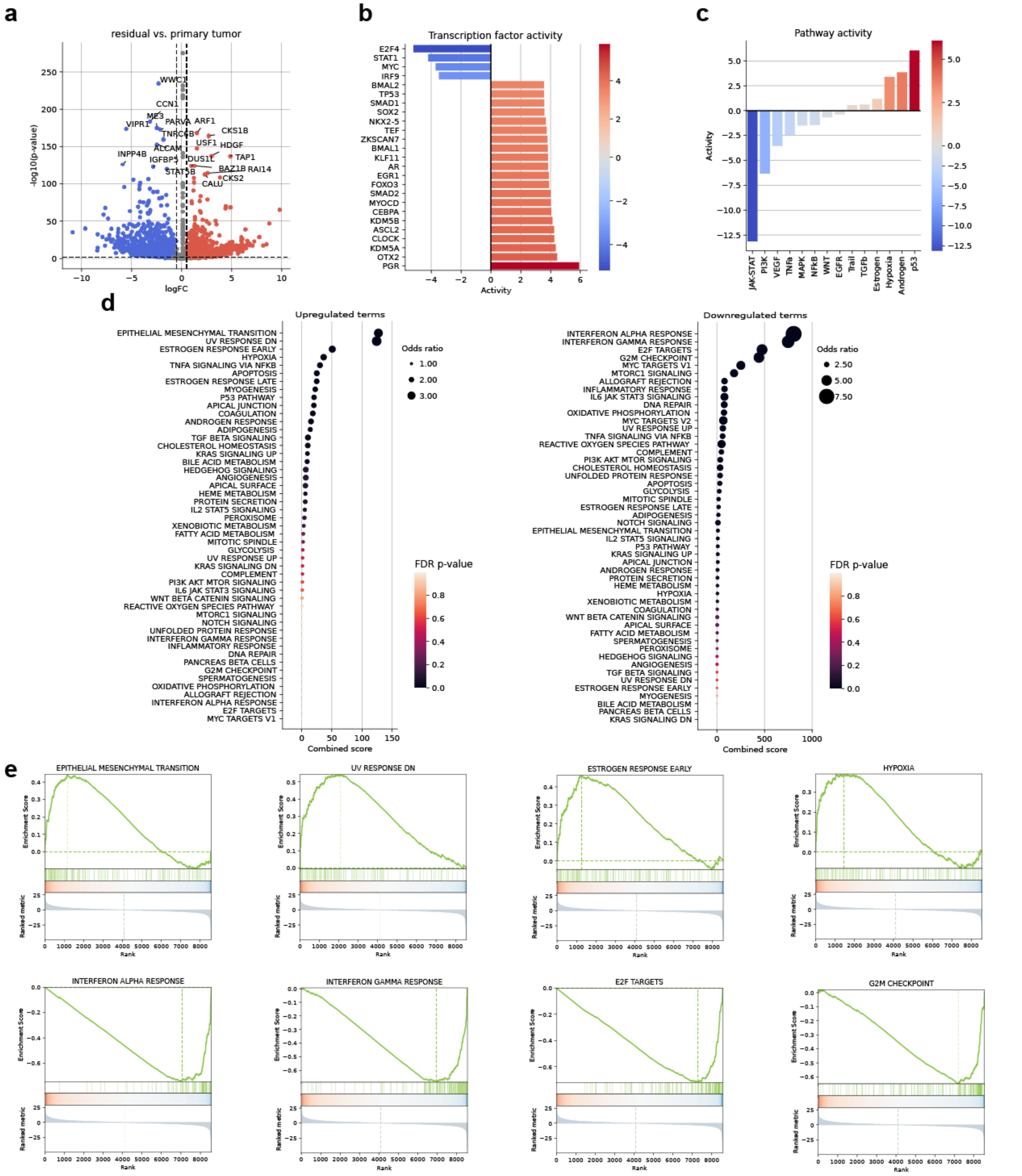

**Supplementary Fig. 22.** | **DGEA of the residual and proliferating tumour niches in humans. a,** Pseudo-bulk DGEA results for capture spots with dominant EMT and proliferating niche scores. **b,** Transcription factor activity inferred from the DGEA results. **c,** Pathway activity inference based on differentially expressed genes. **d,** Dot plots showing upregulated and downregulated terms from the Hallmarks gene set collection. **e,** Enrichment plots of the top five upregulated and downregulated gene sets from the Hallmarks gene set collection.

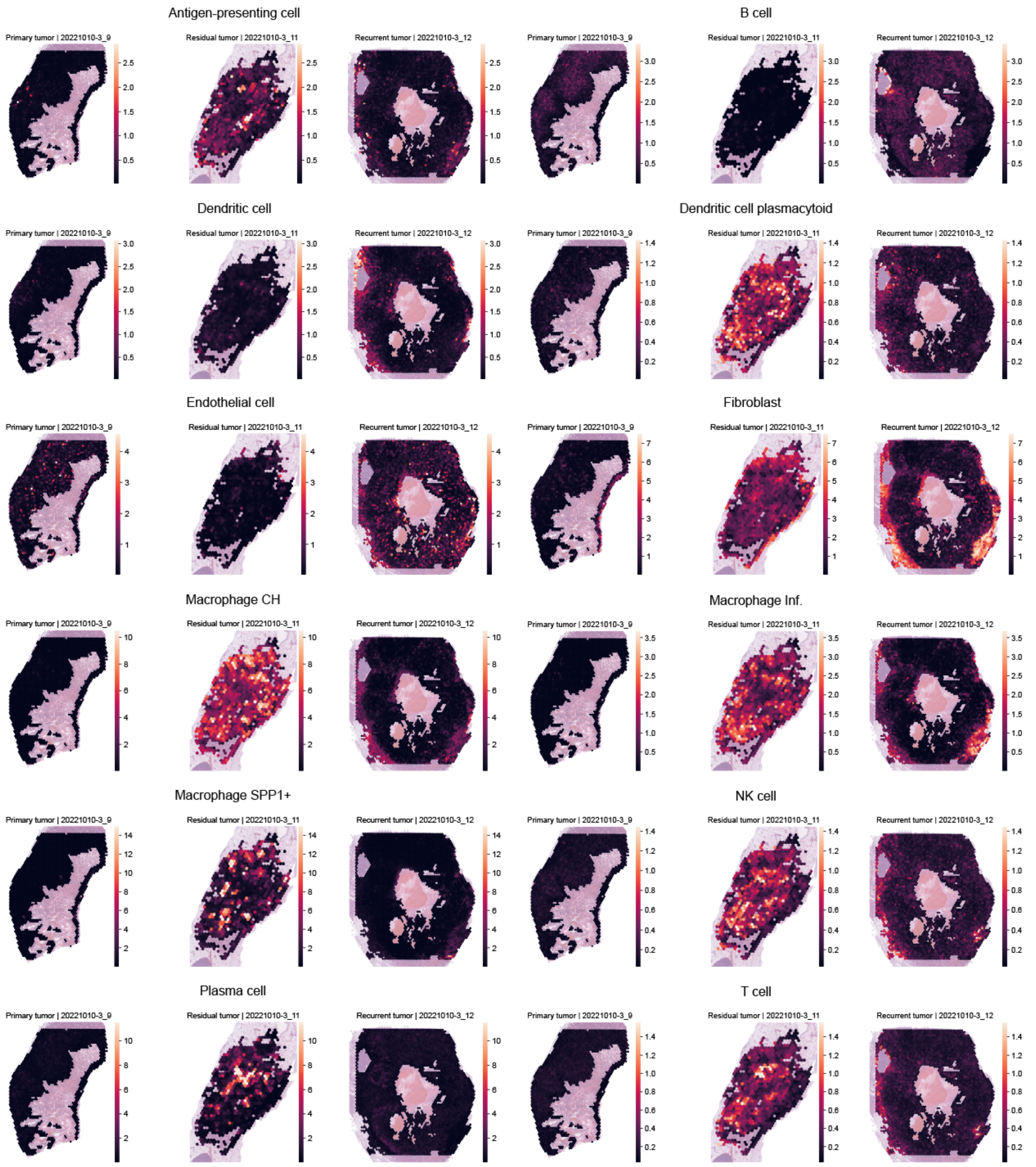


**Supplementary Fig. 23.** | **Cell type abundances inferred with cell2location 1/2.** Spatial plots of cell type abundances inferred using cell2location, shown for representative samples.

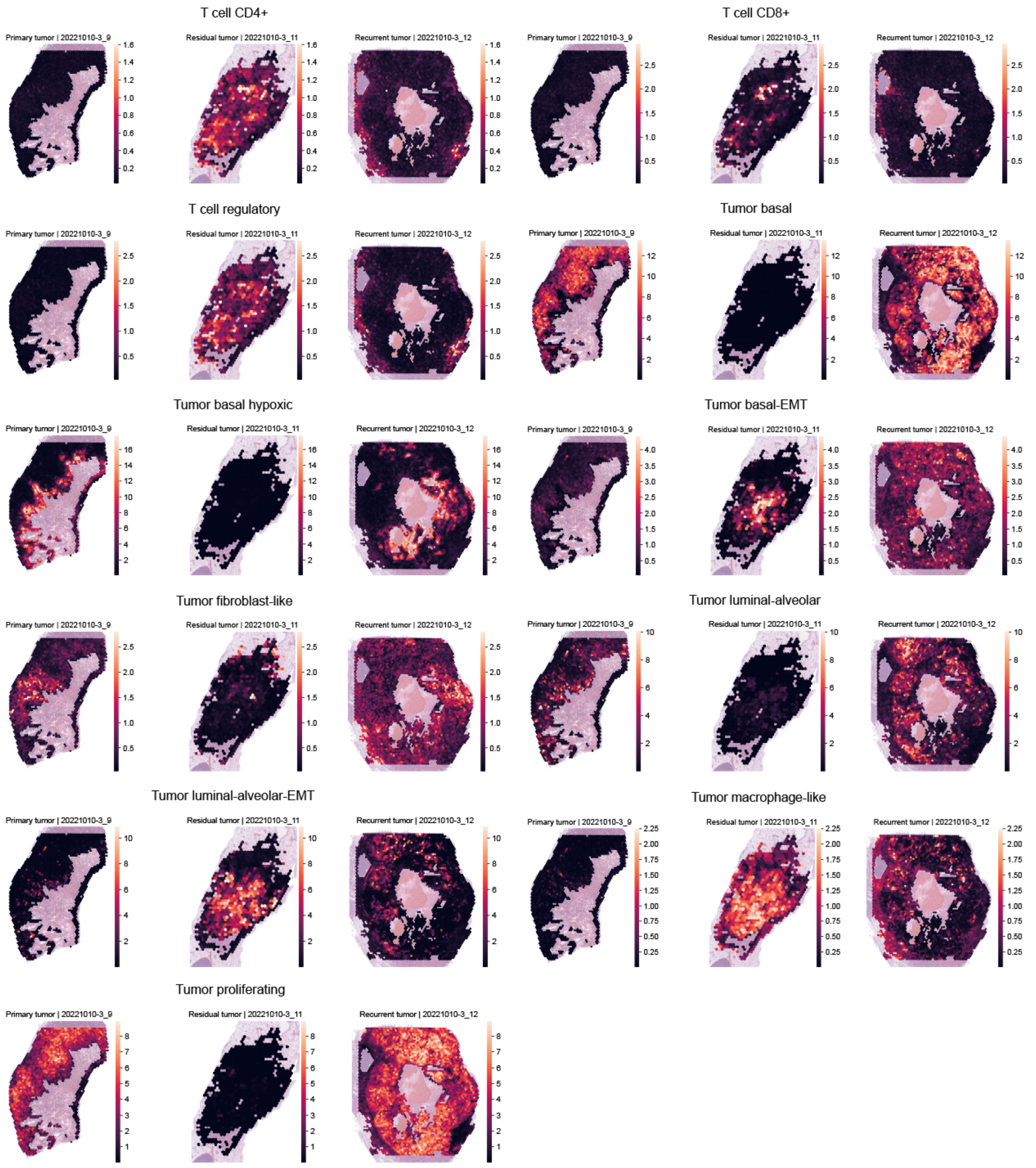


**Supplementary Fig. 24.** | **Cell type abundances inferred with cell2location 2/2.** Spatial plots of cell type abundances inferred using cell2location, shown for representative samples.

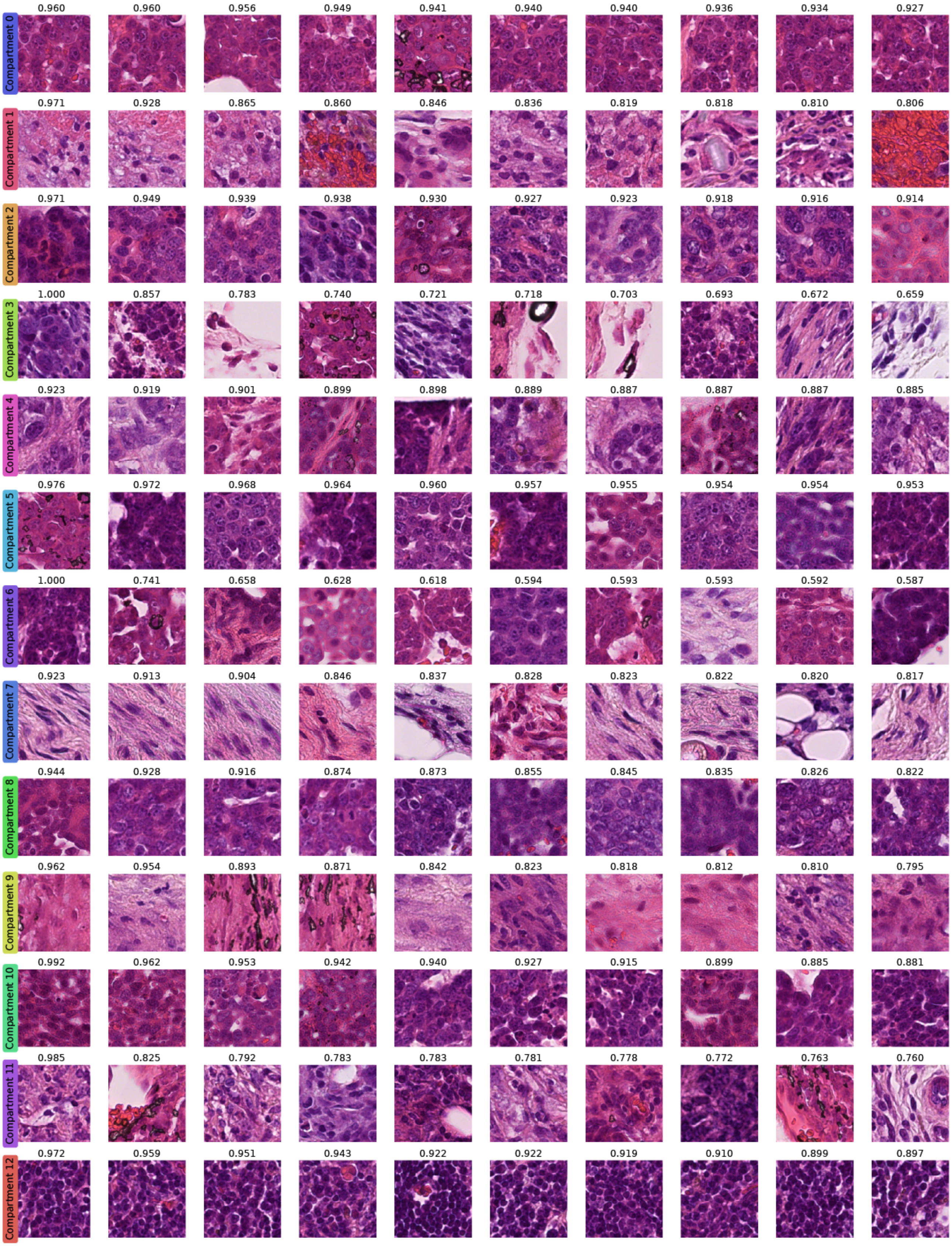


**Supplementary Fig. 25.** | **Image tiles of highest-scoring capture spots for cellular niches.** Image tiles (55 × 55 µm) of the 10 highest scoring capture spots for each cellular niche, with compartment scores inferred by Chrysalis, in the main ST dataset.
